# Temporal dissociation between local and global functional adaptations of the maternal brain to childbirth: A longitudinal assessment

**DOI:** 10.1101/2023.08.15.553345

**Authors:** Leon D. Lotter, Susanne Nehls, Elena Losse, Juergen Dukart, Natalia Chechko

## Abstract

The maternal brain undergoes significant reorganization during birth and the postpartum period. However, the temporal dynamics of these changes remain unclear. Using resting-state functional magnetic resonance imaging, we report on local and global brain function alterations in 75 mothers in their first postpartum week, compared to 23 nulliparous women. In a subsample followed longitudinally for the next six months, we observed a temporal and spatial dissociation between changes observed at baseline (cluster mass permutation: pFWE < .05). Local activity and connectivity changes in widespread neocortical regions persisted throughout the studied time period (ANCOVAs vs. controls: pFDR < .05), with preliminary evidence linking these alterations to behavioral and psychological adaptations (interaction effect with postpartum time: uncorrected p < .05). In contrast, the initially reduced whole-brain connectivity of putamen-centered subcortical areas returned to control levels within six to nine weeks postpartum (linear and quadratic mixed linear models: pFDR < .05). The whole-brain spatial colocalization with hormone receptor distributions (Spearman correlations: pFDR < .05) and preliminary blood hormone associations (interaction effect with postpartum time: uncorrected p < .05) suggested that the postpartum restoration of progesterone levels may underlie this rapid normalization. These observations enhance our understanding of healthy maternal brain function, contributing to the identification of potential markers for pathological postpartum adaptation processes, which in turn could underlie postpartum psychiatric disorders.

## 1. Introduction

The maternal brain experiences major biological and psychological changes during pregnancy and the early postpartum period, suggesting adult brain neuroplasticity. Widespread changes in gray matter volume (GMV) and cortical thickness distinguish maternal brains from those of nulliparous women^1^. These morphometric changes occur most dynamically within the first 6 weeks postpartum^2^, likely mediated by sex steroids ^3–6^. However, neocortical brain structure alterations may persist beyond 12 weeks postpartum^2,7–10^, suggesting region-specific short-term and long-term effects of pregnancy, childbirth, and motherhood on the brain^11^. Although the biological role of these changes is not fully understood, they may be relevant for maternal attachment toward the offspring^2,7,12^.

Given these morphological changes, along with (i) the hormonal adaptations in the perinatal period^3,13^, (ii) the reported hormonal regulation of brain connectivity^5,14,15^, and (iii) the interactions between sex hormones and GABAergic as well as oxytonergic neurotransmission^16,17^, it may be safe to assume that brain function undergoes similar reorganization. Resting-state functional magnetic resonance imaging (rsfMRI) provides an effective, scalable approach to explore these changes independent of participant compliance or performance^18^. A prior rsfMRI study^19^ reported alterations of the default mode network (DMN) functional connectivity (FC) in postpartum mothers (one session between 1 and 4 months postpartum) compared to women pre-conception. However, high-resolution follow-up data are needed to elucidate the brain’s functional reorganization during the early postpartum period, differentiating short-term biological impacts of pregnancy and early post-partum from those of motherhood^2^. Voxel- or cluster-level MRI inferences are limited in terms of their ability to help identify the biological mechanisms underlying postpartum rsfMRI adaptations. Spatial colocalization analyses, which quantify the alignment between observed MRI maps and maps of a biological process or entity of interest^20–22^, can offer more detailed insights. This approach has been successfully used to study individual-level structural brain development^21^ as well as rsfMRI alterations in Parkinson’s and Huntington’s diseases^22,23^.

We assessed healthy maternal brain function within the first postpartum week in a large baseline sample (n=75) and in a subgroup followed longitudinally at 3, 6, 9, 12, and 24 weeks (6 months) postpartum. Alongside MRI, we measured serum estradiol and progesterone levels and behavioral data at each session. We expected that, compared to healthy nulliparous controls (NP), postpartum women (PP) would show strong alterations of rsfMRI metrics mapping intra- and interregional brain activity and FC. Our study design afforded key insights into the transitory and persistent natures of postpartum rsfMRI changes, allowing us to test for potential interactions with hormonal adaptations. To that end, we tested (i) if whole-brain distributions of rsfMRI activity and connectivity changes colocalized spatially with hormone receptors and functionally-related neurotransmitter receptors, and (ii) if rsfMRI changes covaried with serum hormone levels over time. Additionally, we sought to determine if the enduring changes in brain function related to psychological or behavioral shifts^2,7,8,12,24^, particularly with respect to mother-infant attachment and maternal depression.

## 2. Methods

### 2.1. Study samples

Postpartum women were recruited for two studies run by the Department for Psychiatry, Psychotherapy, and Psychosomatics, University Hospital Aachen, Germany (Supplement 1.1). The baseline sample (T0, Study 1) consisted of 75 non-depressed postpartum women recruited within 7 days of delivery. From this sample, 19 postpartum women took part in the longitudinal Study 2, with MRI measurements at 3, 6, 9, 12, and 24 weeks postpartum (T1–T5). At each time point, blood samples were drawn to determine estradiol and progesterone levels (Supplement 1.2). Post-partum women were assessed for postpartum depression symptoms through the Edinburgh Postnatal Depression Scale (EPDS; T0–T5; total score)^25^ and for mother-child attachment through the Maternal Postnatal Attachment Scale (MPAS; T1–T5; total score and sub-scores for quality of attachment, absence of hostility, and pleasure in interaction)^26^. The control group consisted of 23 healthy nulliparous women with no history of psychiatric disorders. Prior to enrolment, written informed consent was obtained from each participant. After the exclusion of sessions that had exceeded motion cut-offs and had not passed visual quality control (n=1 with MRI data reconstruction failure), the final dataset consisted of data from 98 subjects (PP=75, NP=23) with overall 189 sessions (longitudinal: T0=19, T1=19, T2=18, T3=19, T4=18, T5=17).

The study protocol was in accordance with the Declaration of Helsinki and was approved by the Institutional Review Board of the Medical Faculty, RWTH Aachen University, Germany.

### 2.2. Software and code availability

The software and toolboxes used^27–36^ are listed in Supplement 1.3. The analysis code, organized in a Jupyter notebook, is available as a supplement.

### 2.3. MRI acquisition and preprocessing

MRI acquisition parameters and general processing are described in Supplement 1.4. RsfMRI-specific processing included (i) regression of “Friston-24” motion parameters^37^, mean white matter, and cerebrospinal fluid signals, (ii) linear detrending, and (iii) bandpass-filtering (.01–.08 Hz). For spatial colocalization analyses, data were parcellated into 100 cortical and 16 subcortical parcels^38,39^. Subjects with excessive motion were excluded.

### 2.4. Analysis of demographic, behavioral, hormonal, and MRI motion data

We tested for differences in demographic, behavioral, clinical, and MRI motion characteristics between the postpartum and nulliparous groups as well as between the cross-sectional and longitudinal postpartum groups using Mann-Whitney-U and Chi^2^ tests. Longitudinal trajectories of hormone levels, psychological variables, and fMRI-derived measures were evaluated using linear mixed models (LMM). We assessed the fixed effect of continuous postpartum week on each tested response variable, including random intercepts for each subject. For each response variable, two mixed linear models were calculated: (i) including only the linear postpartum week as fixed effect and (ii) including linear and quadratic effects.

### 2.5. RsfMRI analyses at baseline

To identify brain areas showing robust adaptation effects of pregnancy and childbirth, we followed a comprehensive rsfMRI analysis approach^40^, simultaneously assessing postpartum changes in voxel-level *intra-*regional/local rsfMRI activity (fractional amplitude of low frequency fluctuations, fALFF)^41^, *intra-*regional/local FC (Local Correlation, LCOR), as well as *inter-*regional/global FC (Global Correlation, GCOR)^42^. For descriptions of these metrics, please refer to Lotter et al.^40^. To determine rsfMRI alterations occurring in the first postpartum week, we compared baseline postpartum and nulliparous groups in general linear models (GLM) controlling for age. Cluster-level significance was determined in separate positive (PP>NP) and negative contrasts (PP<NP) based on permutation of cluster masses (10,000 iterations) using a voxel-level threshold of p<.01 and a cluster-level threshold of p<.05. Cluster-permutation approaches have been shown to accurately control the family-wise error (FWE) rate, while avoiding false negatives due to excessive multiple comparison correction on the voxel-level^43^. Permutation of cluster *masses* (sum of the test statistic across voxels in a cluster) can be more powerful than permutation of cluster *sizes* (number of voxels)^44^.

For further analyses, average rsfMRI data for each significant cluster was extracted for all subjects and sessions. For comparisons with future studies, and as the cross-sectional and longitudinal postpartum samples differed in some demographic, clinical, and MRI motion variables, we tested for associations between these variables and the cluster-level MRI data using ANCOVAs and linear regressions controlled for age.

### 2.6. Longitudinal rsfMRI analyses

To assess how rsfMRI changes at baseline developed during the first 6 postpartum months, we tested the averaged cluster data for (i) differences between postpartum and nulliparous groups at each time point using ANCOVAs controlled for age, (ii) longitudinal development in postpartum women using LMMs, and (iii) within-subject differences between time points in multiple paired t-tests. LMMs were designed to assess the fixed effect of postpartum week on each MRI metric, while accounting for the within-subject variance. In sensitivity analyses, we (i) included average gray matter signals within the same clusters, extracted from modulated gray matter maps, as co-variates^40^ and (ii) evaluated the possibility of regression-toward-the-mean effects driving longitudinal normalization patterns of rsfMRI alterations (Supplement 1.5). As for the composition of our study samples, we focused on the temporal development of rsfMRI clusters identified at baseline relative to controls, rather than on longitudinal GLMs estimated in only the comparably small longitudinal postpartum subgroup. Nevertheless, we sought to determine whether whole-brain patterns in the longitudinal postpartum data mirrored those observed in the cross-sectional postpartum-vs.-control data by spatially correlating beta maps from each rsfMRI metric and contrast (Supplement 1.6). During the same process, we also compared beta maps between different rsfMRI metrics to ascertain if the three metrics (fALFF, LCOR, GCOR) captured different biological processes.

### 2.7. Spatial colocalization between rsfMRI changes and receptor distributions

Assuming that the change of a brain function metric is influenced by a brain system on another neurobiological level, the spatial distribution of this metric across the brain may colocalize with the biological system in question^21–23^. We therefore examined if postpartum rsfMRI changes and their normalization patterns after childbirth were distributed across the brain following spatial distributions of pregnancy-related hormonal receptors (progesterone: PGR, estrogen: ESR1/ESR2, cortisol: NR3C1/NR3C2) and functionally close neurotransmitter receptors (oxytocin: OXTR, GABA: GABAA, glutamate: mGluR5). Maps of receptor distributions were obtained from Allen Brain Atlas postmortem gene expression^45^ (PGR, ESR1/ESR2, NR3C1/NR3C2, and OXTR) and in-vivo nuclear imaging data^46–48^ (GABAA and mGluR5; Supplement 1.7). We first assessed if the mean spatial colocalization between receptor distributions and baseline rsfMRI alterations in post-partum women relative to controls exceeded those of permutated data. Briefly, using the JuSpyce toolbox^30^, we calculated Z-scores for 116 whole-brain parcels for each postpartum woman at baseline [for each parcel: *“(PP – mean(NP)) / std(NP*)*”*] and correlated these with each receptor map across space using Spearman correlations^22^. Non-parametric exact p values [“p(exact)”] were derived from group permutation and, to accurately apply rank-based multiple comparison correction, final p values were calculated from Gaussian curves fitted to the null distributions [“p(norm)”]^49^. See Supplement 1.8 and Dukart et al.^22^ for the detailed workflow. RsfMRI-receptor combinations that showed significant colocalization at baseline were followed up in the longitudinal data using the same colocalization workflow and LMMs as described above. To provide a quantification of the extent to which postpartum rsfMRI alterations were explained by receptor distributions, we assessed the variance explained in multivariate linear regression analyses (R^2^) “predicting” baseline rsfMRI alterations from receptor maps^21,50^. Finally, to aid interpretation, we additionally evaluated the spatial colocalization between receptor maps (procedure as in Supplement 1.6).

We emphasize that spatial colocalization analyses have to be interpreted with care, especially when involving the Allen Brain Atlas gene expression data, which were obtained from only 6 postmortem brains (one 49-year-old female of Hispanic origin).

### 2.8. Analyses of MRI, hormone, and behavior associations

In order to identify possible physiological and behavioral correlates of the rsfMRI findings, we fitted LMMs to evaluate the effects of (i) interactions between hormone levels (log-transformed progesterone, estrogen, and progesterone/estrogen ratio) and postpartum week on MRI metrics (extracted cluster averages and FDR-significant spatial correlations) and (ii) the interaction between MRI metrics and postpartum week on EPDS and MPAS scores. We focused on the interaction effects to explore temporal covariance patterns between MRI and hormonal/behavioral data. To reduce between-subject variance and emphasize the longitudinal development patterns, data of each independent variable were z-standardized within subjects. For reference and visualization, we additionally calculated whole-brain beta maps for all interaction effects.

Given the small longitudinal sample size, these analyses must be considered preliminary, and their interpretation ought to be limited to evaluating patterns rather than specific associations. To allow for this, results were visualized as a heatmap focusing on effect sizes. We also highlight the associative nature of the analyses, precluding any causal conclusions.

### 2.9. Multiple comparison correction

Within the discovery-oriented framework of this study, we followed a balanced approach to multiple comparison correction, separately applying false discovery-rate (FDR) correction to results from each post-GLM analysis step outlined above (Supplement 1.9 for details). For the sake of transparency with respect to the reported significance levels, throughout the manuscript, “p” refers to uncorrected, “q” to FDR-corrected, and “pFWE” to FWE-corrected p values.

## 3. Results

### 3.1. Demographic, hormonal, and psychological assessments

The baseline assessment was conducted on average 3.43 days postpartum (T0, SD=2.03 days). The longitudinal subgroup was measured again at postpartum day 22.42 (T1, SD=2.52), 43.57 (T2, SD=3.06), 64.61 (T3, SD=2.8), 84.91 (T4, SD=3.71), and 173.32 (T5, SD=10.15). Tables S1–3 provide detailed sample characteristics. Postpartum and nulliparous groups differed in baseline age (mean=30.52 vs. 28.04, SD=3.54 vs. 4.91 years; Mann-Whitney-U-Test: U=1075.5, p<.001; Table S3). Compared to cross-sectional subjects, the longitudinal postpartum group showed higher age (32.68>29.79 years; U=292, p=.003), lower maximum motion (U=759, p=.006), a later baseline assessment (6.0>2.5 days postpartum; U=80, p<.001), higher postpartum depression symptoms (U=312, p=.007), and a higher rate of C-sections (63.16%>30.23%, χ^2^=11.71, p=.008; Tables S1 and S2). To assess the potential influence of these variables on the subsequent MRI results, we evaluated their associations with rsfMRI metrics in the baseline sample and included them as covariates in the cross-sectional and longitudinal sensitivity analyses.

In longitudinal analyses, we found a quadratic effect of postpartum week on log-transformed progesterone (β=.005, p=0.018), but not on estradiol levels or the estradiol/progesterone ratio. Maternal attachment (MPAS)^26^ and postpartum depression (EPDS)^25^ scores showed linear and quadratic effects of time (p<.05; Table S3).

### 3.2. Strong regional alterations of resting-state activity and connectivity in the first post-partum week

Our integrative approach to rsfMRI analysis^40^ revealed both increased and decreased local activity (fALFF) in postpartum mothers, while FC was consistently decreased (LCOR and GCOR; Table 1, Figure 1). Specifically, we found two clusters with increased local activity (fALFF) in postpartum relative to nulliparous groups in right precentral and left medial temporal areas, and one cluster with decreased local activity in the right superior frontal gyrus. For LCOR, postpartum women showed decreased local connectivity in bilateral insulae compared to controls. In contrast, GCOR was decreased in the postpartum group relative to controls in bilateral putamen and pallidum. In sensitivity analyses, group differences were robust against control for motion, GMV, and education (Table S4).

**Table 1:**
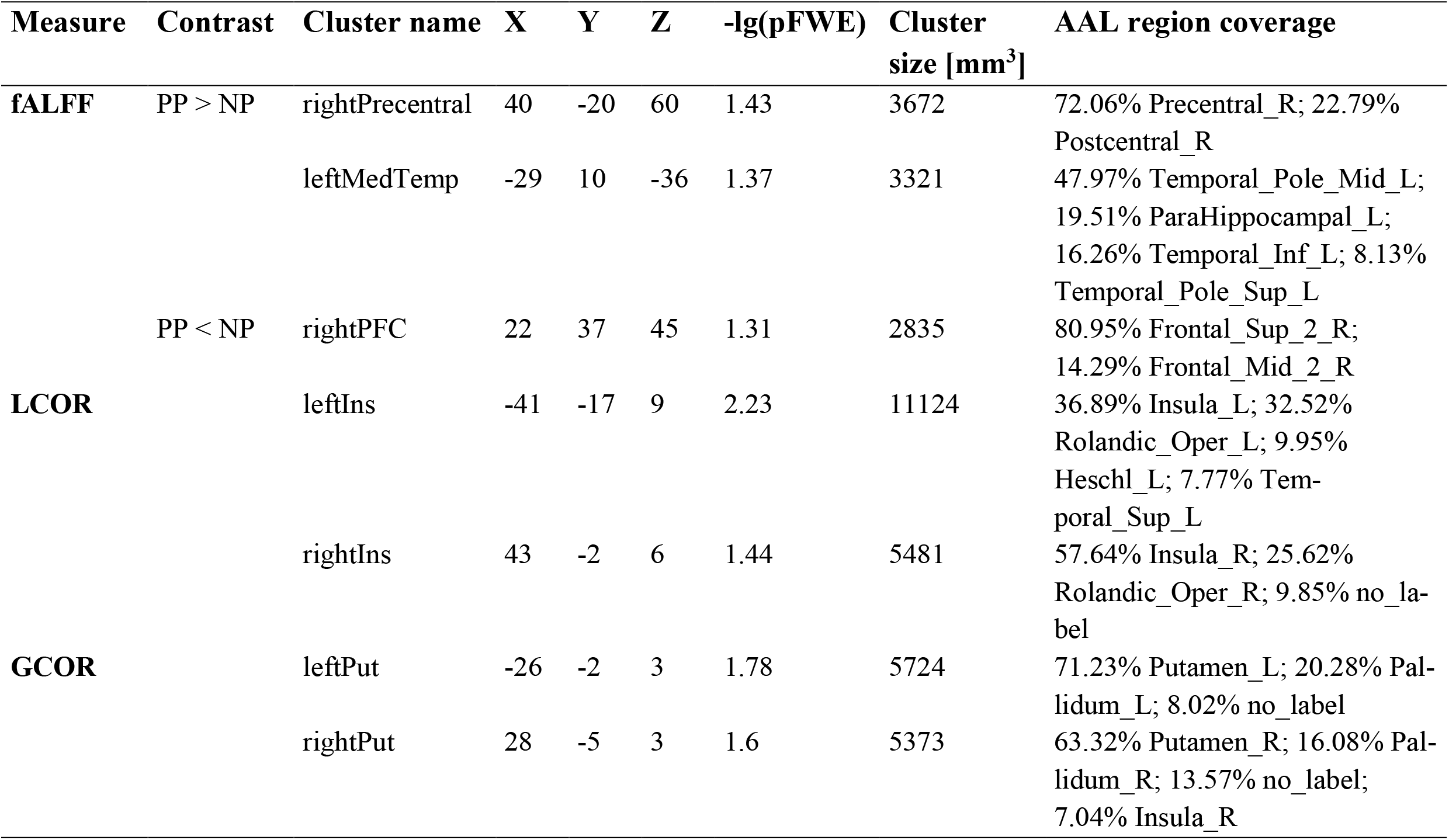
Statistical and anatomical characteristics of baseline rsfMRI clusters. Clusters were named after their main location. X, Y, and Z show coordinates in MNI-152 space. The non-parametric pFWE value associated with each cluster based on its cluster mass is displayed negative log10 transformed. Clusters were characterized anatomically based on the AAL atlas in terms of the percentage to which each AAL region was contained in each cluster. Abbreviations: AAL = Automated Anatomic Labeling atlas, fALFF = fractional amplitude of low frequency fluctuations, LCOR = local correlation, GCOR = global correlation, PP = postpartum, NP = nulliparous, MED = medial, PFC = prefrontal cortex, Ins = insula, Put = putamen.

**Figure 1.**
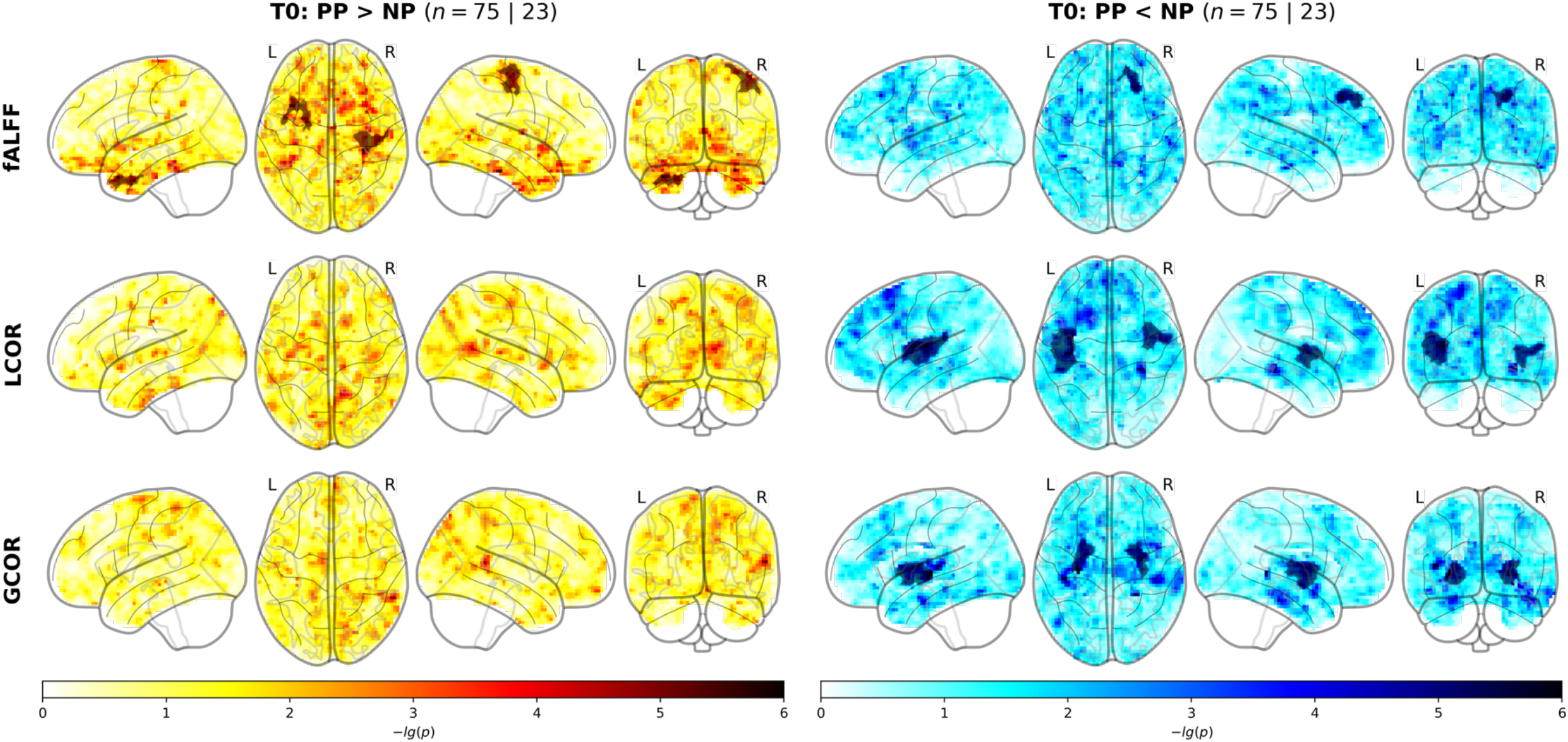
Clusters of differing local activity (fALFF), local connectivity (LCOR), and global connectivity (GCOR) in the nulliparous and postpartum groups at baseline. Cluster-level and whole-brain results from baseline (T0) rsfMRI analyses. Brain maps for each rsfMRI metric (rows) and each contrast (columns) show voxel-level negative log10-transformed p values overlaid by cluster-level results in darker shades (non-parametric cluster mass permutation). Abbreviations: PP = postpartum, NP = nulliparous, fALFF = fractional amplitude of low frequency fluctuations, LCOR = local correlation, GCOR = global correlation.

In the postpartum sample, the cluster-level rsfMRI results showed no clear association between the pregnancy-related variables and in-scanner motion (Figure S1). Most notably, barring a weak negative association between the prefrontal fALFF cluster and maximum motion (p=.045), no variable that differed between the two postpartum subgroups had an effect on the rsfMRI metrics.

### 3.3. Temporal dissociation of intra- and interregional rsfMRI metrics in the postpartum period

Assessing the temporal development of our baseline findings, we observed a general dissociation between clusters obtained from local measures (fALFF and LCOR) and the global connectivity measure (GCOR) (Figure 2, Tables S5–7). *Intra*-regional neocortical alterations of activity and connectivity, involving particularly the fALFF and right insular LCOR clusters, and less clearly the left insular LCOR, persisted across the 6 observed postpartum months. In contrast, the reduced *inter*-regional whole-brain FC of subcortical areas showed normalization patterns, with no significant or visible group differences between postpartum and nulliparous groups at about 6–9 weeks postpartum. In line with that, only the GCOR clusters showed significant linear and quadratic longitudinal development [left Put: β(linear)=.022, q=.003, β(quadratic)=-.0016, q=.014; rightPut: β(linear)=.020, q=.002, β(quadratic)=-.0018, q=.002]. Controlling all analyses for the underlying GMV reduced effect sizes, while group differences and longitudinal trends remained stable (Tables S8–10). Temporal GCOR trajectories were robust against baseline age, baseline postpartum week, in-scanner motion, day of gestation, birth mode, baby weight or sex, breastfeeding, or total number of children (Table S11). We also ensured that the findings were not driven by regression-toward-the-mean effects (Supplement 2.1, Figure S2, Table S12).

**Figure 2:**
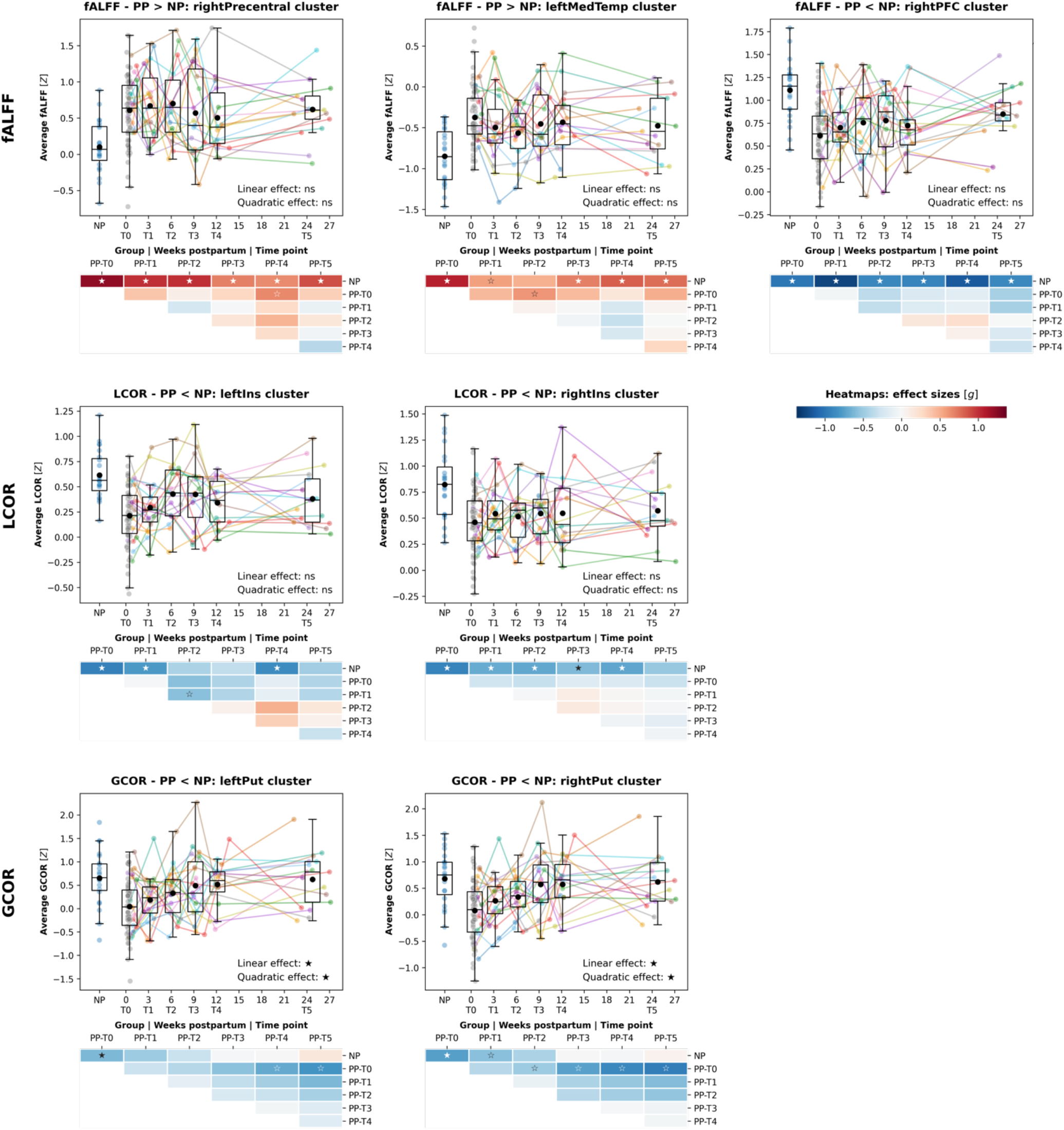
Longitudinal development of rsfMRI clusters. Development of baseline cluster-wise averaged local and global connectivity and activity metrics across 6 postpartum months. Boxplots: x-axes show time in weeks postpartum (dimensional scale), y-axes show the Z-standardized rsfMRI metric. Each dot represents one subject at one individual time point, lines connect longitudinal scans. Boxplots show the distribution across subjects at each time point (black dot = mean, middle line = median, boxes = quartile 1 and 3, whiskers = range if below 1.5 * interquartile range from quartile 1 or 3). Heatmaps show effect sizes (Hedge’s g) of within- and between-group comparisons, overlaid labels mark significances (filled star: q<.05, empty star: p<.05, none: p>.05). Significance of linear mixed models is printed in the lower right corner. Abbreviations: fALFF = fractional amplitude of low frequency fluctuations, LCOR = local correlation, GCOR = global correlation, PP = postpartum, NP = nulliparous, ns = not significant.

Whole-brain beta maps for the effect of postpartum week on each rsfMRI metric in the longitudinal sample (Figure S3) colocalized with the corresponding cross-sectional beta maps comparing early postpartum women with controls (Figure S4). In line with the strong longitudinal effects, the two GCOR maps in particular showed a high degree of colocalization [Z(rho)=-.90, p(exact)=.0001], supporting our focus on clusters obtained in the well-powered baseline comparison. Furthermore, cross-sectional fALFF and LCOR beta maps colocalized with each other [Z(rho)=.60, p(exact)=.0001] but not with the GCOR map, indicating that the local and global metrics captured different biological processes (Figure S4).

### 3.4. Whole-brain rsfMRI changes colocalize with potentially mediating biological systems

The colocalization analyses aimed at identifying possible biological mechanisms underlying the observed postpartum rsfMRI adaptations revealed multiple spatial associations and temporal trajectories in line with the cluster-level findings. The baseline postpartum changes of fALFF and LCOR showed comparable patterns, colocalizing positively with progesterone [q(norm)=.019] and negatively with the mGluR5 distribution [fALFF: q(norm)<.001; LCOR: q(norm)=.038]. GCOR changes colocalized positively with cortisol and GABAA receptors [NR3C1: q(norm)=.030; NR3C2: q(norm)=.019] but negatively with progesterone [q(norm)=.030] and oxytocin receptors [q(norm)=.019; Figure 3, upper right; Table S13]. The spatial relationships, especially for GCOR, were driven by brain regions with the most pronounced postpartum changes (i.e., subcortical regions; Figure 3, left; Figure S5). In line with cluster-level results, fALFF/LCOR-mGluR5 and ALFF-PGR colocalizations (the latter less clearly) persisted across the first 6 postpartum months, while the GCOR colocalizations showed normalization trends with significant linear trajectories for cortisol, oxytocin, and GABAA receptors (NR3C1: β=-.007, q=.043; NR3C2: β=-.008, q=.027; OXTR: β=.007, q=.043; GABAA: β=-.006, q=.043; Figure 3, lower right; Tables S13 and S14). Baseline GCOR colocalization results replicated in the cross-sectional sample with longitudinal subjects being excluded (Supplement 2.1, Figure S2). A positive correlation was found between the cortisol and GABAA receptors, limiting specific interpretation of their colocalization patterns (Figure S6).

**Figure 3:**
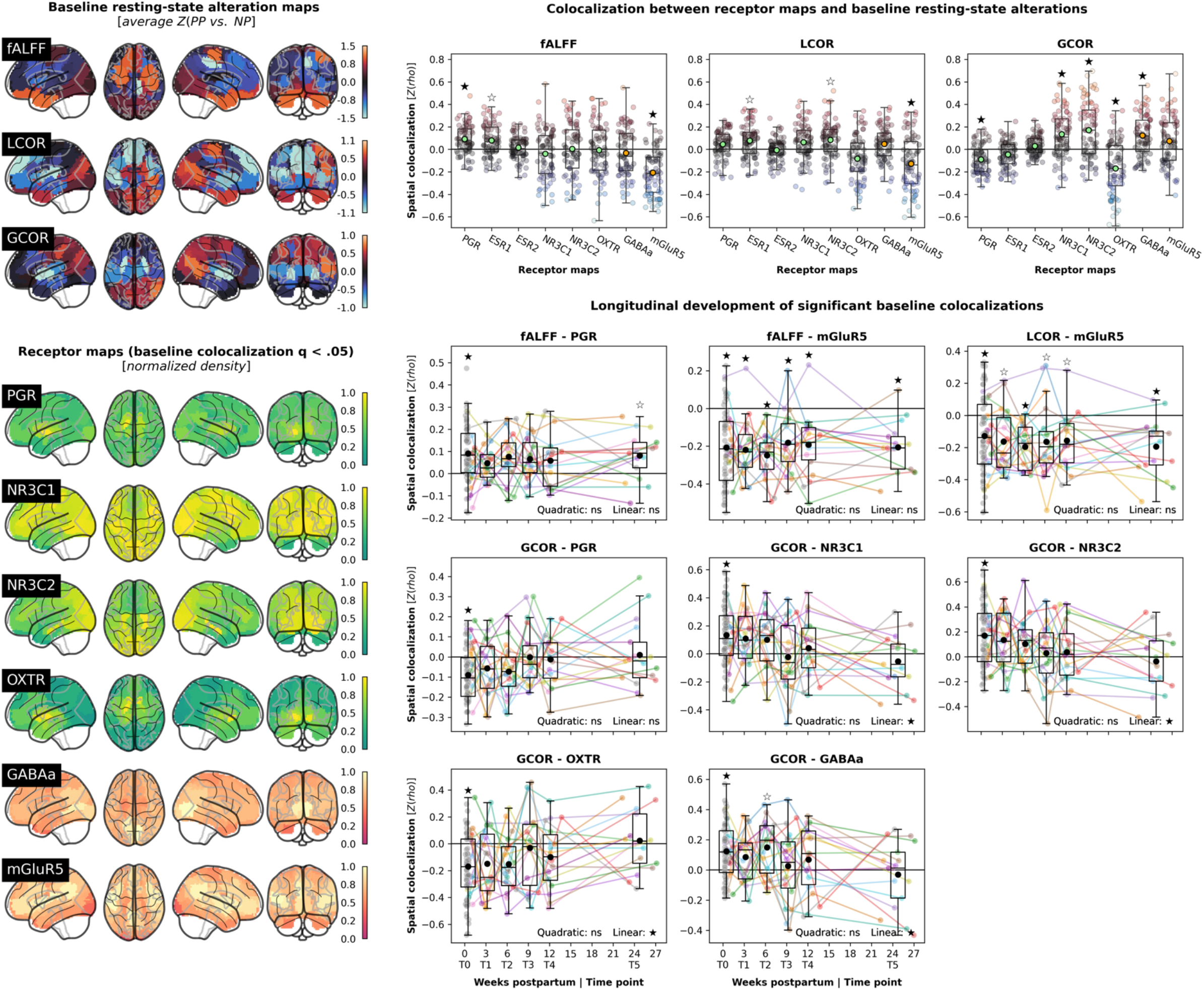
Subject-level spatial colocalization between postpartum rsfMRI alterations and hormonal/neurotransmitter receptor densities. Upper left: Baseline (T0) rsfMRI changes in postpartum women relative to the control cohort; average across subject-wise parcel-level Z score maps. E.g., for GCOR, the bright blue subcortical regions correpond to the initial GLM finding (PP<NP). Lower left: Parcellated receptor maps for which significant baseline colocalization with any rsfMRI metric was detected; normalized to range 0–1. E.g., the positive spatial colocalization of GCOR with PGR and OXTR is driven by decreased GCOR (PP<NP) and high PGR/OXTR density in subcortical regions. Upper right: Baseline analyses testing for colocalization between receptor distributions and rsfMRI data of each postpartum subject at baseline relative to the control group. X-axis: Z-transformed Spearman correlation coefficients, y-axis: rsfMRI metrics, scatter colors: colocalization strength. Lower right: Longitudinal analyses following up on each baseline results if associated group permutation q(norm)<.05 (filled stars). The plot design equals Figure 2 with the y axis showing Z-transformed Spearman correlation coefficients. Abbreviations: fALFF = fractional amplitude of low frequency fluctuations, LCOR = local correlation, GCOR = global correlation, ns = not significant.

### 3.5. Receptor distributions provide a multivariate explanation of postpartum rsfMRI changes

Multivariate models “predicting” baseline rsfMRI alteration patterns in postpartum women from the 6 colocalizing receptor maps^20,21^ showed that receptor distributions explained baseline alterations of GCOR to 15.8% [average, range 0–42.2%, p(exact)=.0180] and fALFF to 13.8% [0– 36.3%, p(exact)=.0004] but not of LCOR [8.9%, p(exact)=.5280].

### 3.6. Temporally dissociated intra- and interregional rsfMRI metric changes show diverging behavioral and hormonal association patterns

The temporal and spatial dissociation between postpartum alterations of local and global rsfMRI metrics indicates diverging underlying mechanisms. Given the strong hormonal fluctuations of the first postpartum weeks (Figure 4, right), it would be plausible if the observed transitory rsfMRI changes were hormone-related, while the long-lasting changes might be linked to more permanent behavioral or psychological adaptations following pregnancy and childbirth. No LMM targeting these temporal relationships survived FDR correction (Figure 4, left; Tables S15 and S16). Seeking to discover potential links for future targeted exploration, we nevertheless evaluated the emerging patterns of association (all p<.05). In line with the above hypotheses, GCOR-derived metrics seemed generally associated with hormone levels, most notably with progesterone, indicating a potential biological correlate of postpartum long-range connectivity adaptations. In contrast, we observed weaker associations of postpartum depression symptoms and mother-child attachment with local resting-state metrics (LCOR and depression/EPDS; fALFF and attachment/MPAS), in line with persistent changes in postpartum brain function as a possible neural signature of behavioral adaptations. We maintain, however, that these findings are of a preliminary nature. They do not support any causal claims and require targeted evaluation in an independent sample.

**Figure 4:**
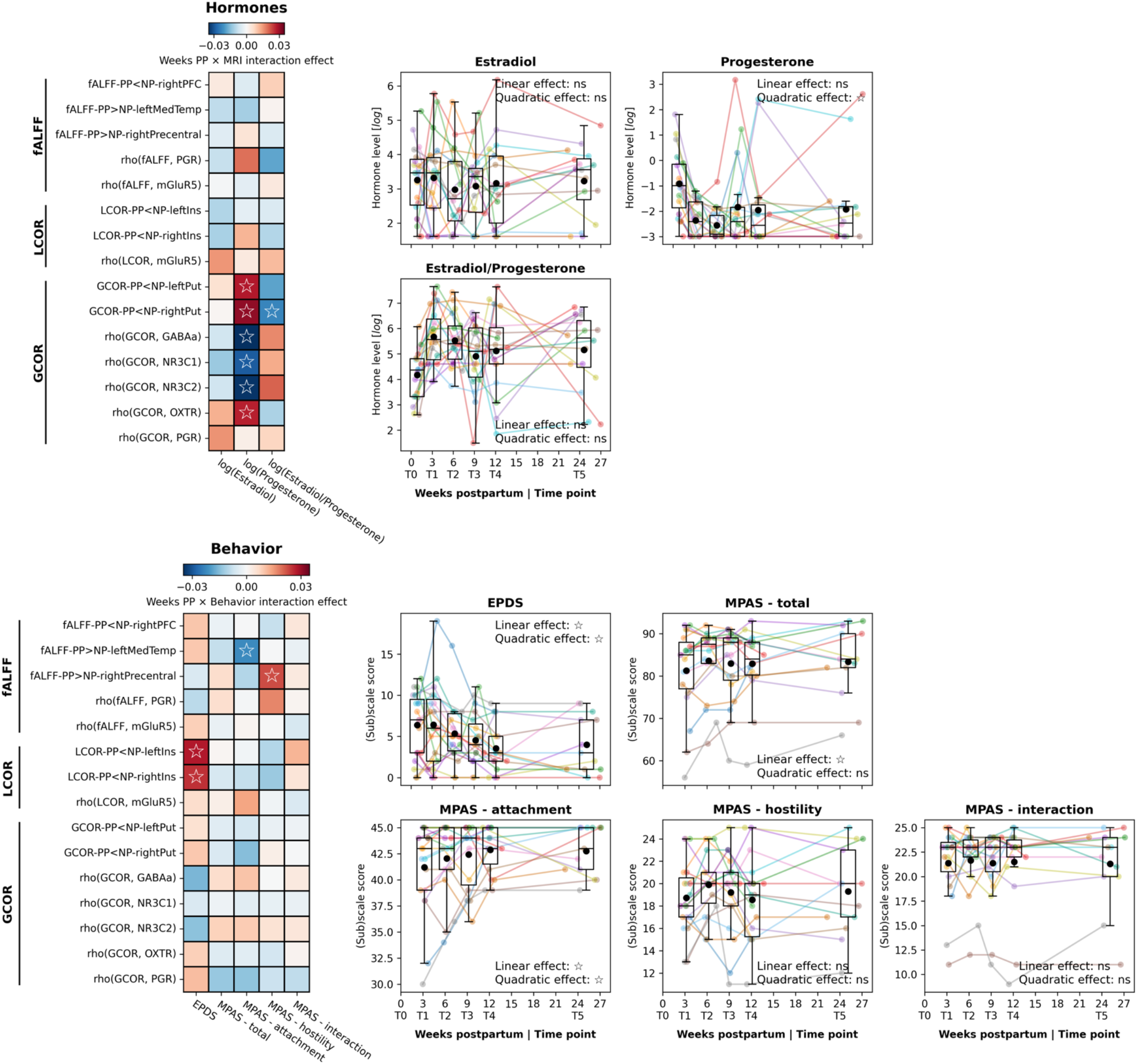
Patterns of association over time between rsfMRI results and hormone levels as well as behavior. Heatmaps: Results of linear mixed models to determine time-dependent associations between rsfMRI metrics (cluster-level results, spatial colocalizations with q<.05) and hormone levels (upper) as well as mother-child attachment and postpartum depression symptoms (lower). Heatmap colors represent standardized effect sizes of interaction effects between weeks postpartum and (upper) hormone levels to “predict” MRI estimates or (lower) MRI estimates to “predict” behavioral scales. Overlaid labels show p values (empty star: p<.05, no test survived false-discovery rate correction). The presented results are intended to identify general patterns of association rather than to interpret specific tests. Scatter-/Boxplots: Levels or scores, respectively, of the tested hormones (upper) and behavioral scales (lower) in the postpartum sample over time (see Figure 2). Abbreviations: fALFF = fractional amplitude of low frequency fluctuations, LCOR = local correlation, GCOR = global correlation, PP = postpartum, NP = nulliparous, MPAS = Maternal Postnatal Attachment Scale, EPDS = Edinburgh Postnatal Depression Scale.

## 4. Discussion

In this study, we tracked the reorganization of brain function in healthy mothers from shortly after childbirth until 6 months postpartum. While neocortical intraregional resting-state activity and connectivity alterations persisted throughout the study period, interregional connectivity of subcortical areas normalized to nulliparous levels within the first 2 postpartum months. Whole-brain colocalization patterns with hormone and related neurotransmitter distributions, as well as temporal associations to maternal hormone levels, indicated the sex steroid system as a potential underlying biological target, warranting future research.

The first 6 postpartum weeks, referred to as the (*sub-)acute postpartum period*, are marked by extensive physiological and hormonal adaptations^51^. In postpartum women, the strongly decreased bilateral putamen FC and its normalization during this period in interaction with progesterone blood levels suggest effects of progesterone on neural function, in line with progesterone-specific and general steroid hormone effects on basal ganglia morphology and function^52,53^. Supporting this relationship beyond the level of brain regions, postpartum inter-regional connectivity changes were found to negatively colocalize with progesterone and oxytocin receptors across brain regions (*less* FC in areas with *higher* receptor gene expression) and positively colocalize with cortisol and GABAergic receptors. The colocalizations with inter-regional FC alterations reduced with postpartum time, possibly indicating that hormonal adaptations may underly a postpartum reorganization of the functional connectome. Previous research had suggested a link between the post-childbirth drop in progesterone and its metabolites and the development of baby blues and postpartum depression through dysregulated GABAergic neurotransmission^54^. The observed effects on a specific rsfMRI metric indicate promising targets for future research on postpartum depression. However, it is still unclear how these postpartum physiological adaptations impact the mental health of postpartum mothers. Genetic predisposition and life experiences likely contribute to the individual susceptibility to mood disorders amidst hormonal fluctuations^55^. Future studies in populations at risk for postpartum depression can utilize our findings to identify clinically valuable biomarkers.

It is unclear when the post-pregnancy recovery of the maternal brain is completed and whether pregnancy has long-lasting neural effects. Structural changes can be detected early in the postpartum phase^11,56,57^, lasting up to 6 months^56^ or even longer^7,58^. They encompass a widespread network including prefrontal, cingulate, temporal, and insular cortices. We observed persistent changes in local activity in the same brain regions, which are thought to be part of the “parental caregiving” brain network^59^. These changes were strongest in areas with higher progesterone receptor gene expression and lowest in areas with higher glutamate receptor density, possibly indicating underlying biological mechanisms for further evaluation. While the local rsfMRI metrics did not show normalization patterns, there was preliminary evidence of associations with postpartum behavioral/psychological (mal-)adaptations. In particular, a potential association between intra-insular connectivity and postpartum depression symptoms warrants further investigation^60,61^. Hoekzema et al.^19^ reported long-term effects of pregnancy or motherhood on resting-state connectivity within the DMN. Although direct comparisons are limited due to major differences in the covered peripartum time period and analysis methodology, our results align with those of Hoekzema et al. with respect to potential associations of resting-state changes and mother-child bonding. Given prior reports of time-sensitive relationships between postpartum amygdala volume and hostile behavior toward the child^2^, as well as the protective neural effects of parenting, particularly in striatal and limbic regions^8–10,24^, it may be safe to assume that the experience of birth and motherhood influences the maternal brain.

The neural phenomena observed in this study, along with potentially underlying biological mechanisms, may constitute important postpartum adaptation processes^51^. However, factors such as pain, medication, inflammation, fluid retention, or sleep disruption may similarly influence neurophysiology, requiring additional research. Although well suited to examine the adaptation trajectories of brain function, our study was limited by (i) the small longitudinal postpartum sample, (ii) some clinical differences between the cross-sectional and longitudinal samples, and (iii) the lack of a follow-up in the nulliparous group, posing the risk of complex statistical models being underpowered. Due to the lack of pre-/during-pregnancy rsfMRI, we could not disentangle the effects of pregnancy from those of postpartum adaptations or preexisting factors. The preliminary hormonal and behavioral association patterns require replication in larger groups and should be assessed in postpartum and control participants alike. Finally, spatial colocalization analyses are generally limited by their inability to prove causal relationships and their reliance on heterogeneous external datasets. This applied in particular to the hormone receptor maps, which originated from 6 post-mortem brains, including only one 49-year-old female, and which are not guaranteed to correspond to in-vivo protein activity^62^. Finally, specific colocalization results ought to be interpreted with care as it is unclear to which extent the partially intercorrelated receptor maps constitute independent biological entities with potential effects on rsfMRI signals.

### 4.1. Conclusion and future outlook

We provide evidence of dynamic functional adaptation processes in the early postpartum brain. While the subcortex-centered global connectivity changes may be of transient nature and related to hormonal adaptations, neocortical activity and connectivity show persistent alterations, possibly associated with behavioral and psychological processes. Building on our data, further longitudinal studies involving larger cohorts and pre-pregnancy assessments will bring valuable insights into the post-childbirth recovery of brain morphology and function. Along with longitudinal multimodal neuroimaging, cognitive, and lifestyle assessments, tracking of ovarian hormones from pre-pregnancy through the postpartum phase will help identify factors contributing to the phenomena observed in this study. It is imperative to understand the long-term effects of pregnancy-related adaptations, which, in addition their protective role, may also render women more vulnerable to certain risks. It needs to be ascertained if these changes contribute to the development of psychiatric conditions, especially in women at higher risks for postpartum psychiatric disorders.

## 5. Data sharing

Due to privacy protection, we cannot provide subject data openly. All group-level results are provided in supplementary tables. Group-level MRI volumes will be uploaded to Neurovault following publication. Individual data are available from one of the corresponding authors, NC, upon reasonable request.

## Supporting information

Supplementary material (analysis code notebook)

## 6. Acknowledgments

Leon D. Lotter received financial support from the Max Planck Society (MPG) and the German Ministry of Education and Research (BMBF), Germany. This work was supported by the Brain Imaging Facility of the Interdisciplinary Center for Clinical Research (IZKF) Aachen within the Faculty of Medicine at RWTH Aachen University.

## 7. Author contributions

Conceptualization: NC, SN. Data Curation: NC, SN, EL. Formal Analyses: LL. Funding acquisition: NC. Investigation: NC, SN, EL. Methodology: JD, LL. Project administration: NC, SN. Supervision: NC, JD. Visualization: LL. Writing – original draft: NC, LL. Writing - review and editing: all authors. All authors confirm that they had full access to all data in the study and accept responsibility to submit for publication.

