## Supplementary material (text & figures) for "Temporal dissociation between local and global functional adaptations of the maternal brain to childbirth: A longitudinal assessment"

#### Supplementary Materials

Leon D. Lotter*, Susanne Nehls, Elena Losse, Juergen Dukart, Natalia Chechko*

* Corresponding authors: nchechko [at] ukaachen.de; l.lotter [at] fz-juelich.de

Content:

Supplementary Methods and Results

Supplementary Figures S1–S6

Supplementary Tables: see separate Excel file

Analysis code: see separate Jupyter notebook

### 1. Supplementary Methods

#### 1.1. Study samples

Postpartum women were recruited in the Department of Gynecology and Obstetrics, University Hospital Aachen, Germany, within two studies run in parallel by the Department for Psychiatry, Psychotherapy, and Psychosomatics at the same hospital. Both studies were led by the same investigators and applied the same testing protocols for MRI, hormonal, and behavioral measurements. Study 1 targeted early postpartum brain changes with a focus on early prediction of postpartum depression[1]. Cross-sectional morphological data was published in Chechko et al. (2022)[2]. In Study 2, a healthy subgroup of women recruited for Study 1 (no psychiatric diagnosis, no risk factors, narrow age span) was followed up for a longitudinal exploration of specific aspects of the postpartum period. Longitudinal morphological changes and their behavioral associations up to postpartum week 12 were assessed in Nehls et al. (2023)[3]. Additionally, a yet unpublished nulliparous control sample was recruited for the baseline comparison, scanned once at the same scanner. While our prior studies took a psychological viewpoint, focusing on psychopathological and behavioral correlates of postpartum brain changes, here we approached the topic from a physiological perspective, using the dense 6-months longitudinal data to gain insights into biological mechanisms underlying these neural adaptations.

#### 1.2. Hormonal assays

Progesterone and estradiol serum concentrations were measured before each scanning session and analyzed by competitive immunometric electrochemistry luminescence detection at the Laboratory Diagnostic Center, University Hospital RWTH Aachen, Germany. The samples were run on a Roche Cobas e601 and on a Roche Cobas e801 with Cobas Elecsys estradiol and progesterone reagent kits, respectively (Roche Diagnostics, Bromma, Sweden). For progesterone, the measurement interval was .05–60 ng/ml with an intra-assay coefficient of variation of 2.33–2.49%. For estradiol, the measurement interval was 5–3000 pg/ml with a coefficient of variation of 1.77–2.91%.

#### 1.3. Software

Processing of functional and structural brain images was conducted in a MATLAB (R2022a) environment using CONN (21a)[4], building on SPM12 routines. Statistical analyses were calculated in a Python (3.9.12) environment using Nilearn (0.10.0)[5] for voxel-wise rsfMRI group comparisons. Resulting spatial clusters were characterized using atlasreader (0.1.2)[6]. Spatial colocalization estimates were calculated in JuSpyce (0.0.3)[7] available from https://github.com/LeonDLotter/JuSpyce. Further statistical analyses were conducted with statsmodels (0.13.5) and pingouin (0.5.3)[8,9]. Brain gene expression and nuclear imaging data were obtained and processed with abagen (0.1.3)[10] and neuromaps (0.0.3)[11]. Visualizations were created with matplotlib (3.5.2)[12], seaborn (0.12.1)[13], and Nilearn.

##### 1.4. MRI acquisition and preprocessing

All participants were scanned at the same Siemens Magnetom Prisma 3T MRI scanner. MRI Structural T1-weighted images were acquired using a Rapid Acquisition Gradient Echo sequence with following parameters: TR = 2300 ms, TE = 1.99 ms, FoV = 256 x 256 mm, number of slices = 176, voxel size = 1 x 1 x 1 $mm^3$. For rsfMRI, all probands underwent a 6.6 minutes gradient-echo Echo Planar Imaging protocol with TR = 2200 ms, TE = 30 ms, flip-angle = 90 °, FoV = 200 x 200 mm, number of slices = 36, number of volumes = 300, voxel-size = 3.1 x 3.1 x 3.1 $mm^3$.

Preprocessing of functional images consisted of removal of the first four frames, realignment for motion correction, and co-registration to structural images with subsequent spatial normalization into Montreal Neurological Institute space using parameters derived from structural data. The normalization parameters were applied with modulation to segmented gray matter probability maps to obtain corresponding voxel-wise GMV. Functional and structural images were interpolated to 3-mm and 1-mm isotopic resolution, respectively. A Gaussian smoothing kernel of 6-mm full width at half maximum was applied to rsfMRI data. Twenty-four motion parameters along with mean white matter and cerebrospinal fluid signals were regressed out of the functional data[14]. The resulting images were linearly detrended and temporally bandpass filtered (.01–.08 Hz). A gray matter mask (probability > .2) was applied to all images to restrict analyses to gray matter tissue. For spatial colocalization analyses, data were parcellated into 100 cortical and 16 subcortical parcels[15,16]. All structural and functional MRI volumes before and after preprocessing were visually quality-controlled and subject exceeding motion cut-offs of a mean and maximum framewise displacement of .5 and 3 mm were excluded.

##### 1.5. Controlling for regression-toward-the-mean effects

We systematically tested if the observed temporal normalization effects were due to regression-toward-the-mean effects. For this, we (i) replicated baseline voxel-level GLM and spatial colocalization analyses while *excluding* the subjects with longitudinal data, (ii) extracted cluster-average data from subjects *with* longitudinal data using the clusters estimated on the independent cross-sectional sample, and (iii) tested for longitudinal linear and quadratic effects in the cluster-average data.

##### 1.6. Spatial colocalization between rsfMRI effect size maps

In our longitudinal analyses, we focused on the development of cluster-level average results instead of performing GLMs in only the longitudinal sample. However, to demonstrate that the longitudinal whole-brain-level effects mirrored those observed in the cross-sectional (case-control) analysis, we compared beta maps obtained from both approaches to each other using spatial colocalization analyses. In this context, we also calculated the degree of spatial colocalization between rsfMRI effect maps from different metrics (fALFF, LCOR, GCOR) to each other within and across the cross-sectional and longitudinal analyses.

We obtained beta effect maps quantifying (i) the baseline group difference for the three rsfMRI metrics (*"rsfMRI ~ group + age"*, NP < PP) and (ii) the effect of postpartum time on the rsfMRI metrics (*"rsfMRI ~ weeks_postpartum + age"*). We (i) parcellated the maps into 116 parcels covering cortex[17] and subcortex[18], (ii) Z-standardized each map, (iii) regressed parcel-wise grey matter probabilities from a standard MNI152 template from each map to account for partial volume effects[19,20], and (iv) correlated the 6 vectors (3 rsfMRI metrics × 2 analyses) with each other using Spearman correlations. As parametric p values from these analyses convey little meaning, we generated 10,000 null maps for each receptor map using a spatial autocorrelation-preserving procedure[21] and estimated exact p values from null distributions obtained by correlating – for each pair of maps – the original map with one of the null maps. This resulted in two p values for each pair of maps, which we finally averaged. The process was implemented using JuSpyce[22].

##### 1.7. Sources of receptor maps

Corticosteroid hormone and oxytocin receptor maps (PGR, ESR1/ESR2, NR3C1/NR3C2, OXTR) were constructed from postmortem Allen Brain Atlas gene expression data (1 female, ages 24.0–57.0 years, mean = 42.50, SD = 13.38)[23] in 116-region volumetric atlas space. The abagen toolbox was used for map construction, while applying bilateral mirroring across hemispheres. GABAergic ($GABA_A$) and glutamatergic (mGluR5) receptor maps were obtained with positron emission tomography from independent groups of healthy adult subjects ($GABA_A$: [11C]flumazenil, n = 10, mean = 26.60, SD = 7.00 years; mGluR5: [11C]ABP688, n = 73, mean = 19.90, SD = 3.04 years)[24–26].

##### 1.8. Spatial colocalization analyses

Spatial relationships between postpartum rsfMRI alterations and receptor maps were tested for by (i) parcellating all rsfMRI and receptor data in 116 cortical and subcortical parcels, (ii) regressing average grey matter probability taken from a MNI152 standard template from all maps across parcels to control for partial volume effects[19,20], (iii) calculating parcel-wise rsfMRI Z scores for each postpartum subject at a given time point by subtraction of the mean and division by the standard deviation of the rsfMRI metric in the nulliparous control sample[19], (iii) correlating these subject-level 116-parcel Z score maps with each receptor map using Spearman correlations, (iv) calculating the average Spearman "colocalization" across the postpartum sample, (v) generating a null distribution of average Spearman coefficients by repeating this process after permuting the PP-NP labels, (vi) estimating exact p values for each observed average Spearman correlation from this distributions ["p/q(exact)"], and (vii) fitting Gaussian curves to each null distribution to estimate p values with enough decimal places to accurately apply rank-based multiple comparison correction [i.e., FDR, referred to as "p/q(norm)"][27]. The group comparison concept was described in detail in Dukart et al.[19] and implemented via JuSpyce[7]. If significant colocalization was observed on the group level at baseline, longitudinal development of these colocalization metrics was tested for using LMMs as described in the main Methods.

A similar approach was used for multivariate regression colocalization. All receptors that showed significant colocalization with any of the three rsfMRI metrics were used as independent variables in three multivariate regressions per postpartum subject with each of the individual rsfMRI alteration maps as dependent variable. The resulting adjusted $R^2$ values (average across the postpartum sample), and 10,000 null values obtained by group permutation, were used to calculate exact p values for each metric.

In neuroimaging colocalization analyses, it is important to control for spatial autocorrelation patterns present in most brain maps. We assume that our approach of standardization on the control cohort paired with group permutation to be robust against these confounds, as (i) the standardization in permuted groups should effectively remove general spatial patterns present in all subject and (ii) the receptor map spatial autocorrelation should influence both the observed and the null colocalization estimates to similar degrees.

##### 1.9. Multiple comparison correction

Postpartum development of brain function is an under-investigated phenomenon, hindering the formulation of specific hypotheses. Therefore, our overarching study approach could be considered "exploratory", requiring targeted replication of all findings in independent samples. Within this context, we followed a balanced approach to multiple comparison correction, attempting to reduce excessive alpha error accumulation due to repeated testing on the one side but also avoiding false-negatives due to conservative correction on the other. First, we applied stringent non-parametric family-wise error correction within each rsfMRI GLM contrast (3 metrics × 2 contrasts = 6 tests) in the well-powered baseline sample (section 2.5). False-discovery correction (FDR) was applied independently to longitudinal rsfMRI post-hoc analyses (section 2.6): time point-wise ANCOVAs between postpartum and nulliparous groups (7 clusters × 6 time points = 42 tests), cluster-wise linear and quadratic LMMs (7 clusters × 2 = 14 tests), and paired t-tests between all postpartum time points (7 clusters × 15 combinations = 105 tests). Similarly, FDR correction was applied independently to baseline spatial colocalization analyses (section 2.7; 3 metrics × 7 receptors = 21 tests), to follow-up analyses of significant baseline colocalization at each longitudinal time point (8 metric-receptor pairs × 5 time points = 40 tests), and to linear/quadratic LMMs on the same data (8 metric-receptor pairs × 2 = 16 tests). Finally, the same correction was applied separately to preliminary MRI-hormone and -behavior interaction analyses (section 2.8; 15 rsfMRI metrics × 3 hormone levels = 45 tests; 15 metrics × 5 scales = 75 tests).

No multiple comparison correction was applied to group comparisons and longitudinal assessments of non-MRI demographic, clinical, and hormonal data (section 2.4) as these did mainly serve informational purposes. The same procedures were used in all sensitivity analyses to retain comparability with the main results.

#### 2. Supplementary Results

##### 2.1. Early postpartum normalization of global connectivity is not due to regression-to-the-mean effects

We performed control analyses to ensure that normalization effects observed in GCOR-related variables were not influenced by regression-toward-the-mean effects. The reduced postpartum sample (n = 56) showed the same bilateral putamen-related clusters of reduced global connectivity at baseline. Temporal development of the cluster-average data in independent longitudinal subjects followed similar trajectories as reported above. Baseline spatial colocalization patterns in the reduced sample were similar to those in the full sample (Figure S2, Table S12).

#### 4. Supplementary Figures

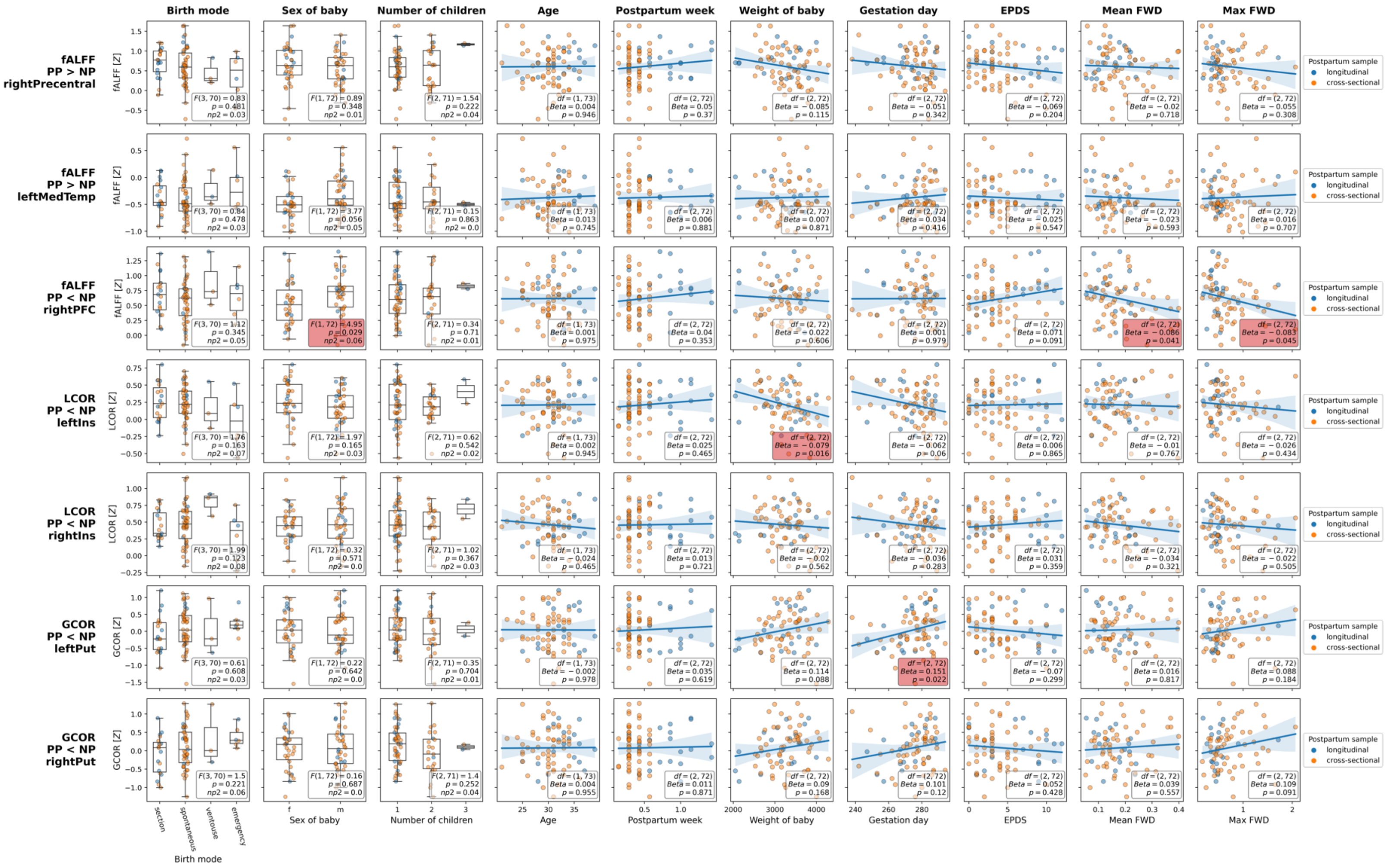


**Figure S1: Sensitivity association analyses between pregnancy-related variables, motion, and cluster-level rsfMRI results in the baseline sample.**

Y-axes: Cluster-level rsfMRI metrics. X-axes: pregnancy-related variables and in-scanner motion. For categorical variables (first three columns), ANCOVAs controlled for age were calculated to assess potential group differences. For continuous variables (column 4 onwards), we fitted linear regression analyses with age and the respective variable (except for the "age" analysis) as independent variables, reporting respective beta coefficient. P values were not corrected as we aimed to err on the side of false positives ($p < .05$ highlighted in red). Postparum subgroups are marked by scatter point color to allow for assessment of potential variation by subgroup. Note that, except for maximum FWD, no variable that showed group differences in the preceeding comparison of cross-sectional vs. longitudinal postpartum subjects (Tables S1 and S2) showed a significant association with rsfMRI metrics in this analysis. Abbreviations: PP = postpartum, NP = nulliparous, fALFF = fractional amplitude of low frequency fluctuations, LCOR = local correlation, GCOR = global correlation, EPDS = Edinburgh Postnatal Depression Scale, FWD = frame-wise displacement.

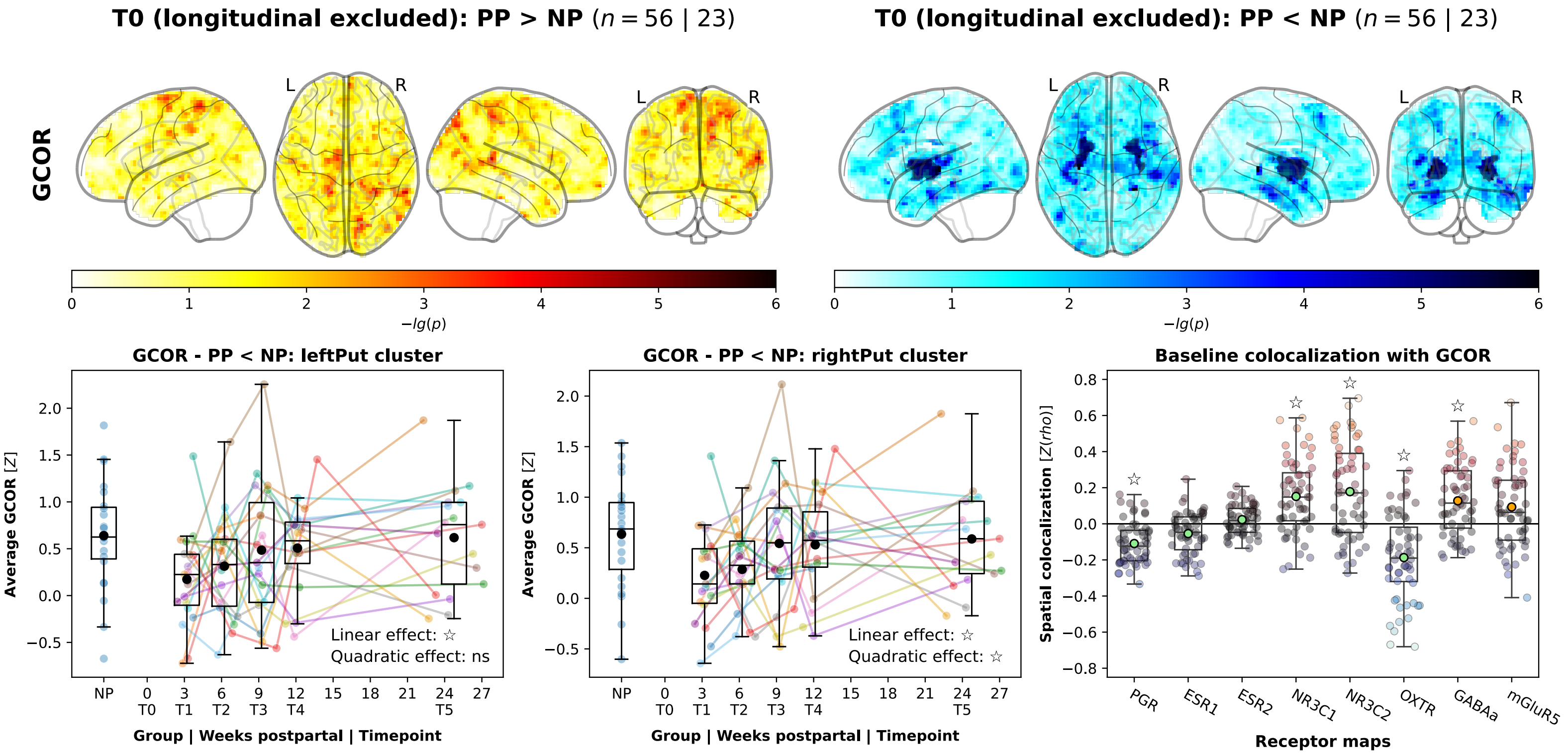


**Figure S2: GCOR sensitivity analyses to control for regression to the mean effects.**

Upper: GCOR whole-brain cluster-level analyses were repeated while excluding subjects with longitudinal data. See Figure 1 for descriptions of plot elements.The resulting clusters almost exactly mirrored the main result. Lower left and center: Longitudinal trajectories of the cluster-average data in the held-out longitudinal sample. See Figure 2 for plot elements. Lower right: Baseline colocalization results without longitudinal subjects. See Figure 3 for plot elements. Temporal trajectories and spatial colocalization results mirrored those from the main analyses. Abbreviations: GCOR = global correlation, PP = postpartum, NP = nulliparous.

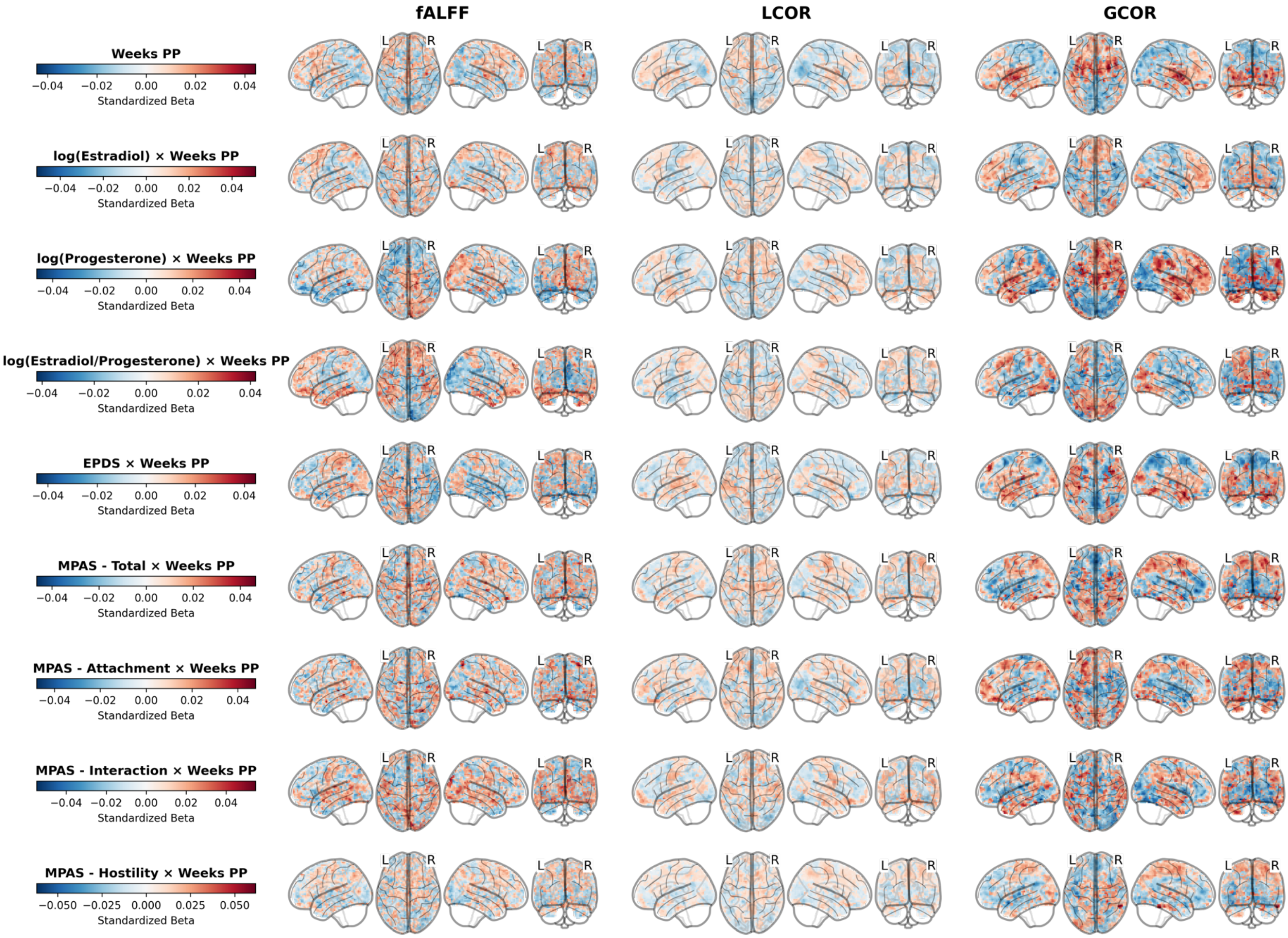


**Figure S3: Longitudinal effect maps of postpartum time, hormone levels, and behavior in the postpartum group.**

The plot is provided for reference and comparison with the cross-sectional baseline analyses (Figure 1) and longitudinal association analyses (Figure 4). Maps show effect sizes (standardized beta) obtained from estimating general linear models evaluating the main effect of weeks postpartum (first row) or the interaction between weeks postpartum and hormone levels (rows 2 to 4) or behavioral scales, respectively (row 5 onwards). Figure S3 shows the spatial correlation between the cross-sectional (PP vs. NP) and longitudinal (weeks postpartum) effect size maps. Abbreviations: PP = postpartum, NP = nulliparous, fALFF = fractional amplitude of low frequency fluctuations, LCOR = local correlation, GCOR = global correlation, MPAS = Maternal Postnatal Attachment Scale, EPDS = Edinburgh Postnatal Depression Scale.

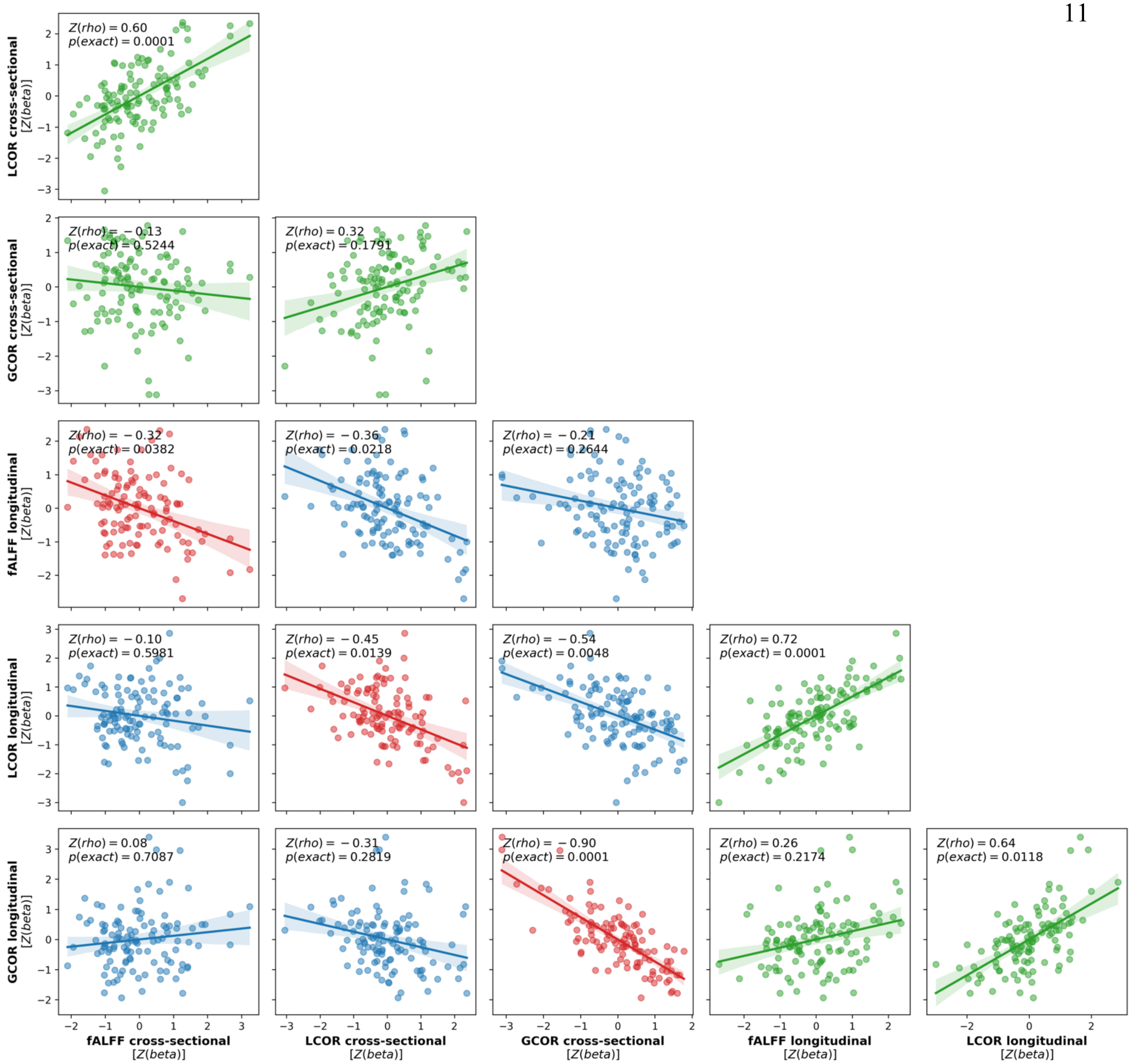


**Figure S4: Spatial correlation between effect size maps obtained from cross-sectional (PP vs. NP) and longitudinal (weeks postpartum) GLMs for all rsfMRI metrics.**

Axes: Z-transformed beta values from the cross-sectional PP vs. NP group comparison GLM (see Figure 1; "rsfMRI ~ group + age") and longitudinal effect of weeks postpartum on each rsfMRI metric (see Figure S2, row 1; "rsfMRI ~ weeks_postpartum + age"). Note that, due to Z-standardization and grey matter probability regression, zero on the scale does correspond to zero effects. Scatters: Effect size for each of 116 parcels. Line: Linear regression fit with bootstrapped 95% confidence intervals. Statistics: Spearman's rho and exact p value calculated from a null distribution obtained from correlation with 10,000 spatial autocorrelation-preserving null maps. The p value for each pair is the average of two p values calculated from null distributions after permutation of either one map. Red: Colocalization between longitudinal and cross-sectional maps from the same rsfMRI metric. Green: Colocalization between different rsfMRI metrics from either the cross-sectional (upper left) or longitudinal analysis (lower right). Abbreviations: PP = postpartum, GLM = general linear model, NP = nulliparous, fALFF = fractional amplitude of low frequency fluctuations, LCOR = local correlation, GCOR = global correlation.

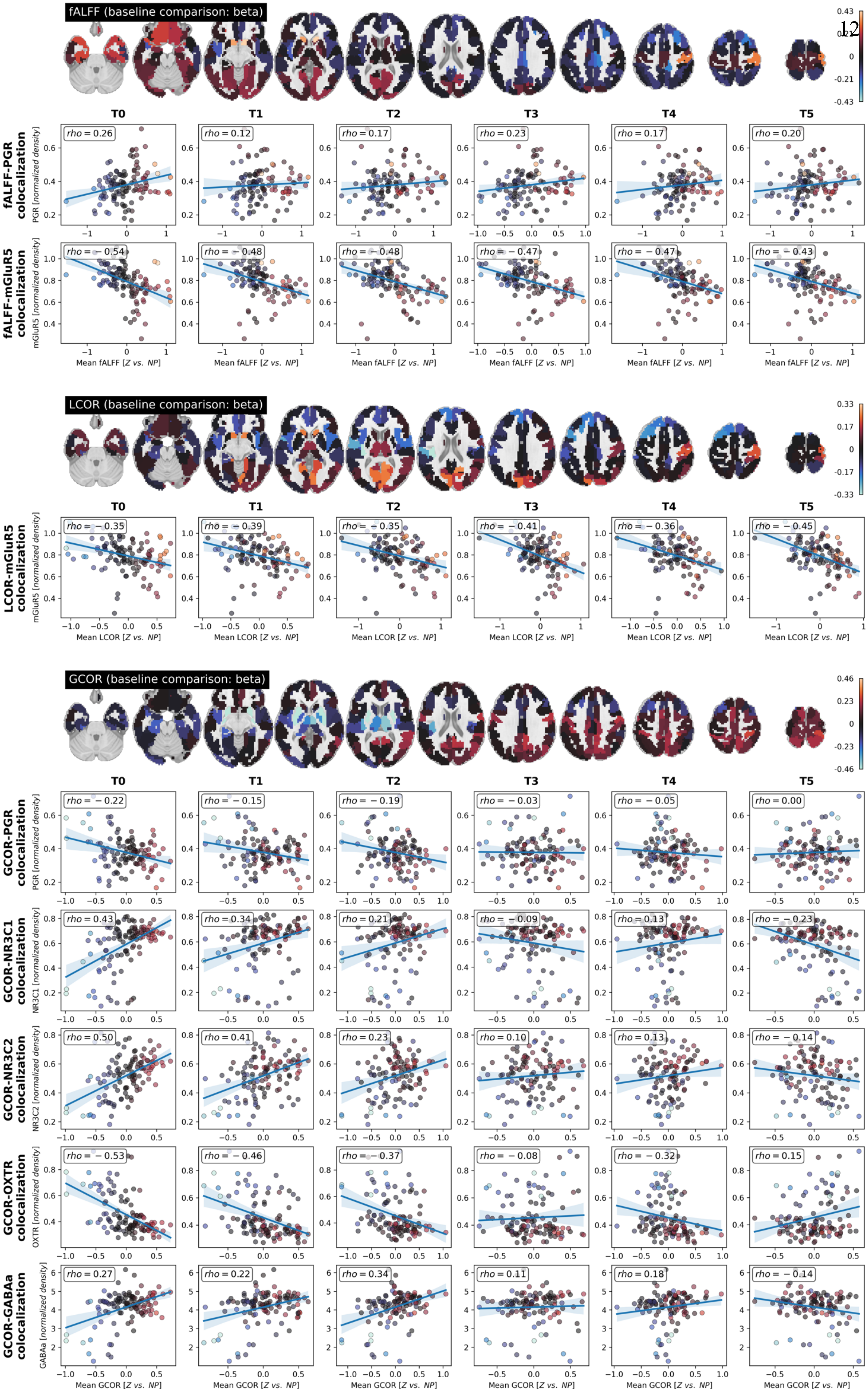

fALFF (baseline comparison: beta)
0.43
0
-0.21
-0.43
T0
T1
T2
T3
T4
T5
fALFF-PGR colocalization
PGR [normalized density]
rho = 0.26
rho = 0.12
rho = 0.17
rho = 0.23
rho = 0.17
rho = 0.20
fALFF-mGluR5 colocalization
mGluR5 [normalized density]
rho = −0.54
rho = −0.48
rho = −0.48
rho = −0.47
rho = −0.47
rho = −0.43
Mean fALFF [Z vs. NP]
LCOR (baseline comparison: beta)
0.33
0.17
0
-0.17
-0.33
LCOR-mGluR5 colocalization
mGluR5 [normalized density]
rho = −0.35
rho = −0.39
rho = −0.35
rho = −0.41
rho = −0.36
rho = −0.45
Mean LCOR [Z vs. NP]
GCOR (baseline comparison: beta)
0.46
0.23
0
-0.23
-0.46
GCOR-PGR colocalization
PGR [normalized density]
rho = −0.22
rho = −0.15
rho = −0.19
rho = −0.03
rho = −0.05
rho = 0.00
GCOR-NR3C1 colocalization
NR3C1 [normalized density]
rho = 0.43
rho = 0.34
rho = 0.21
rho = −0.09
rho = 0.13
rho = −0.23
GCOR-NR3C2 colocalization
NR3C2 [normalized density]
rho = 0.50
rho = 0.41
rho = 0.23
rho = 0.10
rho = 0.13
rho = −0.14
GCOR-OXTR colocalization
OXTR [normalized density]
rho = −0.53
rho = −0.46
rho = −0.37
rho = −0.08
rho = −0.32
rho = 0.15
GCOR-GABAa colocalization
GABAa [normalized density]
rho = 0.27
rho = 0.22
rho = 0.34
rho = 0.11
rho = 0.18
rho = −0.14
Mean GCOR [Z vs. NP]

**Figure S5: Spatial colocalizations between average postpartum resting-state alterations and receptors maps in relation to the baseline resting-state group comparison.**

Brain plots of the beta maps (NP > PP) for each rsfMRI metric accompanied by scatter plots of significant (q < .05) spatial colocalizations between the respective resting-state alterations in postpartum women (y-axes; average parcellated data of postpartum women at each timepoint standardized by parcellated data from the control cohort) and receptor maps (x-axes). Scatter points: brain parcels, average postpartum Z score relative to the control cohort, colored by parcel-average effect sizes from the cross-sectional baseline GLM (PP vs. NP) as displayed in the brain maps for each rsfMRI metric. Statistic: Spearman's rho. Line: linear regression fit and bootstrapped 95% confidence interval. This plots aims to demonstrate that the observed spatial colocalizations were driven by initial cross-sectional GLM findings. E.g., for GCOR, the bright blue colors in subcortical regions that correspond to negative beta values in the baseline PP < NP group comparison (i) match the baseline cluster-level results, (ii) match the PP vs. NP Z-score maps from Figure 3, (iii) are the regions with highest density of PGR and OXTR (rows "GCOR-PGR" and "GCOR-OXTR colocalization", and (iv) therefore influentce the spatial correlation pattern. Abbreviations: PP = postpartum, GLM = general linear model, NP = nulliparous, fALFF = fractional amplitude of low frequency fluctuations, LCOR = local correlation, GCOR = global correlation.

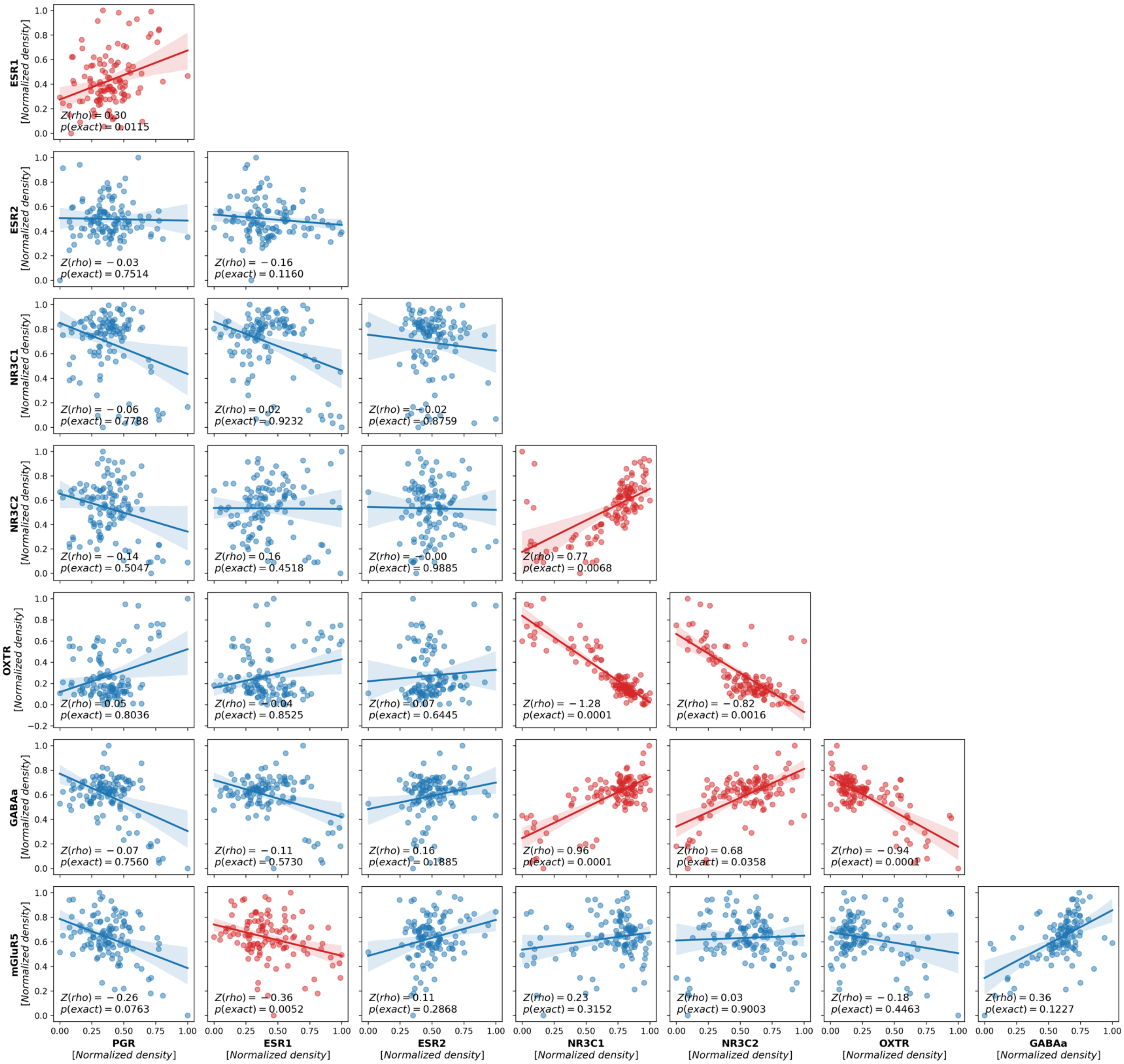


**Figure S6: Spatial correlation between mRNA expression and PET receptor maps.**

Axes: Receptor density of seven receptors, normalized to range 0–1. Scatters: Normalized receptor density for each of 116 parcels. Line: Linear regression fit with bootstrapped 95% confidence intervals. Statistics: Z-transformed Spearman's rho and exact p value calculated from a null distribution obtained from correlation with 10,000 spatial autocorrelation-preserving null maps. The p value for each pair is the average of two p values calculated from null distributions after permutation of either one map. Red: Colocalization p value < .05.
