## Supplementary material (analysis code notebook) for "Temporal dissociation between local and global functional adaptations of the maternal brain to childbirth: A longitudinal assessment": 3_results_analyses_plots.ipynb

```
# %% [markdown]
# PPRS: results, analyses, & plots

# %% [code]
import os
from os.path import join
from glob import glob
import pandas as pd
import numpy as np
from IPython import display
from nilearn.image import load_img, math_img, threshold_img
from nilearn.plotting import plot_design_matrix, plot_glass_brain, find_parcellation_cut_coords, plot_stat_map
from nilearn.glm.second_level import SecondLevelModel, non_parametric_inference
from nilearn.reporting import get_clusters_table
from nilearn.maskers import NiftiLabelsMasker
from sklearn.preprocessing import minmax_scale
import statsmodels.formula.api as smf
from scipy import stats
from pingouin import compute_effsize, ancova, pairwise_tests
from IPython.display import display
import matplotlib.pyplot as plt
import matplotlib as mpl
from matplotlib.cm import get_cmap
from matplotlib.gridspec import GridSpec, GridSpecFromSubplotSpec
from atlasreader import get_statmap_info
import seaborn as sns

from utils_image import index_clusters_bin

from abagen import get_expression_data
from nilearn.input_data import NiftiLabelsMasker
from utils_image import parcel_data_to_volume
import sys
sys.path.append(os.path.dirname("/Users/llotter/projects/juspyce/juspyce"))
from juspyce.api import JuSpyce
from juspyce.stats import corr, residuals, mc_correction

wd = "/Users/llotter/projects/pprs"

def na():
    return slice(None)

# %% [markdown]
## Data

# %% [markdown]
### Dataset

# %% [code]
tps = ["T0", "T1", "T2", "T3", "T4", "T5"]
    
# load processed dataset
data_all = pd.read_csv(join(wd, "pheno", "pprs_pheno_mri_proc.csv"))
data_all = data_all.set_index(["ID", "tp", "group_str"], drop=False)
data_all.index.names = ["idx_id", "idx_tp", "idx_group"]

# load additional test scores
data_scores = pd.read_csv(join(wd, "pheno", "pprs_pheno_mri_scores.csv"))
data_scores = data_scores.set_index(["id", "tp"])
data_scores.index.names = ["idx_id", "idx_tp"]

# merge
data_all = data_all.merge(data_scores, left_index=True, right_index=True, how="left", suffixes=["", "_DROP"])
data_all = data_all[[c for c in data_all.columns if not c.endswith("_DROP")]]

# print numbers
def print_numbers(data, tp):
    print(f"TP {tp}: PP = {len(data[(data.group_str=='PP') & (data.tp==tp)])}, "
          f"NP = {len(data[(data.group_str=='NP') & (data.tp==tp)])}")

print("Original dataset")
for tp in tps:
    print_numbers(data_all, tp)
    
# exclude sessions with reconstruction problem
data = data_all.copy()
data = data[~((data.ID=="sub-002PP") & (data.tp=="T4"))]
    
# apply motion cutoff
data = data[(data.fwd_mean <= 0.5) & (data.fwd_max <= 3)]

# print numbers
print("After motion cutoff")
for tp in tps:
    print_numbers(data, tp)
n = data.shape[0]
n_pp = len(data[data.group_str=="PP"])
n_np = len(data[data.group_str=="NP"])

# new variables
data["weeks_postpartal"] = data.days_since_delivery / 7
data["ep_ratio"] = data.estradiol / data.progesterone
for horm in ["estradiol", "progesterone", "ep_ratio"]:
    data[horm+"_log"] = np.log(data[horm])

# sort variables
data.contraception = pd.Categorical(data.contraception.replace({1:"yes", 0:"no"}))
data.baby_blues = pd.Categorical(data.baby_blues.replace({1:"yes", 0:"no"}))
data.breastfeeding = pd.Categorical(data.breastfeeding.replace({1:"yes", 0:"no"}))
data.n_children = data.n_children.astype(float).astype('Int64')
data = data.rename(columns={
    "nbs_falff_path":"falff_path",
    "nbs_gcor_path":"gcor_path",
    "nbs_lcor_path":"lcor_path"
})

# %% [markdown]
#### Subject lists

# %% [code]
subs_alltps = [sub for sub in data.ID.unique() if data.loc[(sub,),:].shape[0]==6]
subs_NP = data.loc[(na(),na(),"NP")].ID.unique().tolist()
subs_PP = data.loc[(na(),na(),"PP")].ID.unique().tolist()
subs_PP_long = [sub for sub in data.ID.unique() if data.loc[(sub,),:].shape[0]>1]
subs_PP_onlybl = [sub for sub in data.loc[(na(),na(),"PP")].ID.unique() if data.loc[(sub,),:].shape[0]==1]

# new variable with cross vs long subjects
data["group_crosslong_str"] = data.group_str.copy()
data.loc[(subs_PP_long, na(), "PP"), "group_crosslong_str"] = "PP-long"
data.loc[(subs_PP_onlybl, na(), "PP"), "group_crosslong_str"] = "PP-cross"

# %% [markdown]
### Templates

# %% [code]
# brain mask
mask_brain = load_img(join(wd, "templates", "maskbrain_mni152_3mm.nii"))

# gm mask
mask_gm = load_img(join(wd, "templates", "maskgm_mni152_3mm.nii"))

# gm probability
tpm_gm = load_img(join(wd, "templates", "gmprob_mni152_1mm.nii.gz"))

# parcellation
parc = load_img(join(wd, "templates", "parc_schaefer100-7-tianS1_3mm.nii"))
# parcellation info
parc_xyz, parc_idc = find_parcellation_cut_coords(parc, return_label_names=True)
parc_labels = pd.read_csv(join(wd, "templates", "parc_schaefer100-7-tianS1_3mm.txt"), header=None)[0].to_list()
parc_n = len(parc_idc)
parc_info = pd.DataFrame({
    "idx": parc_idc,
    "label": parc_labels,
    "x": parc_xyz[:, 0],
    "y": parc_xyz[:, 1],
    "z": parc_xyz[:, 2]
})

# analysis mask
mask = math_img("parc != 0", parc=parc)

# %% [markdown]
## Analyses: demographics

# %% [markdown]
### Cross-sectional: Continuous data

# %% [code]
vars = ["age", "edu", "fwd_mean", "fwd_max", "weeks_postpartal", "gestation_day", "baby_weight", "epds"]
vars_desc = ["count", "mean", "std", "min", "50%", "max"]

dem_data = []
for name, idc in zip(["Nulliparous (NP)", "Postpartum, all (PP)", "Postpartum, cross-sectional", "Postpartum, longitudinal"], 
                     [data.group_str=="NP", 
                      (data.group_str=="PP") & (data.tp=="T0"), 
                      data.group_crosslong_str=="PP-cross", 
                      (data.group_crosslong_str=="PP-long") & (data.tp=="T0")]):
    dem_data.append(
        data.loc[idc, vars].describe().T[vars_desc].rename(columns=dict(zip(vars_desc, [(name, v) for v in vars_desc]))).round(2)
    )
dem_data = pd.concat(dem_data, axis=1)

# analyses
for var in vars:
    for name, idc in zip(["MWU-test: NP vs. PP",
                          "MWU-test: PP-cross vs. PP-long"],
                         [[data.group_str=="NP", (data.group_str=="PP") & (data.tp=="T0")],
                          [data.group_crosslong_str=="PP-cross", (data.group_crosslong_str=="PP-long") & (data.tp=="T0")]]):
        try:
            mwu = stats.mannwhitneyu(
                x=data.loc[idc[0], var].dropna(),
                y=data.loc[idc[1], var].dropna(),
            )
            dem_data.loc[var, (name, "U")] = np.round(mwu[0], 2)
            dem_data.loc[var, (name, "p")] = np.round(mwu[1], 3)
        except:
            pass

dem_data.columns = pd.MultiIndex.from_tuples(dem_data.columns)
dem_data = pd.concat(
    [dem_data.loc[:, (c, na())] 
     for c in ["Nulliparous (NP)", "Postpartum, all (PP)", "MWU-test: NP vs. PP", 
               "Postpartum, cross-sectional", "Postpartum, longitudinal", "MWU-test: PP-cross vs. PP-long"]], 
    axis=1
)
dem_data = dem_data.rename(index={
    "age":"Age", 
    "edu":"Education", 
    "fwd_mean":"Mean FWD", 
    "fwd_max":"Max FWD",
    "weeks_postpartal":"Weeks postpartum",
    "gestation_day": "Gestation day",
    "baby_weight":"Weight of baby",
    "epds": "EPDS"
})

# save
dem_data.to_csv(join(wd, "tables", "demographics_cross1.csv"))
dem_data.to_html(join(wd, "tables", "demographics_cross1.html"))
dem_data

# %% [markdown]
### Cross-sectional: Discrete data

# %% [code]
vars = ["ethnicity", "baby_blues", "baby_sex_str", "birth_mode_str", "breastfeeding", "n_children", "contraception"]

dem_data_disc = []
for var in vars:
    for temp in [data.loc[(na(), "T0", na()), :].groupby(["group_str", var])[var].count(),
                 data.loc[(na(), "T0", "PP"), :].groupby(["group_crosslong_str", var])[var].count()]:
        dem_data_disc.append(
            temp \
                .to_frame() \
                .rename(columns={var: "Count"}) \
                .reset_index(drop=False) \
                .rename(columns={"group_str": "Group", "group_crosslong_str": "Group", var: "Category"}) \
                .assign(Variable=var) 
            )
dem_data_disc = pd.concat(dem_data_disc)
dem_data_disc = dem_data_disc.pivot(columns=["Group"], index=["Variable", "Category"])
dem_data_disc.loc[(["baby_blues", "breastfeeding"], na()), ("Count", "NP")] = np.nan
dem_data_disc.loc[("ethnicity", na()), :] = dem_data_disc.loc[("ethnicity", na()), :].replace({np.nan: 0})
dem_data_disc.loc[("ethnicity", na()), ("Count", "PP-long")] = np.nan
for var in ["baby_blues", "baby_sex_str", "birth_mode_str", "breastfeeding", "n_children"]:
    temp = dem_data_disc.loc[var, (na(), ["PP-cross", "PP-long"])]  
    # if var == "birth_mode_str":
    #     temp = pd.DataFrame({"section": temp.loc[["emergency", "section"], :].sum(),
    #                          "spontaneous": temp.loc[["spontaneous", "ventouse"], :].sum()}).T
    chi2 = stats.chi2_contingency(temp)
    dem_data_disc.loc[(var, na()), ("Chi2-test: PP-cross vs. PP-long", "Chi2")] = chi2[0].round(2)
    dem_data_disc.loc[(var, na()), ("Chi2-test: PP-cross vs. PP-long", "df")] = chi2[2]
    dem_data_disc.loc[(var, na()), ("Chi2-test: PP-cross vs. PP-long", "p")] = chi2[1].round(3)

temp = {
        "ethnicity": "Country of origin",
        "baby_blues": "Baby blues",
        "baby_sex_str": "Sex of baby",
        "birth_mode_str": "Birth mode",
        "breastfeeding": "Breastfeeding",
        "n_children": "Number of children",
        "Deutschland": "Germany",
        "deutschland": "Germany",
        "Bulgarien": "Bulgaria",
        "Italien": "Italy",
        "Schweiz": "Switzerland",
        "Belgien": "Belgium",
        "Niederlande": "Netherlands",
        "Polen": "Poland",
        "Russland": "Russia",
        "Frankreich": "France",
        "Slowakei": "Slovakia",
        "Spanien": "Spain",
        "Litauen": "Lithuania",
        "Tadschikistan": "Tajikistan",
        "f": "female",
        "m": "male",
        "NP": "Nulliparous (NP)", 
        "PP": "Postpartum (PP)",
        "PP-cross": "PP, cross-sectional",
        "PP-long": "PP, longitudinal",
        "contraception": "Contraception"
}
dem_data_disc.index = pd.MultiIndex.from_tuples(
    [(temp[v] if v in temp else v, temp[c] if c in temp else c) for v, c in dem_data_disc.index],
    names=dem_data_disc.index.names
)
dem_data_disc.columns = pd.MultiIndex.from_tuples(
    [(v, temp[c] if c in temp else c) for v, c in dem_data_disc.columns],
)

# save
dem_data_disc.to_csv(join(wd, "tables", "demographics_cross2.csv"))
dem_data_disc.to_html(join(wd, "tables", "demographics_cross2.html"))
dem_data_disc

# %% [markdown]
### Longitudinal: Continuous Data

# %% [code]
vars = ["weeks_postpartal", 
        "epds",  "mpas", "mpas_attachment", "mpas_hostility", "mpas_interaction",
        "estradiol", "estradiol_log", "progesterone", "progesterone_log", "ep_ratio", "ep_ratio_log",
        "fwd_mean", "fwd_max"]

# demographic & motion data
dem_data_long = {}
for var in vars:
    print(var)
    data_desc = data.loc[(subs_PP_long, na(), "PP"),:].groupby("idx_tp")[var].describe()[["count", "mean", "std"]]
    data_desc = pd.concat(
        [data_desc.loc[tp] for tp in tps], 
        axis="index", 
        keys=[f"Postpartum (PP): {tp}" for tp in tps]
    ).round(2)
    dem_data_long[var] = data_desc
    # calculate LMM 
    if var!= "weeks_postpartal":
        
        temp = data.loc[(subs_PP_long, na(), "PP"), [var, "weeks_postpartal", "ID"]].dropna()
        
        lmm = smf.mixedlm(f"{var} ~ weeks_postpartal", groups="ID", data=temp).fit()
        lmm2 = smf.mixedlm(f"{var} ~ weeks_postpartal + np.square(weeks_postpartal)", groups="ID", data=temp).fit()
        
        data_desc[("LMM", "df")] = f"{(lmm.df_modelwc, lmm.df_resid)}"
        data_desc[("Weeks PP", "converged")] = lmm.converged
        data_desc[("Weeks PP", "coef")] = lmm.params["weeks_postpartal"].round(3)
        data_desc[("Weeks PP", "p")] = lmm.pvalues["weeks_postpartal"].round(3)
        data_desc[("Weeks PP^2", "converged")] = lmm2.converged
        data_desc[("Weeks PP^2", "coef")] = lmm2.params["np.square(weeks_postpartal)"].round(3)
        data_desc[("Weeks PP^2", "p")] = lmm2.pvalues["np.square(weeks_postpartal)"].round(3)
dem_data_long = pd.DataFrame(dem_data_long) \
    .round(2) \
    .T
dem_data_long = pd.concat([
    dem_data_long.iloc[:,1:-6], 
    dem_data_long.iloc[:,0], 
    dem_data_long.iloc[:,-5], dem_data_long.iloc[:,-6], dem_data_long.iloc[:,-4],
    dem_data_long.iloc[:,-2], dem_data_long.iloc[:,-3], dem_data_long.iloc[:,-1]
], axis=1)

# rename variables  
dem_data_long = dem_data_long.rename(index={
    "age":"Age", 
    "mpas":"MPAS - total",
    "mpas_attachment": "MPAS - attachment", 
    "mpas_hostility": "MPAS - hostility", 
    "mpas_interaction": "MPAS - interaction",
    "epds":"EPDS",
    "weeks_postpartal":"Weeks postpartum",
    "estradiol": "Estradiol", 
    "estradiol_log": "log(Estradiol)",
    "progesterone": "Progesterone", 
    "progesterone_log": "log(Progesterone)", 
    "ep_ratio": "Estradiol/Progesterone", 
    "ep_ratio_log": "log(Estradiol/Progesterone)",
    "fwd_mean":"Mean FWD", 
    "fwd_max":"Max FWD",
})

# save
dem_data_long.to_csv(join(wd, "tables", "demographics_long1.csv"))
dem_data_long.to_html(join(wd, "tables", "demographics_long1.html"))
dem_data_long

# %% [markdown]
## 2nd level cross-sectional MRI analyses

# %% [markdown]
#### General analysis settings

# %% [code]
# rerun analyses?
rerun_sec_lev = False

# settings
vox_thresh = 0.01
cl_thresh = -np.log10(0.05)
cl_inference = "mass"
n_perm = 10000

# %% [markdown]
#### Inspect stored data paths

# %% [code]
for sub in ["sub-001PP", "sub-040PP"]:
    for c in ["swu_rest_path", "wu_rest_path", "falff_path", "lcor_path", "gcor_path"]:
        for tp in data.loc[(sub,na(),na()), "tp"]:
            temp = data.loc[(sub,tp,na()), c]
            print(sub, tp, c, temp.values[0])

# %% [markdown]
### Voxel-wise measures

# %% [markdown]
#### Prepare model

# %% [code]
sec_lev_desi = pd.DataFrame({
    "pp": (data.loc[(na(),"T0",na()),:].group_str == "PP").astype(int),
    "np": (data.loc[(na(),"T0",na()),:].group_str == "NP").astype(int),
    "age": data.loc[(na(),"T0",na()),:].age
})

print(f"N = {data.loc[(na(),'T0',na()),:].shape[0]} subjects")
plot_design_matrix(sec_lev_desi, rescale=False)

# %% [markdown]
#### Estimate

# %% [code]
def run_2nd_level(data, contrast_par, contrast_nonpar, design=sec_lev_desi, 
                  vox_thresh=vox_thresh, cl_thresh=cl_thresh, cl_inference=cl_inference,
                  save_dir=wd, save_prefix="",
                  mask=mask, n_perm=n_perm, n_jobs=8, seed=42,
                  plot=True, cmap="cold_white_hot", title=""):
    # load data
    data = [load_img(d) for d in data]

    # fit 2nd level model
    sec_lev = SecondLevelModel(
        mask_img=mask,
        smoothing_fwhm=None, 
        n_jobs=n_jobs).fit(
            second_level_input=data,
            design_matrix=design,
        )

    # parametric
    sec_lev_par = sec_lev.compute_contrast(
        second_level_contrast=contrast_par,
        output_type="all")
    
    # non-parametric
    sec_lev_nonpar = non_parametric_inference(
        second_level_input=data,
        design_matrix=design,
        second_level_contrast=contrast_nonpar,
        mask=mask,
        model_intercept=True,
        n_perm=n_perm,
        two_sided_test=False,
        smoothing_fwhm=None,
        n_jobs=n_jobs,
        random_state=seed,
        threshold=vox_thresh
    )
    
    # extract and save results maps
    maps = {}
    maps["map_par_effectsize"] = sec_lev_par["effect_size"]
    maps["map_par_stat"] = sec_lev_par["stat"]
    maps["map_par_logp"] = math_img("-np.log10(i)", i=sec_lev_par["p_value"])
    maps["map_nonpar_logp"] = sec_lev_nonpar[f"logp_max_{cl_inference}"]
    cl_sig = (maps["map_nonpar_logp"].get_fdata() > cl_thresh).any()
    if cl_sig:
        maps["map_nonpar_bin"] = math_img(f"i > {cl_thresh}", i=maps["map_nonpar_logp"])
        maps["map_nonpar_idx"], _, cl_sizes = index_clusters_bin(maps["map_nonpar_bin"])
        print(f"{cl_sizes.shape[0]} significant cluster(s) detected.")
    else:
        print("No significant clusters detected.")
    for m in maps:
        maps[m].to_filename(join(save_dir, f"{save_prefix}{m}.nii.gz"))
    
    # plot con1
    if plot:
        gb = plot_glass_brain(
            maps["map_par_logp"], 
            display_mode = "lzry",
            colorbar=True, 
            cmap=cmap,
            title=title
        )
        if cl_sig:
            gb.add_contours(
                img=maps["map_nonpar_logp"], 
                levels=[cl_thresh], 
                colors="k", 
            )
            gb.add_overlay(
                img=maps["map_nonpar_idx"],
                cmap="tab10",
                alpha=0.8
            )
        plt.show()
    
    # return
    return maps

# %% [code]
# iterate voxe-wise measures
sec_level_res = {}

for measure in ["falff", "lcor", "gcor"]:
    print(measure)
    sec_level_res[measure] = {}
    
    # RERUN
    if rerun_sec_lev:
        # get data files
        vols = [load_img(d) for d in data.loc[(na(),"T0",na()), measure + "_path"].to_list()]
        
        # con1
        print("con1")
        sec_level_res[measure]["con1"] = run_2nd_level(
            data=vols,
            contrast_par=np.array([1, -1, 0]),
            contrast_nonpar=np.array([1, 0, 0]),
            save_dir=join(wd, "results"),
            save_prefix=f"{measure}_con1_",
            cmap="cold_white_hot", 
            title=f"{measure}: PP > NP [-lg(p)]"
        )
        
        # con2
        print("con2")
        sec_level_res[measure]["con2"] = run_2nd_level(
            data=vols,
            contrast_par=np.array([-1, 1, 0]),
            contrast_nonpar=np.array([0, 1, 0]),
            save_dir=join(wd, "results"),
            save_prefix=f"{measure}_con2_",
            cmap="cold_white_hot_r", 
            title=f"{measure}: PP < NP [-lg(p)]"
        )
    
    else:
        # NOT RERUN, JUST LOAD SAVED FILES
        for con in ["con1", "con2"]:
            sec_level_res[measure][con] = {}
            for file in glob(join(wd, "results", f"{measure}_{con}_*.nii.gz")):
                name = os.path.basename(file).replace(f"{measure}_{con}_", "").replace(".nii.gz", "")
                sec_level_res[measure][con][name] = load_img(file)

# show results dict
sec_level_res

# %% [markdown]
## 2nd level longitudinal MRI analyses

# %% [code]
rerun_sec_lev_long = False

# %% [code]
sec_level_res_long = dict()

for measure in ["falff", "lcor", "gcor"]:
    print(measure)
    sec_level_res_long[measure] = dict()
    
    if rerun_sec_lev_long:
        
        for var in ["", "progesterone_log", "estradiol_log", "ep_ratio_log", 
                    "epds",  "mpas", "mpas_attachment", "mpas_hostility", "mpas_interaction"]:
            
                vars = ["age", "weeks_postpartal"]
                if var != "":
                    vars += [var]
                    
                sec_lev_desi_long = data.loc[(subs_PP_long, na(), "PP"), vars].copy().dropna()
                
                for sub in subs_PP_long:
                    sec_lev_desi_long[sub] = 0
                    for tp in sec_lev_desi_long.loc[(sub, na(), "PP"), :].index.get_level_values("idx_tp"):        
                        sec_lev_desi_long.loc[(sub, tp, "PP"), sub] = 1
                    if var != "":
                        temp = sec_lev_desi_long.loc[(sub, na(), "PP"), var]
                        if len(temp.unique()) == 1:
                            temp = temp - temp.mean()
                        else:
                            temp = (temp - temp.mean()) / temp.std()
                        if temp.isna().any():
                            print(f"na values for subject {sub}:", temp)
                        sec_lev_desi_long.loc[(sub, na(), "PP"), var] = temp
                        
                if var != "":
                    sec_lev_desi_long[var] = (sec_lev_desi_long[var] - sec_lev_desi_long[var].mean()) / sec_lev_desi_long[var].std()
                    sec_lev_desi_long[f"{var}*weeks_postpartal"] = sec_lev_desi_long[var] * sec_lev_desi_long["weeks_postpartal"]
                    contrast = f"{var}*weeks_postpartal"
                else:
                    contrast = "weeks_postpartal"
                print(contrast)
                # plot_design_matrix(sec_lev_desi_long, rescale=True)
                # plt.show()
        
                sec_lev = SecondLevelModel(
                    mask_img=mask,
                    smoothing_fwhm=None, 
                    n_jobs=-1
                ).fit(
                    second_level_input=[load_img(d) for d in data.loc[sec_lev_desi_long.index, f"{measure}_path"].to_list()],
                    design_matrix=sec_lev_desi_long,
                )
                    
                res = sec_lev.compute_contrast(
                    contrast, output_type="all"
                )
                
                sec_level_res_long[measure][contrast] = res["effect_size"]
                res["effect_size"].to_filename(join(wd, "results", f"{measure}_long_{contrast}.nii.gz"))

                plot_glass_brain(
                    res["effect_size"], 
                    display_mode="lzry",
                    colorbar=True, 
                    cmap="RdBu_r",
                    plot_abs=False,
                    symmetric_cbar=True,
                    title=f"{measure}: {contrast}"
                )
                plt.show()
        
    else:
        for file in glob(join(wd, "results", f"{measure}_long_*.nii.gz")):
            contrast = os.path.basename(file).replace(f"{measure}_long_", "").replace(".nii.gz", "")
            sec_level_res_long[measure][contrast] = load_img(file, wildcards=False)

# show results dict
sec_level_res_long

# %% [markdown]
## Post-hoc MRI analyses

# %% [markdown]
### Cluster statistics

# %% [code]
cl_stat = list()
for measure, measure_name in zip(["falff", "lcor", "gcor"], ["fALFF", "LCOR", "GCOR"]):
    for con, con_name in zip(["con1", "con2"], ["PP > NP", "PP < NP"]):
        
        if "map_nonpar_bin" in sec_level_res[measure][con].keys():
            map_thresh = sec_level_res[measure][con]["map_nonpar_logp"]
        
            cl_stat_stat = get_clusters_table(
                map_thresh,
                cl_thresh
            )
            cl_stat_atlas = get_statmap_info(
                map_thresh,
                cluster_extent=1,
                voxel_thresh=cl_thresh
            )[0]
            cl_stat.append(
                pd.concat([
                    cl_stat_stat[["X","Y","Z","Peak Stat","Cluster Size (mm3)"]], 
                    cl_stat_atlas["aal"]
                ], axis="columns") \
                    .assign(Measure=measure_name, Contrast=con_name) \
                    .set_index(["Measure", "Contrast"], drop=True)
            )

cl_stat = pd.concat(cl_stat, axis="index")
cl_stat["Peak Stat"] = cl_stat["Peak Stat"].astype("float").round(2)
cl_stat[["X", "Y", "Z"]] = cl_stat[["X", "Y", "Z"]].astype(int)
cl_stat = cl_stat.rename(columns={
    "Peak Stat": "-lg(p)",
    "Cluster Size (mm3)": "Cluster size [mm^3]",
    "aal": "AAL region coverage"
})

cl_names = [
    "fALFF-PP>NP-rightPrecentral", "fALFF-PP>NP-leftMedTemp",
    "fALFF-PP<NP-rightPFC",
    "LCOR-PP<NP-leftIns", "LCOR-PP<NP-rightIns",
    "GCOR-PP<NP-leftPut", "GCOR-PP<NP-rightPut", 
]
cl_stat["Cluster name"] = [name.split("-")[2] for name in cl_names]
cl_stat = cl_stat.set_index("Cluster name", append=True)

cl_stat.to_csv(join(wd, "tables", "cluster_stat.csv"))
cl_stat.to_html(join(wd, "tables", "cluster_stat.html"))
cl_stat

# %% [markdown]
### Data extraction

# %% [code]
rerun_cl_extract = False

# %% [code]
if rerun_cl_extract:
    # structural data
    vols_struct = [load_img(d) for d in data.gm_anat_path.to_list()]
    vbm_nona = ~data.vbm_path.isnull()
    vols_vbm = [load_img(d) for d in data.vbm_path[vbm_nona].to_list()]

    # VOXEL-WISE
    for measure in ["falff", "lcor", "gcor"]:
        print(measure)
        
        # image data
        vols_func = [load_img(d) for d in data[measure + "_path"].to_list()]
            
        for con in ["con1", "con2"]:
            
            # indexed thresholded image
            if "map_nonpar_idx" in sec_level_res[measure][con].keys():
                
                # masker
                masker = NiftiLabelsMasker(
                    labels_img=sec_level_res[measure][con]["map_nonpar_idx"],
                    standardize=False,
                )
                # extract functional and structural data
                temp_func = masker.fit_transform(vols_func)
                temp_struct = masker.fit_transform(vols_struct)
                temp_vbm = masker.fit_transform(vols_vbm)
                for cl in range(temp_func.shape[1]):
                    data[f"{measure}_{con}_cl{cl+1}"] = temp_func[:, cl]
                    data[f"gm_{measure}_{con}_cl{cl+1}"] = temp_struct[:, cl]    
                    data.loc[vbm_nona, f"vbm_{measure}_{con}_cl{cl+1}"] = temp_vbm[:, cl]    

    # SAVE    
    data.to_csv(join(wd, "pheno", "pprs_pheno_mri_proc_res.csv"))
    
else:
    data = pd.read_csv(join(wd, "pheno", "pprs_pheno_mri_proc_res.csv"))
    data = data.set_index(["idx_id", "idx_tp", "idx_group"])

# %% [markdown]
#### Cluster/Network labels

# %% [code]
cl_labs = [c for c in data.columns if c.startswith("falff_con") | c.startswith("lcor_con") | c.startswith("gcor_con")]
print(len(cl_labs), cl_labs)

cl_names = cl_names
print(len(cl_names), cl_names)

cl_iter = []
for lab, name in zip(cl_labs, cl_names):
    name = name.split("-")
    cl_iter.append((
        lab, 
        name[0], 
        name[1].replace(">", " > ").replace("<", " < "), 
        "" if lab.startswith("nbs") else name[2] 
    ))
print(len(cl_iter), cl_iter)

# %% [markdown]
### MRI baseline sensitivity analyses

# %% [markdown]
#### Covariates PP vs NP comparison

# %% [code]
mri_bl_confounds = list()

for lab, measure, contrast, name in cl_iter:
            
    # get MRI data
    temp_mri = data.loc[(na(), "T0", na()), lab]

    # initiate save df
    temp_df = pd.DataFrame({
        "Measure": measure,
        "Contrast": contrast,
        "Cluster name": name
    }, index=range(1))
    
    # iterate confounds
    for confound, confound_name in zip(
        [["age"], ["age", "fwd_mean", "fwd_max"], ["age", f"gm_{lab}"], ["age", "edu"], ],
        ["Age", "Age, FWD", "Age, GM", "Age, Education"],
    ):
        
        temp_conf = data.loc[(na(), "T0", na()), confound]
        temp_na = ~temp_conf.isna().any(1).values
        # regress confound from data
        temp_resid = residuals(x=temp_conf[temp_na].values, y=temp_mri[temp_na].values)       
        
        # calc statistics
        aov = ancova(
            data=data.loc[(na(), "T0", na()),:][temp_na],
            dv=lab,
            between="group",
            covar=confound
        )
        f, p, np2 = [aov.loc[aov.Source=="group", c].values[0] for c in ["F", "p-unc", "np2"]]
        df = f"({aov.DF[0]}, {aov.DF.values[-1]})"
        
        g = compute_effsize(
            x=temp_resid[data[data.tp=="T0"].loc[temp_na].group_str == "PP"],
            y=temp_resid[data[data.tp=="T0"].loc[temp_na].group_str == "NP"],
            paired=False,
            eftype="hedges"
        )

        # save
        temp_df[("ANCOVA: PP vs. NP: " + confound_name, "F")] = f.round(2)
        temp_df[("ANCOVA: PP vs. NP: " + confound_name, "df")] = df
        temp_df[("ANCOVA: PP vs. NP: " + confound_name, "-lg(p)")] = -np.log10(p).round(3) #p.round(3) if p >= 0.001 else "< 0.001"
        temp_df[("ANCOVA: PP vs. NP: " + confound_name, "np2")] = np2.round(2)
        temp_df[("ANCOVA: PP vs. NP: " + confound_name, "g")] = g.round(2)
    
    # save
    temp_df = temp_df.set_index(["Measure", "Contrast", "Cluster name"])
    mri_bl_confounds.append(temp_df)

mri_bl_confounds = pd.concat(mri_bl_confounds)
mri_bl_confounds.columns = pd.MultiIndex.from_tuples(mri_bl_confounds.columns)

mri_bl_confounds.to_csv(join(wd, "tables", "mri_baseline_conf.csv"))
mri_bl_confounds.to_html(join(wd, "tables", "mri_baseline_conf.html"))
mri_bl_confounds

# %% [markdown]
#### Covariates within-PP

# %% [code]
vars = ["birth_mode", "baby_sex", "n_children", "age", "weeks_postpartal", "baby_weight", "gestation_day", "epds", "fwd_mean", "fwd_max"]
var_names = ["Birth mode", "Sex of baby", "Number of children", "Age", "Postpartum week", "Weight of baby", "Gestation day", "EPDS", "Mean FWD", "Max FWD"]

mri_bl_assoc = pd.DataFrame(
    index=pd.MultiIndex.from_tuples(cl_iter, names=["lab", "Measure", "Contrast", "Cluster name"])
)

for var, var_name in zip(vars, var_names):
    for lab, measure, contrast, name in cl_iter:
        
        temp = data.loc[(na(), "T0", "PP"), [var, lab] + (["age"] if var!="age" else [])].dropna()
        temp = temp.sort_values(by=var)
        
        if var in ["birth_mode", "baby_sex", "n_children"]:
            aov = ancova(
                data=temp,
                dv=lab,
                between=var,
                covar="age"
            )
            f, p, np2 = [aov.loc[aov.Source==var, c].values[0] for c in ["F", "p-unc", "np2"]]
            df = f"({aov.DF[0]}, {aov.DF.values[-1]})"
            mri_bl_assoc.loc[(lab, measure, contrast, name), [(f"ANCOVA: {var_name} - {s}") for s in ["F", "df", "p", "np2"]]] = \
                (f.round(2), df, p.round(3), np2.round(2))
                
        else:
            temp[var] = (temp[var] - temp[var].mean()) / temp[var].std()
            lr = smf.ols(f"{lab} ~ {var}{' + age' if var!='age' else ''}", data=temp).fit()
            coef, df, p = lr.params[var], f"({lr.df_model:.0f}, {lr.df_resid:.0f})", lr.pvalues[var]
            mri_bl_assoc.loc[(lab, measure, contrast, name), [(f"MLR: {var_name} - {s}") for s in ["coef", "df", "p"]]] = \
                (coef.round(3), df, p.round(3))
            
            
mri_bl_assoc.columns = pd.MultiIndex.from_tuples([tuple(c.split(" - ")) for c in mri_bl_assoc.columns])
mri_bl_assoc = mri_bl_assoc.droplevel("lab")

mri_bl_assoc.to_csv(join(wd, "tables", "mri_baseline_assoc.csv"))
mri_bl_assoc.to_html(join(wd, "tables", "mri_baseline_assoc.html"))
mri_bl_assoc     

# %% [markdown]
#### Spatial correlation between metrics and with longitudinal maps

# %% [code]
rerun_spc_longcross = False

# %% [code]
if rerun_spc_longcross:
    measures = ["falff", "lcor", "gcor"]
    method = "spearman"
    
    temp = [sec_level_res[m]["con1"]["map_par_effectsize"] for m in measures] + [sec_level_res_long[m]["weeks_postpartal"] for m in measures]
    temp_lab = [m+"_cross" for m in measures] + [m+"_long" for m in measures]
    
    jsp = JuSpyce(
        y=temp,
        y_labels=temp_lab,
        x=temp,
        x_labels=temp_lab,
        z=[tpm_gm],
        data_space="mni152",
        parcellation=parc,
        parcellation_space="mni152",
        parcellation_labels=parc_info.label.to_list(),
        standardize="xy",
        n_proc=-1,
    ).fit()
    jsp.transform("partial", dataset="X", replace=True, verbose=True)
    jsp.transform("partial", dataset="Y", replace=True, verbose=True)
    
    jsp.predict(method)
    jsp.permute_maps(method, permute="x", n_perm=10000, seed=42)
    mri_longcross_cor = pd.concat([
        jsp.predictions[method],
        jsp.p_predictions[method]
    ], axis=1)
    mri_longcross_cor.columns = pd.MultiIndex.from_tuples([(c, "rho") for c in jsp.x_lab] + [(c, "p") for c in jsp.x_lab])

    # save
    mri_longcross_cor.to_csv(join(wd, "results", "spc_longcross.csv"))
    jsp.X.to_csv(join(wd, "results", "spc_longcross_data.csv"))
    
else:
    mri_longcross_cor = pd.read_csv(join(wd, "results", "spc_longcross.csv"), index_col=0, header=[0,1])

# show
mri_longcross_cor

# %% [markdown]
## Longitudinal MRI analyses

# %% [markdown]
#### Cross: PP vs NP each timepoint

# %% [code]
mri_long_unpaired = list()
p_list = list()

for tp in tps:
    
    subs_PP_tp = data.loc[(subs_PP_long, tp, "PP")].ID.unique().tolist()
    
    temp_df2 = list()
    for lab, measure, contrast, name in cl_iter:
            
        temp = pd.concat([
            data.loc[(subs_PP_tp, tp, "PP"), :],
            data.loc[(subs_NP, "T0", "NP"), :]
        ])
            
        # calc statistics
        aov = ancova(
            data=temp,
            dv=lab,
            between="group",
            covar="age"
        )
        f, p, np2 = [aov.loc[aov.Source=="group", c].values[0] for c in ["F", "p-unc", "np2"]]
        df = f"({aov.DF[0]}, {aov.DF.values[-1]})"
        p_list.append(p)
        
        temp_resid = residuals(
            temp.age, 
            temp[lab]
        )
        
        g = compute_effsize(
            x=temp_resid[temp.group_str == "PP"],
            y=temp_resid[temp.group_str == "NP"],
            paired=False,
            eftype="hedges"
        )
        
        # initiate save df
        temp_df = pd.DataFrame({
            "Measure": measure,
            "Contrast": contrast,
            "Cluster name": name,
            (f"ANCOVA: PP-{tp} vs. NP: Age", "F"): f.round(2),
            (f"ANCOVA: PP-{tp} vs. NP: Age", "df"): df,
            (f"ANCOVA: PP-{tp} vs. NP: Age", "-lg(p)"): -np.log10(p).round(3), 
            (f"ANCOVA: PP-{tp} vs. NP: Age", "np2"): np2.round(2),
            (f"ANCOVA: PP-{tp} vs. NP: Age", "g"): g.round(2)
        }, index=range(1))
        
        # save
        temp_df = temp_df.set_index(["Measure", "Contrast", "Cluster name"])
        temp_df2.append(temp_df)
        
    temp_df2 = pd.concat(temp_df2)
    mri_long_unpaired.append(temp_df2)

# p correction
q_list = -np.log10(mc_correction(p_list)[0]).reshape(len(cl_names), len(tps), order="F").round(3)
for i, tp in enumerate(tps):
     mri_long_unpaired[i][(f"ANCOVA: PP-{tp} vs. NP: Age", "-lg(q)")] = q_list[:,i]
     mri_long_unpaired[i] = mri_long_unpaired[i].iloc[:,[0,1,2,5,3,4]]

mri_long_unpaired = pd.concat(mri_long_unpaired, axis="columns")
mri_long_unpaired.columns = pd.MultiIndex.from_tuples(mri_long_unpaired.columns)

mri_long_unpaired.to_csv(join(wd, "tables", "mri_longitudinal_unpaired_stat.csv"))
mri_long_unpaired.to_html(join(wd, "tables", "mri_longitudinal_unpaired_stat.html"))
mri_long_unpaired

# %% [markdown]
### Cross: PP vs NP each timepoint controlled for GM

# %% [code]
mri_long_unpaired_gm = list()
p_list = list()

for tp in tps:
    
    subs_PP_tp = data.loc[(subs_PP_long, tp, "PP")].ID.unique().tolist()
    
    temp_df2 = list()
    for lab, measure, contrast, name in cl_iter:
            
        temp = pd.concat([
            data.loc[(subs_PP_tp, tp, "PP"), :],
            data.loc[(subs_NP, "T0", "NP"), :]
        ])
            
        # calc statistics
        aov = ancova(
            data=temp,
            dv=lab,
            between="group",
            covar=["age", "gm_"+lab]
        )
        f, p, np2 = [aov.loc[aov.Source=="group", c].values[0] for c in ["F", "p-unc", "np2"]]
        df = f"({aov.DF[0]}, {aov.DF.values[-1]})"
        p_list.append(p)
        
        temp_resid = residuals(
            temp[["age", "gm_"+lab]], 
            temp[lab]
        )
        
        g = compute_effsize(
            x=temp_resid[temp.group_str == "PP"],
            y=temp_resid[temp.group_str == "NP"],
            paired=False,
            eftype="hedges"
        )
        
        # initiate save df
        temp_df = pd.DataFrame({
            "Measure": measure,
            "Contrast": contrast,
            "Cluster name": name,
            (f"ANCOVA: PP-{tp} vs. NP: Age", "F"): f.round(2),
            (f"ANCOVA: PP-{tp} vs. NP: Age", "df"): df,
            (f"ANCOVA: PP-{tp} vs. NP: Age", "-lg(p)"): -np.log10(p).round(3), 
            (f"ANCOVA: PP-{tp} vs. NP: Age", "np2"): np2.round(2),
            (f"ANCOVA: PP-{tp} vs. NP: Age", "g"): g.round(2)
        }, index=range(1))
        
        # save
        temp_df = temp_df.set_index(["Measure", "Contrast", "Cluster name"])
        temp_df2.append(temp_df)
        
    temp_df2 = pd.concat(temp_df2)
    mri_long_unpaired_gm.append(temp_df2)

# p correction
q_list = -np.log10(mc_correction(p_list)[0]).reshape(len(cl_names), len(tps), order="F").round(3)
for i, tp in enumerate(tps):
     mri_long_unpaired_gm[i][(f"ANCOVA: PP-{tp} vs. NP: Age", "-lg(q)")] = q_list[:,i]
     mri_long_unpaired_gm[i] = mri_long_unpaired_gm[i].iloc[:,[0,1,2,5,3,4]]

mri_long_unpaired_gm = pd.concat(mri_long_unpaired_gm, axis="columns")
mri_long_unpaired_gm.columns = pd.MultiIndex.from_tuples(mri_long_unpaired_gm.columns)

mri_long_unpaired_gm.to_csv(join(wd, "tables", "mri_longitudinal_unpaired_gm_stat.csv"))
mri_long_unpaired_gm.to_html(join(wd, "tables", "mri_longitudinal_unpaired_gm_stat.html"))
mri_long_unpaired_gm

# %% [markdown]
### Long: LMM

# %% [code]
mri_long_lmm = list()
p_values = list()

for lab, measure, contrast, name in cl_iter:
    print(lab)
    
    # longitudinal data, only PP, doesnt matter if all timepoints
    temp_mri_long = data.loc[(subs_PP_long, na(), "PP"), :]
    
    # calculate LMM and pairwise paired t tests
    lmm = smf.mixedlm(f"{lab} ~ weeks_postpartal", data=temp_mri_long, groups="ID").fit()
    lmm2 = smf.mixedlm(f"{lab} ~ weeks_postpartal + np.square(weeks_postpartal)", data=temp_mri_long, groups="ID").fit()
    p_values.append(lmm.pvalues[["weeks_postpartal"]].to_list() + lmm2.pvalues[["weeks_postpartal"]].to_list())

    temp_df = pd.DataFrame({
        "Measure": measure,
        "Contrast": contrast,
        "Cluster name": name,
        ("", "df"): f"({lmm.df_modelwc}, {lmm.df_resid})",
        ("Weeks PP", "converged"): lmm.converged,
        ("Weeks PP", "coef"): lmm.params["weeks_postpartal"].round(4),
        ("Weeks PP", "-lg(p)"): -np.log10(lmm.pvalues["weeks_postpartal"]).round(3),
        ("Weeks PP^2", "converged"): lmm2.converged,
        ("Weeks PP^2", "coef"): lmm2.params["np.square(weeks_postpartal)"].round(4),
        ("Weeks PP^2", "-lg(p)"): -np.log10(lmm2.pvalues["np.square(weeks_postpartal)"]).round(3),
    }, index=range(1))
    
    # save
    temp_df = temp_df.set_index(["Measure", "Contrast", "Cluster name"])
    mri_long_lmm.append(temp_df)
        
mri_long_lmm = pd.concat(mri_long_lmm)
mri_long_lmm.columns = pd.MultiIndex.from_tuples(mri_long_lmm.columns)

p_values = np.array(p_values).flatten()
q_values = mc_correction(p_values)[0]
mri_long_lmm["Weeks PP", "-lg(q)"] = -np.log10(q_values[::2]).round(3)
mri_long_lmm["Weeks PP^2", "-lg(q)"] = -np.log10(q_values[1::2]).round(3)

mri_long_lmm = mri_long_lmm.iloc[:, [0, 1,2,3,-2, 4,5,6,-1]]

mri_long_lmm.to_csv(join(wd, "tables", "mri_longitudinal_lmm_stat.csv"))
mri_long_lmm.to_html(join(wd, "tables", "mri_longitudinal_lmm_stat.html"))
mri_long_lmm

# %% [markdown]
### Long: LMM controlled for GM

# %% [code]
mri_long_lmm_gm = list()
p_values = list()

for lab, measure, contrast, name in cl_iter:
    print(lab)
    
    # longitudinal data, only PP, doesnt matter if all timepoints
    temp_mri_long = data.loc[(subs_PP_long, na(), "PP"), :]
    
    # calculate LMM and pairwise paired t tests
    lmm = smf.mixedlm(f"{lab} ~ weeks_postpartal + gm_{lab}", data=temp_mri_long, groups="ID").fit()
    lmm2 = smf.mixedlm(f"{lab} ~ weeks_postpartal + np.square(weeks_postpartal) + gm_{lab}", data=temp_mri_long, groups="ID").fit()
    p_values.append(lmm.pvalues[["weeks_postpartal"]].to_list() + lmm2.pvalues[["weeks_postpartal"]].to_list())

    temp_df = pd.DataFrame({
        "Measure": measure,
        "Contrast": contrast,
        "Cluster name": name,
        ("", "df"): f"({lmm.df_modelwc}, {lmm.df_resid})",
        ("Weeks PP", "converged"): lmm.converged,
        ("Weeks PP", "coef"): lmm.params["weeks_postpartal"].round(4),
        ("Weeks PP", "-lg(p)"): -np.log10(lmm.pvalues["weeks_postpartal"]).round(3),
        ("Weeks PP^2", "converged"): lmm2.converged,
        ("Weeks PP^2", "coef"): lmm2.params["np.square(weeks_postpartal)"].round(4),
        ("Weeks PP^2", "-lg(p)"): -np.log10(lmm2.pvalues["np.square(weeks_postpartal)"]).round(3),
    }, index=range(1))
    
    # save
    temp_df = temp_df.set_index(["Measure", "Contrast", "Cluster name"])
    mri_long_lmm_gm.append(temp_df)
        
mri_long_lmm_gm = pd.concat(mri_long_lmm_gm)
mri_long_lmm_gm.columns = pd.MultiIndex.from_tuples(mri_long_lmm_gm.columns)

p_values = np.array(p_values).flatten()
q_values = -np.log10(mc_correction(p_values)[0])
mri_long_lmm_gm["Weeks PP", "-lg(q)"] = q_values[::2].round(3)
mri_long_lmm_gm["Weeks PP^2", "-lg(q)"] = q_values[1::2].round(3)

mri_long_lmm_gm = mri_long_lmm_gm.iloc[:, [0, 1,2,3,-2, 4,5,6,-1]]

mri_long_lmm_gm.to_csv(join(wd, "tables", "mri_longitudinal_lmm_gm_stat.csv"))
mri_long_lmm_gm.to_html(join(wd, "tables", "mri_longitudinal_lmm_gm_stat.html"))
mri_long_lmm_gm

# %% [markdown]
### Long: LMM controlled for other covariates

Only sign results (-> GCOR in putamen)

# %% [code]
mri_long_lmm_cov = list()
mri_long_sig = (mri_long_lmm.loc[:, (na(), "-lg(q)")] > -np.log10(0.05)).any(axis=1)

for cov, cov_name, tps_cov in zip([None, "age", "fwd_mean + fwd_max", "weeks_postpartal", "gestation_day", "birth_mode", "baby_weight", "baby_sex", "breastfeeding", "n_children"],
                                  ["None", "Age", "FWD", "Postpartum week (BL)", "Gestation day", "Birth mode", "Weight of baby", "Sex of baby", "Breastfeeding", "Number of children"],
                                  [tps, tps, tps, tps, tps, tps, tps, tps, ["T0", "T4", "T5"], tps, tps, tps,]):
    print(cov)  
    # longitudinal data, only PP, doesnt matter if all timepoints
    temp_mri_long = data.loc[(subs_PP_long, tps_cov, "PP"), :].copy()
    if cov == "weeks_postpartal":
        temp_mri_long["weeks_postpartal_bl"] = np.nan
        for sub in subs_PP_long:
            temp_mri_long.loc[(sub, na(), "PP"), "weeks_postpartal_bl"] = temp_mri_long.loc[(sub, "T0", "PP"), "weeks_postpartal"]
        cov = "weeks_postpartal_bl"
          
    for lab, measure, contrast, name in np.array(cl_iter)[mri_long_sig]:
        
        # calculate LMM and pairwise paired t tests
        lmm = smf.mixedlm(f"{lab} ~ weeks_postpartal {'+ ' + cov if cov is not None else ''}", data=temp_mri_long, groups="ID").fit()
        lmm2 = smf.mixedlm(f"{lab} ~ weeks_postpartal + np.square(weeks_postpartal) {'+ ' + cov if cov is not None else ''}", data=temp_mri_long, groups="ID").fit()

        temp_df = pd.DataFrame({
            "Covariate": cov_name,
            "Measure": measure,
            "Contrast": contrast,
            "Cluster name": name,
            ("", "df"): f"({lmm.df_modelwc}, {lmm.df_resid})",
            ("Weeks PP", "converged"): lmm.converged,
            ("Weeks PP", "coef"): lmm.params["weeks_postpartal"].round(4),
            ("Weeks PP", "-lg(p)"): -np.log10(lmm.pvalues["weeks_postpartal"]).round(3),
            ("Weeks PP^2", "converged"): lmm2.converged,
            ("Weeks PP^2", "coef"): lmm2.params["np.square(weeks_postpartal)"].round(4),
            ("Weeks PP^2", "-lg(p)"): -np.log10(lmm2.pvalues["np.square(weeks_postpartal)"]).round(3),
        }, index=range(1))
        
        # save
        temp_df = temp_df.set_index(["Covariate", "Measure", "Contrast", "Cluster name"])
        mri_long_lmm_cov.append(temp_df)
        
mri_long_lmm_cov = pd.concat(mri_long_lmm_cov)
mri_long_lmm_cov.columns = pd.MultiIndex.from_tuples(mri_long_lmm_cov.columns)

mri_long_lmm_cov.to_csv(join(wd, "tables", "mri_longitudinal_lmm_cov_stat.csv"))
mri_long_lmm_cov.to_html(join(wd, "tables", "mri_longitudinal_lmm_cov_stat.html"))
mri_long_lmm_cov

# %% [markdown]
### Long: PP vs PP each timepoint

# %% [code]
mri_long_paired = list()
p_list = list()

for lab, measure, contrast, name in cl_iter:
    print(lab)
    
    # longitudinal data, only PP, doesnt matter if all timepoints
    temp_mri_long = data.loc[(subs_PP_long, na(), "PP"), :].copy()
    
    temp_df = pd.DataFrame({
        "Measure": measure,
        "Contrast": contrast,
        "Cluster name": name,
    }, index=range(1))
    
    ph = pairwise_tests(
        data=temp_mri_long,
        dv=lab,
        within="tp",
        subject="ID",
        padjust="fdr_bh",
        #alternative="less" if contrast=="PP < NP" else "greater"
    )
                
    # calculate single T-Tests PP vs PP
    # iterate timepoints
    for tp in ph.A.unique():
        for tp_comp in ph.loc[ph.A==tp, "B"].unique():
            
            temp_df[(f"T-test: {tp} vs. {tp_comp}", "T")] = ph.loc[(ph.A==tp) & (ph.B==tp_comp), "T"].values[0].round(2)
            temp_df[(f"T-test: {tp} vs. {tp_comp}", "-lg(p)")] = -np.log10(ph.loc[(ph.A==tp) & (ph.B==tp_comp), "p-unc"].values[0]).round(3)
            temp_df[(f"T-test: {tp} vs. {tp_comp}", "-lg(q)")] = -np.log10(ph.loc[(ph.A==tp) & (ph.B==tp_comp), "p-corr"].values[0]).round(3)
            temp_df[(f"T-test: {tp} vs. {tp_comp}", "g")] = ph.loc[(ph.A==tp) & (ph.B==tp_comp), "hedges"].values[0].round(2)
            
    # save
    temp_df = temp_df.set_index(["Measure", "Contrast", "Cluster name"])
    mri_long_paired.append(temp_df)
        
mri_long_paired = pd.concat(mri_long_paired)
mri_long_paired.columns = pd.MultiIndex.from_tuples(mri_long_paired.columns)

mri_long_paired.to_csv(join(wd, "tables", "mri_longitudinal_paired_stat.csv"))
mri_long_paired.to_html(join(wd, "tables", "mri_longitudinal_paired_stat.html"))
mri_long_paired

# %% [markdown]
### Long: PP vs PP each timepoint controlled for GM

# %% [code]
mri_long_paired_gm = list()
p_list = list()

for lab, measure, contrast, name in cl_iter:
    print(lab)
    
    # longitudinal data, only PP, doesnt matter if all timepoints
    temp_mri_long = data.loc[(subs_PP_long, na(), "PP"), :].copy()
    
    # regress gm
    temp_mri_long[lab] = residuals(temp_mri_long["gm_"+lab], temp_mri_long[lab])
    
    temp_df = pd.DataFrame({
        "Measure": measure,
        "Contrast": contrast,
        "Cluster name": name,
    }, index=range(1))
    
    ph = pairwise_tests(
        data=temp_mri_long,
        dv=lab,
        within="tp",
        subject="ID",
        padjust="fdr_bh",
        #alternative="less" if contrast=="PP < NP" else "greater"
    )
                
    # calculate single T-Tests PP vs PP
    # iterate timepoints
    for tp in ph.A.unique():
        for tp_comp in ph.loc[ph.A==tp, "B"].unique():
            
            temp_df[(f"T-test: {tp} vs. {tp_comp}", "T")] = ph.loc[(ph.A==tp) & (ph.B==tp_comp), "T"].values[0].round(2)
            temp_df[(f"T-test: {tp} vs. {tp_comp}", "-lg(p)")] = -np.log10(ph.loc[(ph.A==tp) & (ph.B==tp_comp), "p-unc"].values[0]).round(3)
            temp_df[(f"T-test: {tp} vs. {tp_comp}", "-lg(q)")] = -np.log10(ph.loc[(ph.A==tp) & (ph.B==tp_comp), "p-corr"].values[0]).round(3)
            temp_df[(f"T-test: {tp} vs. {tp_comp}", "g")] = ph.loc[(ph.A==tp) & (ph.B==tp_comp), "hedges"].values[0].round(2)
            
    # save
    temp_df = temp_df.set_index(["Measure", "Contrast", "Cluster name"])
    mri_long_paired_gm.append(temp_df)
        
mri_long_paired_gm = pd.concat(mri_long_paired_gm)
mri_long_paired_gm.columns = pd.MultiIndex.from_tuples(mri_long_paired_gm.columns)

mri_long_paired_gm.to_csv(join(wd, "tables", "mri_longitudinal_paired_gm_stat.csv"))
mri_long_paired_gm.to_html(join(wd, "tables", "mri_longitudinal_paired_gm_stat.html"))
mri_long_paired_gm

# %% [markdown]
## Spatial correlation analyses

# %% [markdown]
### Get receptor data

# %% [markdown]
#### Get ABA data

# %% [code]
reload_aba = False

# %% [code]
if reload_aba:
    # abagen parcellation info
    parc_info_aba = parc_info[["idx", "label"]].rename(columns={"idx":"id"})
    parc_info_aba["hemisphere"] = [l.split("_")[0][0] for l in parc_info_aba.label]
    parc_info_aba["structure"] = ["cortex"]*100 + ["subcortex/brainstem"]*16

    # get ABA data using abagen
    aba_data, aba_report = get_expression_data(
        atlas=parc, atlas_info=parc_info_aba, n_proc=6, lr_mirror="bidirectional", 
        return_donors=False, sim_threshold=None, ibf_threshold=0,
        return_report=True)
    print(aba_data.shape)   
    
    # save report
    with open(join(wd, "templates", "aba_report.md"), "w") as f:
        f.write(aba_report)
    
    # get progesterone data
    pred_aba = aba_data[["PGR", "ESR1", "ESR2", "NR3C1", "NR3C2", "OXTR"]]
    pred_aba.index = pd.Index(parc_labels, name="labels")
    pred_aba.to_csv(join(wd, "templates", "aba_genes.csv"))

else:
    pred_aba = pd.read_csv(join(wd, "templates", "aba_genes.csv"), index_col=0)   
    
pred_aba

# %% [markdown]
#### Get PET data

# %% [code]
parcellater = NiftiLabelsMasker(parc, parc_labels)
pred_pet = pd.DataFrame({
    "GABAa": parcellater.fit_transform(join(wd, "templates", "GABAa-flumazenil-10-kaulen2022.nii.gz"))[0],
    "mGluR5": parcellater.fit_transform(join(wd, "templates", "mGluR5-abp688-73-smart2019.nii.gz"))[0], 
}, index=pd.Index(parc_labels, name="labels"))

# %% [markdown]
#### Merge

# %% [code]
pred_aba_pet = pd.concat([pred_aba, pred_pet], axis=1)
pred_aba_pet.to_csv(join(wd, "templates", "spc_data.csv"))

# %% [markdown]
#### Spatial correlation between receptors

# %% [code]
rerun_spc_receptor = False

# %% [code]
if rerun_spc_receptor:
    method = "spearman"
    
    jsp = JuSpyce(
        y=pred_aba_pet.T,
        x=pred_aba_pet.T,
        z=[tpm_gm],
        data_space="mni152",
        parcellation=parc,
        parcellation_space="mni152",
        parcellation_labels=parc_info.label.to_list(),
        standardize="xy",
        n_proc=-1,
    ).fit()
    jsp.transform("partial", dataset="X", replace=True, verbose=True)
    jsp.transform("partial", dataset="Y", replace=True, verbose=True)
    
    jsp.predict(method)
    jsp.permute_maps(method, permute="x", n_perm=10000, seed=42)
    pred_spc = pd.concat([
        jsp.predictions[method],
        jsp.p_predictions[method]
    ], axis=1)
    pred_spc.columns = pd.MultiIndex.from_tuples([(c, "rho") for c in jsp.x_lab] + [(c, "p") for c in jsp.x_lab])

    # save
    pred_spc.to_csv(join(wd, "results", "spc_receptors.csv"))
    
else:
    pred_spc = pd.read_csv(join(wd, "results", "spc_receptors.csv"), index_col=0, header=[0,1])

# show
pred_spc

# %% [markdown]
### Run spatial correlations

# %% [code]
rerun_spc = False

# %% [code]
metrics = ["fALFF", "LCOR", "GCOR"]

if rerun_spc:
    
    # run
    res_jsp = dict()
    for m in metrics:
        res_jsp[m] = dict()
        print(m)

        jsp = JuSpyce(
            x=pred_aba_pet.T,
            y=data[m.lower() + "_path"].to_list(),
            z=[tpm_gm],
            data_space="mni152",
            parcellation=parc,
            parcellation_space="mni152",
            parcellation_labels=parc_info.label.to_list(),
            y_labels=data.index,
            standardize="x",
            drop_nan=False,
            n_proc=8
        ).fit(verbose=False)
        jsp.transform("partial", dataset="X", replace=True, verbose=False)
        jsp.transform("partial", dataset="Y", replace=True, verbose=False)
            
        # save data
        jsp.Y.to_csv(join(wd, "results", f"spc_pp_data_{m}.csv"))

        for tp in tps:
            res_jsp[m][tp] = dict()
            print(tp)
            
            groups = (data.group.replace({0:2}) * ((data.tp == tp) | (data.group_str == "NP"))).replace({0:np.nan})
            jsp.compare(comparison="z(A,B)", groups=groups, verbose=False)
            jsp.predict(comparison="z(A,B)", method="spearman", r_to_z=True, verbose=False)
            jsp.permute_groups(
                comparison="z(A,B)",
                groups=groups,
                method="spearman",
                p_from_average_y="mean",
                r_to_z=True,
                n_perm=10000,
                seed=42,
                verbose=True
            )
            res_jsp[m][tp]["zr"] = jsp.predictions["z(A,B)-spearman"]
            res_jsp[m][tp]["zrp"] = jsp.p_comparisons["z(A,B)-spearman-norm"]
            res_jsp[m][tp]["zrp_exact"] = jsp.p_comparisons["z(A,B)-spearman"]
            jsp.nulls = dict()
            
    # collect results
    spc_zr, spc_zrp, spc_zrp_exact = [], [], [] 
    for m in metrics:
        spc_zr.append(pd.concat([res_jsp[m][tp]["zr"] for tp in tps]).assign(idx_metric=m).set_index("idx_metric", append=True))
        spc_zrp.append(pd.concat([res_jsp[m][tp]["zrp"] for tp in tps]).assign(idx_metric=m).set_index("idx_metric", append=True))
        spc_zrp_exact.append(pd.concat([res_jsp[m][tp]["zrp_exact"] for tp in tps]).assign(idx_metric=m).set_index("idx_metric", append=True))
    spc_zr, spc_zrp, spc_zrp_exact = pd.concat(spc_zr), pd.concat(spc_zrp), pd.concat(spc_zrp_exact)
    spc_zrp.index = pd.MultiIndex.from_tuples([(m, tp) for m in metrics for tp in tps])
    spc_zrp_exact.index = pd.MultiIndex.from_tuples([(m, tp) for m in metrics for tp in tps])

    # save
    spc_zr.to_csv(join(wd, "results", "spc_zr.csv"))
    spc_zrp.to_csv(join(wd, "results", "spc_zr_sig.csv"))
    spc_zrp_exact.to_csv(join(wd, "results", "spc_zr_sig_exact.csv"))
    
else:
    
    # load
    spc_zr = pd.read_csv(join(wd, "results", "spc_zr.csv"), index_col=[0,1,2,3])
    spc_zrp = pd.read_csv(join(wd, "results", "spc_zr_sig.csv"), index_col=[0,1])
    spc_zrp_exact = pd.read_csv(join(wd, "results", "spc_zr_sig_exact.csv"), index_col=[0,1])
    
# merge and print overview results
data_spc = data.copy()
for m in metrics:
    data_spc = data_spc.merge(
        spc_zr.loc[na(),na(),na(), m].rename(columns=dict(zip(spc_zr.columns, [m+"_"+c for c in spc_zr.columns]))), 
        left_index=True, 
        right_index=True
    )

print("Average Spearman colocalization")
display(spc_zr.groupby(["idx_metric", "idx_tp"], sort=False).mean().loc[(na(), "T0"),:].round(2))
print("Associated group-permutation p-values (norm)")
display(spc_zrp.loc[(na(), "T0"),:].round(5))
print("Associated group-permutation p-values (exact)")
display(spc_zrp_exact.loc[(na(), "T0"),:].round(5))

# %% [markdown]
### SPC statistics

# %% [markdown]
#### Cross: PP-NP each timepoint

# %% [code]
spc_cross = list()

for tp in tps:
    temp = dict()
    
    for m in metrics:
        for receptor in spc_zr.columns:
                
            temp[(m, receptor)] = pd.DataFrame({
                (tp, "rho (mean)"): spc_zr.groupby(["idx_metric", "idx_tp"]).mean().loc[(m, tp),receptor].round(2),
                (tp, "rho (sd)"): spc_zr.groupby(["idx_metric", "idx_tp"]).std().loc[(m, tp),receptor].round(2),
                (tp, "p(exact)"): spc_zrp_exact.loc[(m, tp), receptor],
                (tp, "p(norm)"): spc_zrp.loc[(m, tp), receptor]
            }, index=range(1))    

    spc_cross.append(pd.concat(temp))    

spc_cross = pd.concat(spc_cross, axis=1)
spc_cross.index = spc_cross.index.droplevel(2)
spc_cross.index.names = ["Measure", "Receptor"]

spc_cross[("T0", "q(norm)")] = mc_correction(spc_cross[("T0", "p(norm)")])[0].round(4)
spc_cross.loc[spc_cross["T0", "q(norm)"] > 0.05, (tps[1:],na())] = np.nan

spc_cross.loc[spc_cross["T0", "q(norm)"] <= 0.05, (tps[1:],"p(norm)")]
temp = mc_correction(spc_cross.loc[spc_cross["T0", "q(norm)"] <= 0.05, (tps[1:],"p(norm)")])[0].round(3)
for tp in tps[1:]:
    spc_cross.loc[spc_cross["T0", "q(norm)"] <= 0.05, (tp, "q(norm)")] = temp[(tp, "p(norm)")]

spc_cross = spc_cross[[(tp, c) for tp in tps for c in ["rho (mean)", "rho (sd)", "p(exact)", "p(norm)", "q(norm)"]]]

spc_cross.to_csv(join(wd, "tables", "spc_cross_stat.csv"))
spc_cross.to_html(join(wd, "tables", "spc_cross_stat.html"))
spc_cross 

# %% [markdown]
#### Long: LMM

# %% [code]
spc_long = dict()

for m in metrics:
    for receptor in spc_zr.columns:
        if spc_cross.loc[(m, receptor), ("T0", "q(norm)")] < 0.05:
            print(m, receptor)
        
            temp = data_spc.loc[(subs_PP_long, na(), "PP"), [f"{m}_{receptor}", "weeks_postpartal", "ID"]].dropna()
            lmm = smf.mixedlm(f"{m}_{receptor} ~ weeks_postpartal", data=temp, groups="ID").fit()
            lmm2 = smf.mixedlm(f"{m}_{receptor} ~ weeks_postpartal + np.square(weeks_postpartal)", data=temp, groups="ID").fit()
        
            spc_long[(m, receptor)] = pd.DataFrame({
                ("", "df"): f"({lmm.df_modelwc}, {lmm.df_resid})",
                ("Weeks PP", "converged"): lmm.converged,
                ("Weeks PP", "coef"): lmm.params["weeks_postpartal"].round(4),
                ("Weeks PP", "p"): lmm.pvalues["weeks_postpartal"],
                ("Weeks PP^2", "converged"): lmm2.converged,
                ("Weeks PP^2", "coef"): lmm2.params["np.square(weeks_postpartal)"].round(4),
                ("Weeks PP^2", "p"): lmm2.pvalues["np.square(weeks_postpartal)"],
            }, index=range(1))
        
spc_long = pd.concat(spc_long)
spc_long.index = spc_long.index.droplevel(2)
spc_long.index.names = ["Measure", "Receptor"]


temp = mc_correction(spc_long.loc[:, (["Weeks PP", "Weeks PP^2"], "p")])[0].round(3)
spc_long[("Weeks PP", "q")] = temp[("Weeks PP", "p")]
spc_long[("Weeks PP^2", "q")] = temp[("Weeks PP^2", "p")]

spc_long[("Weeks PP", "p")] = spc_long[("Weeks PP", "p")].round(3)
spc_long[("Weeks PP^2", "p")] = spc_long[("Weeks PP^2", "p")].round(3)

spc_long = spc_long[[("", "df")] + [(t, c) for t in ["Weeks PP", "Weeks PP^2"] for c in ["converged", "coef", "p", "q"]]]

spc_long.to_csv(join(wd, "tables", "spc_long_stat.csv"))
spc_long.to_html(join(wd, "tables", "spc_long_stat.html"))
spc_long 

# %% [markdown]
#### Cross: Explained variance (for in-text reporting)

# %% [code]
rerun_spc_r2 = False

# %% [code]
temp = (spc_cross.loc[:,("T0", "q(norm)")] < 0.05)
receptors = temp[temp == True].index.get_level_values("Receptor").unique().tolist()
print("Receptors with >= 1 FDR-significant Spearman colocalization:")
print(receptors)

if rerun_spc_r2:
    
    metrics = ["fALFF", "LCOR", "GCOR"]
    
    spc_cross_r2 = pd.DataFrame(index=metrics, columns=["mean", "std", "min", "max", "p(exact)"])
    for m in metrics:
        jsp = JuSpyce(
            x=pred_aba_pet[receptors].T,
            y=pd.read_csv(join(wd, "results", f"spc_pp_data_{m}.csv"), index_col=[0,1,2]),
            data_space="mni152",
            parcellation=parc,
            parcellation_space="mni152",
            parcellation_labels=parc_info.label.to_list(),
            standardize="x",
            drop_nan=False,
            n_proc=8
        ).fit(verbose=False)
        groups = (data.group.replace({0:2}) * ((data.tp == "T0") | (data.group_str == "NP"))).replace({0:np.nan})
        jsp.compare(comparison="z(A,B)", groups=groups, verbose=False)
        jsp.predict(comparison="z(A,B)", method="mlr", adjust_r2=True, mlr_individual=False, verbose=False)
        jsp.permute_groups(
            comparison="z(A,B)",
            groups=groups,
            method="mlr",
            p_from_average_y="mean",
            adjust_r2=True, 
            mlr_individual=False,
            n_perm=10000,
            seed=42,
            verbose=True
        )
        spc_cross_r2.loc[m, ["mean", "std", "min", "max"]] = \
            jsp.predictions["z(A,B)-mlr_full_r2"]["mlr_full_r2"].apply(["mean", "std", "min", "max"])
        spc_cross_r2.loc[m, "p(exact)"] = \
            jsp.p_comparisons["z(A,B)-mlr_full_r2"].iloc[0, 0]
        spc_cross_r2.to_csv(join(wd, "results", "spc_r2_res.csv"))
        
else:
    spc_cross_r2 = pd.read_csv(join(wd, "results", "spc_r2_res.csv"), index_col=0)

spc_cross_r2


# %% [markdown]
## Associations between MRI and hormones/behavior

# %% [code]
assoc_dfs = dict()

# iterate hormones/behav
for analysis, y_vars in zip(
    ["horm", "behav"],
    [["progesterone_log", "estradiol_log", "ep_ratio_log"], ["epds",  "mpas", "mpas_attachment", "mpas_hostility", "mpas_interaction"]]
):
    x_vars = cl_labs + [i[0] + "_" + i[1] for i in spc_cross[spc_cross[("T0", "q(norm)")] < 0.05].index]
    x_vars = [x for x in x_vars if "falff" in x.lower()] + [x for x in x_vars if "lcor" in x.lower()] + [x for x in x_vars if "gcor" in x.lower()]
    
    # hormones: "Hormones predict brain function"
    if analysis == "horm":
        x_vars, y_vars = y_vars, x_vars
    # behavior: "Brain function predicts behavior"
    else:
        pass
    print(analysis, "X:", x_vars)
    print(analysis, "Y:", y_vars)
    
    assoc = dict()

    for y in y_vars:
        for x in x_vars:
            
            temp = data_spc.loc[(subs_PP_long, na(), "PP"), [y, x, "ID", "weeks_postpartal", "age"]].dropna()
            for var in [x]:
                for sub in subs_PP_long:
                    temp_sub = temp.loc[(sub, na(), "PP"), var]
                    if len(temp_sub.unique()) == 1:
                        temp_sub = temp_sub - temp_sub.mean()
                    else:
                        temp_sub = (temp_sub - temp_sub.mean()) / temp_sub.std()
                    temp.loc[(sub, na(), "PP"), var] = temp_sub
            temp[x] = (temp[x] - temp[x].mean()) / temp[x].std(ddof=1)
            temp[y] = (temp[y] - temp[y].mean()) / temp[y].std(ddof=1)
                
            formula = f"{y} ~ {x} * weeks_postpartal"
            print(formula)
            lmm = smf.mixedlm(formula, groups="ID", re_formula="1", data=temp).fit()
            
            assoc[(x, y)] = pd.DataFrame({
                ("", "df"): f"({lmm.df_modelwc}, {lmm.df_resid})",
                ("", "converged"): lmm.converged,
                ("X", "coef"): lmm.params[x].round(4),
                ("X", "p"): lmm.pvalues[x].round(3),
                ("Weeks PP", "coef"): lmm.params["weeks_postpartal"].round(4),
                ("Weeks PP", "p"): lmm.pvalues["weeks_postpartal"].round(3),
                ("Interaction", "coef"): lmm.params[x+":weeks_postpartal"].round(4),
                ("Interaction", "p"): lmm.pvalues[x+":weeks_postpartal"],
            }, index=range(1))
            
    assoc = pd.concat(assoc)
    assoc.index = assoc.index.droplevel(2)
    assoc.index.names = ["X", "Y"]
    assoc = assoc \
        .reset_index() \
        .replace(
            {"progesterone_log": "log(Progesterone)",
             "estradiol_log": "log(Estradiol)",
             "ep_ratio_log": "log(Estradiol/Progesterone)",
             "mpas":"MPAS - total",
             "mpas_attachment": "MPAS - attachment", 
             "mpas_hostility": "MPAS - hostility", 
             "mpas_interaction": "MPAS - interaction",
             "epds": "EPDS"} | 
            dict(zip(cl_labs, cl_names)) |
            dict(zip([i[0] + "_" + i[1] for i in spc_cross.index],
                     [f"rho({i[0]}, {i[1]})" for i in spc_cross.index]))
        ) \
        .set_index([("X", ""), ("Y", "")])
    assoc.index.names = ["X", "Y"]
    assoc[("Interaction", "q")] = mc_correction(assoc[("Interaction", "p")])[0].round(3)
    assoc[("Interaction", "p")] = assoc[("Interaction", "p")].round(3)
    
    assoc = assoc.sort_index(level="X" if analysis=="horm" else "Y")
        
    assoc.to_csv(join(wd, "tables", f"assoc_{analysis}_stat.csv"))
    assoc.to_html(join(wd, "tables", f"assoc_{analysis}_stat.html"))
    
    assoc_dfs[analysis] = assoc.copy()
    
    display(assoc[assoc[("Interaction", "p")] < 0.05])

assoc_horm, assoc_behav = assoc_dfs["horm"], assoc_dfs["behav"]

# %% [markdown]
## Regression to the mean sensitivity analyses

# %% [code]
subs_rtm = subs_PP_onlybl + subs_NP

# %% [markdown]
### GLM

# %% [markdown]
#### Prepare model

# %% [code]
rerun_sec_lev_rtm = False

sec_lev_desi_rtm = pd.DataFrame({
    "pp": (data.loc[(subs_rtm,"T0",na()),:].group_str == "PP").astype(int),
    "np": (data.loc[(subs_rtm,"T0",na()),:].group_str == "NP").astype(int),
    "age": data.loc[(subs_rtm,"T0",na()),:].age
})

print(f"N = {data.loc[(subs_rtm,'T0',na()),:].shape[0]} subjects")
plot_design_matrix(sec_lev_desi_rtm, rescale=False)

# %% [markdown]
#### Run only GCOR

# %% [code]
# iterate voxe-wise measures
sec_level_res_rtm = {}
sec_level_res_rtm["gcor"] = {}

# RERUN
if rerun_sec_lev_rtm:
    
    # get data files
    vols = [load_img(d) for d in data.loc[(subs_rtm,"T0",na()), "gcor_path"].to_list()]
    
    # con1
    sec_level_res_rtm["gcor"]["con1"] = run_2nd_level(
        data=vols,
        design=sec_lev_desi_rtm,
        contrast_par=np.array([1, -1, 0]),
        contrast_nonpar=np.array([1, 0, 0]),
        save_dir=join(wd, "results"),
        save_prefix="RTM_gcor_con1_",
        cmap="cold_white_hot", 
        title=f"{measure}: PP > NP [-lg(p)]"
    )
    
    # con2
    sec_level_res_rtm["gcor"]["con2"] = run_2nd_level(
        data=vols,
        design=sec_lev_desi_rtm,
        contrast_par=np.array([-1, 1, 0]),
        contrast_nonpar=np.array([0, 1, 0]),
        save_dir=join(wd, "results"),
        save_prefix="RTM_gcor_con2_",
        cmap="cold_white_hot_r", 
        title=f"{measure}: PP < NP [-lg(p)]"
    )

else:
    # NOT RERUN, JUST LOAD SAVED FILES
    for con in ["con1", "con2"]:
        sec_level_res_rtm["gcor"][con] = {}
        for file in glob(join(wd, "results", f"RTM_gcor_{con}_*.nii.gz")):
            name = os.path.basename(file).replace(f"RTM_gcor_{con}_", "").replace(".nii.gz", "")
            sec_level_res_rtm["gcor"][con][name] = load_img(file)

# show results dict
sec_level_res_rtm

# %% [markdown]
#### Extract cluster data

# %% [code]
rerun_cl_extract_rtm = False

# %% [code]
if rerun_cl_extract_rtm:

    # VOXEL-WISE
        
    # image data
    vols_func = [load_img(d) for d in data["gcor_path"].to_list()]
        
    for con in ["con1", "con2"]:
        
        # indexed thresholded image
        if "map_nonpar_idx" in sec_level_res_rtm["gcor"][con].keys():
            
            # masker
            masker = NiftiLabelsMasker(
                labels_img=sec_level_res["gcor"][con]["map_nonpar_idx"],
                standardize=False,
            )
            # extract functional and structural data
            temp_func = masker.fit_transform(vols_func)
            for cl in range(temp_func.shape[1]):
                data[f"rtm_gcor_{con}_cl{cl+1}"] = temp_func[:, cl]

    # SAVE    
    data.to_csv(join(wd, "pheno", "pprs_pheno_mri_proc_res_rtm.csv"))
    
else:
    data = pd.read_csv(join(wd, "pheno", "pprs_pheno_mri_proc_res_rtm.csv"))
    data = data.set_index(["idx_id", "idx_tp", "idx_group"])

# %% [markdown]
### Spatial correlations

# %% [code]
rerun_spc_rtm = False

# %% [code]
if rerun_spc_rtm:
    
    # run
    res_jsp_rtm = dict()
    res_jsp_rtm["gcor"] = dict()
    res_jsp_rtm["gcor"]["T0"] = dict()

    jsp = JuSpyce(
        x=pd.concat([pred_aba, pred_pet], axis=1).T,
        y=data.loc[(subs_rtm, "T0", na()), "gcor_path"].to_list(),
        z=[tpm_gm],
        data_space="mni152",
        parcellation=parc,
        parcellation_space="mni152",
        parcellation_labels=parc_info.label.to_list(),
        y_labels=data.loc[(subs_rtm, "T0", na()),:].index,
        standardize="x",
        drop_nan=False,
        n_proc=8
    ).fit(verbose=False)
    
    groups = data.loc[(subs_rtm, "T0", na()), "group"].replace({0:2})
    jsp.transform("partial", dataset="X", replace=True, verbose=False)
    jsp.transform("partial", dataset="Y", replace=True, verbose=False)
    jsp.compare(comparison="z(A,B)", groups=groups, verbose=False)
    jsp.predict(comparison="z(A,B)", method="spearman", r_to_z=True, verbose=False)
    jsp.permute_groups(
        comparison="z(A,B)",
        groups=groups,
        method="spearman",
        p_from_average_y="mean",
        r_to_z=True,
        n_perm=10000,
        seed=42,
        verbose=True
    )

    res_jsp_rtm["gcor"]["T0"]["zr"] = jsp.predictions["z(A,B)-spearman"]
    res_jsp_rtm["gcor"]["T0"]["zrp"] = jsp.p_comparisons["z(A,B)-spearman-norm"]
        
    # collect results
    spc_zr_rtm = res_jsp_rtm["gcor"]["T0"]["zr"].assign(idx_metric="gcor").set_index("idx_metric", append=True)
    spc_sig_rtm = res_jsp_rtm["gcor"]["T0"]["zrp"].assign(idx_metric="gcor").set_index("idx_metric", append=True)
    #spc_sig.index = pd.MultiIndex.from_tuples([(m, tp) for m in metrics for tp in tps])
    
    # save
    spc_zr_rtm.to_csv(join(wd, "results", "rtm_spc_zr.csv"))
    spc_sig_rtm.to_csv(join(wd, "results", "rtm_spc_zr_sig.csv"))
    
else:
    
    # load
    spc_zr_rtm = pd.read_csv(join(wd, "results", "rtm_spc_zr.csv"), index_col=[0,1,2,3])
    spc_sig_rtm = pd.read_csv(join(wd, "results", "rtm_spc_zr_sig.csv"), index_col=[0,1])    
    
# merge and print overview results
data_spc_rtm = data
data_spc_rtm = data_spc_rtm.merge(
    spc_zr_rtm.loc[na(),na(),na(), "gcor"].rename(columns=dict(zip(spc_zr_rtm.columns, ["rtm_gcor_"+c for c in spc_zr_rtm.columns]))), 
    left_index=True, 
    right_index=True
)

print("Average Spearman colocalization")
display(spc_zr_rtm.groupby(["idx_metric", "idx_tp"], sort=False).mean().loc[(na(), "T0"),:].round(2))
print("Associated group-permutation p-values")
display(spc_sig_rtm)

# %% [markdown]
### Repeat longitudinal analyses

# %% [code]
cl_iter_rtm = [
    ('rtm_gcor_con2_cl1', 'GCOR', 'PP < NP', 'leftPut'),
    ('rtm_gcor_con2_cl2', 'GCOR', 'PP < NP', 'rightPut')
]

mri_long_lmm_rtm = list()

for lab, measure, contrast, name in cl_iter_rtm:
    print(lab)
    
    # longitudinal data, only PP, doesnt matter if all timepoints
    temp_mri_long = data.loc[(subs_PP_long, na(), "PP"), :]
    
    # calculate LMM and pairwise paired t tests
    lmm = smf.mixedlm(f"{lab} ~ weeks_postpartal", data=temp_mri_long, groups="ID").fit()
    lmm2 = smf.mixedlm(f"{lab} ~ weeks_postpartal + np.square(weeks_postpartal)", data=temp_mri_long, groups="ID").fit()

    temp_df = pd.DataFrame({
        "Measure": measure,
        "Contrast": contrast,
        "Cluster name": name,
        ("", "df"): f"({lmm.df_modelwc}, {lmm.df_resid})",
        ("Weeks PP", "converged"): lmm.converged,
        ("Weeks PP", "coef"): lmm.params["weeks_postpartal"].round(4),
        ("Weeks PP", "-lg(p)"): -np.log10(lmm.pvalues["weeks_postpartal"]).round(3),
        ("Weeks PP^2", "converged"): lmm2.converged,
        ("Weeks PP^2", "coef"): lmm2.params["np.square(weeks_postpartal)"].round(4),
        ("Weeks PP^2", "-lg(p)"): -np.log10(lmm2.pvalues["np.square(weeks_postpartal)"]).round(3),
    }, index=range(1))
    
    # save
    temp_df = temp_df.set_index(["Measure", "Contrast", "Cluster name"])
    mri_long_lmm_rtm.append(temp_df)
        
mri_long_lmm_rtm = pd.concat(mri_long_lmm_rtm)
mri_long_lmm_rtm.columns = pd.MultiIndex.from_tuples(mri_long_lmm_rtm.columns)

mri_long_lmm_rtm.to_csv(join(wd, "tables", "rtm_mri_longitudinal_lmm_stat.csv"))
mri_long_lmm_rtm.to_html(join(wd, "tables", "rtm_mri_longitudinal_lmm_stat.html"))
mri_long_lmm_rtm

# %% [markdown]
## Plots

# %% [code]
def where(condition, x, y):
    out = np.where(condition, x, y)
    if isinstance(condition, pd.DataFrame):
        out = pd.DataFrame(out, columns=condition.columns, index=condition.index)
    return out

def lgp_to_star(lgp_array, lgq_array, empty=""):
    lgp_array_flat = np.array(lgp_array).flatten()
    lgq_array_flat = np.array(lgq_array).flatten()
    a = list()
    for lgp, lgq in zip(lgp_array_flat, lgq_array_flat):
        if lgq > -np.log10(0.05): a.append("★")
        elif lgp > -np.log10(0.05): a.append("☆")
        else: a.append(empty)
    
    return np.array(a).reshape(lgp_array.shape, order="C")

# colors
colors = dict()
for sub in subs_NP: 
    colors[sub] = "tab:blue"
for sub in [s for s in subs_PP if s not in subs_PP_long]: 
    colors[sub] = "0.5"
temp = np.array([
    [0.12156863, 0.46666667, 0.70588235, 1.        ],
    [1.        , 0.49803922, 0.05490196, 1.        ],
    [0.17254902, 0.62745098, 0.17254902, 1.        ],
    [0.83921569, 0.15294118, 0.15686275, 1.        ],
    [0.58039216, 0.40392157, 0.74117647, 1.        ],
    [0.54901961, 0.3372549 , 0.29411765, 1.        ],
    [0.89019608, 0.46666667, 0.76078431, 1.        ],
    [0.49803922, 0.49803922, 0.49803922, 1.        ],
    [0.7372549 , 0.74117647, 0.13333333, 1.        ],
    [0.09019608, 0.74509804, 0.81176471, 1.        ],
    [0.6       , 0.2       , 0.8       , 1.        ],
    [0.        , 0.6       , 0.5       , 1.        ],
    [0.9       , 0.6       , 0.        , 1.        ],
    [0.35      , 0.7       , 0.9       , 1.        ],
    [0.8       , 0.4       , 0.        , 1.        ],
    [0.5       , 0.        , 0.5       , 1.        ],
    [0.        , 0.5       , 0.        , 1.        ],
    [0.9       , 0.1       , 0.1       , 1.        ],
    [0.6       , 0.4       , 0.2       , 1.        ]
])
for i, sub in enumerate(subs_PP_long): 
    colors[sub] = temp[i] #get_cmap("tab20")(i)

# %% [markdown]
### MRI brains - cross-sectional

# %% [code]
vmin = 0
vmax = 6

fig, axes = plt.subplots(4,2, figsize=(20,9),
                         gridspec_kw=dict(
                             hspace=0.05, 
                             wspace=0.05, 
                             height_ratios=(1,1,1, 0.08)))

# iterate contrasts
for c, con, con_name in zip(
    range(2), 
    ["con1", "con2"],
    ["PP > NP", "PP < NP"]):
    
    # VOXELWISE BRAINS
    for r, measure, measure_name in zip(
        range(3), 
        ["falff", "lcor", "gcor"], 
        ["fALFF", "LCOR", "GCOR"]):
        
        # get maps
        map_unthresh = sec_level_res[measure][con]["map_par_logp"]
        map_thresh = sec_level_res[measure][con]["map_nonpar_logp"]
        
        # plot main glass brain
        gb = plot_glass_brain(
            map_unthresh, 
            display_mode="lzry",
            alpha=0.4,
            colorbar=False, 
            cmap="cold_white_hot" if con=="con1" else "cold_white_hot_r",
            figure=fig,
            axes=axes[r,c],
            vmin=vmin,
            vmax=vmax
        )
        
        # significant cluster
        if (map_thresh.get_fdata() > cl_thresh).any():
            
            # plot overlays
            gb.add_contours(
                img=threshold_img(map_thresh, cl_thresh), 
                levels=[1], 
                colors="k", 
                linewidth=5,
                filled=True,
                alpha=0.5
            )
        
        # titles
        if r==0:
            axes[r,c].set_title(f"T0: {con_name} $(n = {data.loc[(na(),'T0','PP'),:].shape[0]}\ |\ "
                                f"{data.loc[(na(),'T0','NP'),:].shape[0]})$", 
                                size=16, weight="semibold", pad=10)
        if c==0: 
            axes[r,c].annotate(
                measure_name, 
                xy=(-0.04,0.5), 
                xycoords="axes fraction", 
                rotation=90,
                size=16, weight="semibold", ha="center", va="center"
            )
    
    # COLORBAR
    norm = mpl.colors.Normalize(vmin=vmin, vmax=vmax)
    sm = plt.cm.ScalarMappable(
        cmap="black_red_r" if con=="con1" else "black_blue_r", 
        norm=norm)
    sm.set_array([])
    plt.colorbar(
        sm,
        label=r"$-lg(p)$",
        orientation="horizontal",
        cax=axes[3,c],
    )
    
# save
fig.savefig(join(wd, "plots", "fig1_brains.pdf"), bbox_inches="tight", dpi=400)
fig.savefig(join(wd, "plots", "fig1_brains.png"), bbox_inches="tight", dpi=400)

# %% [markdown]
### Cross-sectional associations / sensitivity analyses

# %% [code]
vars = ["birth_mode_str", "baby_sex_str", "n_children", 
        "age", "weeks_postpartal", "baby_weight", "gestation_day", "epds", "fwd_mean", "fwd_max"]
var_names = ["Birth mode", "Sex of baby", "Number of children", 
             "Age", "Postpartum week", "Weight of baby", "Gestation day", "EPDS", "Mean FWD", "Max FWD"]

if "group_crosslong_str" not in data.columns:
    # new variable with cross vs long subjects
    data["group_crosslong_str"] = data.group_str.copy()
    data.loc[(subs_PP_long, na(), "PP"), "group_crosslong_str"] = "PP-long"
    data.loc[(subs_PP_onlybl, na(), "PP"), "group_crosslong_str"] = "PP-cross"
    
fig, axes = plt.subplots(len(cl_labs), len(vars), figsize=(27, 17), sharex="col", sharey="row")

for r, (lab, m, contrast, name) in enumerate(cl_iter):
    for c, (var, var_name) in enumerate(zip(vars, var_names)):
        temp = data.loc[(na(), "T0", "PP"), [var, lab, "group_crosslong_str"]].dropna()
        if var=="n_children":
            temp[var] = temp[var].astype(int)
        # if var=="birth_mode_str":
        #     temp = temp.query("birth_mode_str in ['spontaneous','section']")
        ax = axes[r,c]
        kwargs = dict(
            x=temp[var],
            y=temp[lab],
            ax=ax
        )
        kwargs_scat = dict(
            alpha=0.5,
            hue=temp["group_crosslong_str"],
            palette="tab10",
            linewidth=0.5,
            edgecolor="k",
        )
        
        # categorical
        if var in ["birth_mode_str", "baby_sex_str", "n_children"]:
            sns.stripplot(
                **kwargs,
                **kwargs_scat,
                zorder=1,
                size=5
            )
            sns.boxplot(
                **kwargs,
                width=0.5,
                fliersize=0,
                boxprops=dict(facecolor="none"),
                zorder=2,
                linewidth=1,
            )
            f, df, p, np2 = mri_bl_assoc.iloc[r,:][(f"ANCOVA: {var_name}", na())]
            s = f"$F{df} = {f}$\n $p = {p}$\n$np2 = {np2}$"
            
        # continuous
        else:
            sns.scatterplot(
                **kwargs,
                **kwargs_scat,
                sizes=8
            )
            sns.regplot(
                **kwargs,
                scatter=False,
            )   
            coef, df, p = mri_bl_assoc.iloc[r,:][(f"MLR: {var_name}", na())]
            s = f"$df = {df}$\n$Beta = {coef}$\n$p = {p}$" 

        ax.annotate(s, xy=(0.96, 0.05), xycoords="axes fraction", ha="right", va="bottom",
                    bbox=dict(boxstyle="round", fc="tab:red" if p < 0.05 else "w", alpha=0.5))
        
        if ax.get_subplotspec().is_first_row():
            ax.set_title(var_name, weight="semibold", size=14, pad=10)
        
        if ax.get_subplotspec().is_last_row():
            ax.set_xlabel(var_name, size=12)
            if "birth_mode" in var:
                ax.set_xticks(ax.get_xticks())
                ax.set_xticklabels(ax.get_xticklabels(), rotation_mode="anchor", ha="left", rotation=-75)
        else:
            ax.set_xlabel("")
        
        if ax.get_subplotspec().is_first_col():
            ax.set_ylabel(m + " $[Z]$", size=12)
            ax.annotate(f"{m}\n{contrast}\n{name}", xy=(-0.5,0.5), xycoords="axes fraction", 
                        size=15, weight="semibold", ha="right", va="center")
        else:
            ax.set_ylabel("")
        
        if ax.get_subplotspec().is_last_col():
            handles, labels = ax.get_legend_handles_labels()
            ax.legend(handles, ["longitudinal" if "long" in l else "cross-sectional" for l in labels], 
                      loc="center left", bbox_to_anchor=(1, 0.5), title="Postpartum sample")
        else:
            ax.legend().remove()

fig.tight_layout() 
fig.savefig(join(wd, "plots", "figS1_pp_sensitivity.pdf"), bbox_inches="tight")
fig.savefig(join(wd, "plots", "figS1_pp_sensitivity.png"), bbox_inches="tight", dpi=400)

# %% [markdown]
### MRI brains - longitudinal 

# %% [code]
es_lims = {}
for c in sec_level_res_long["falff"].keys():
    es_lims[c] = [np.max(np.abs(sec_level_res_long[m][c].get_fdata())) for m in ["falff", "lcor", "gcor"]]
    es_lims[c] = (-max(es_lims[c]), max(es_lims[c]))

contrasts = [
    'weeks_postpartal', 
    'estradiol_log*weeks_postpartal', 'progesterone_log*weeks_postpartal', 'ep_ratio_log*weeks_postpartal', 
    'epds*weeks_postpartal',
    'mpas*weeks_postpartal',  'mpas_attachment*weeks_postpartal', 'mpas_interaction*weeks_postpartal', 'mpas_hostility*weeks_postpartal'
]
contrast_names = [
    'Weeks PP', 
    'log(Estradiol) × Weeks PP', 'log(Progesterone) × Weeks PP', 'log(Estradiol/Progesterone) × Weeks PP', 
    'EPDS × Weeks PP',
    'MPAS - Total × Weeks PP',  'MPAS - Attachment × Weeks PP', 'MPAS - Interaction × Weeks PP', 'MPAS - Hostility × Weeks PP'
]

fig, axes = plt.subplots(len(contrasts), 3, figsize=(17,16), 
                         gridspec_kw=dict(wspace=0.1, hspace=0.05))
for c, measure in enumerate(["fALFF", "LCOR", "GCOR"]): 
    axes[0,c].set_title(measure, weight="semibold", size=16)
    
    for r, (contrast, contrast_name) in enumerate(zip(contrasts, contrast_names)):
        
        gb = plot_glass_brain(
            sec_level_res_long[measure.lower()][contrast], 
            display_mode="lzry",
            alpha=0.4,
            colorbar=False, 
            cmap="RdBu_r",
            plot_abs=False,
            figure=fig,
            axes=axes[r,c],
            vmin=es_lims[contrast][0],
            vmax=es_lims[contrast][1]
        )  
        
        # ANNOTATE
        if c==0:
        
            # COLORBAR
            ax_cbar = axes[r,c].inset_axes((-0.8, 0.5, 0.7, 0.1))
            sm = plt.cm.ScalarMappable(
                cmap="RdBu_r",
                norm=mpl.colors.Normalize(*es_lims[contrast]))
            sm.set_array([])
            fig.colorbar(
                sm,
                label="Standardized Beta",
                orientation="horizontal",
                cax=ax_cbar
            )
            ax_cbar.set_title(contrast_name, size=12, weight="semibold")
        
# save
fig.savefig(join(wd, "plots", "figS3_brains_long.png"), bbox_inches="tight", dpi=300)

# %% [markdown]
### MRI brains - long/cross correlation

# %% [code]
temp = pd.read_csv(join(wd, "results", "spc_longcross_data.csv"), index_col=0).T
temp_res = pd.read_csv(join(wd, "results", "spc_longcross.csv"), index_col=0, header=[0,1])
fig, axes = plt.subplots(len(temp.columns)-1, len(temp.columns)-1, figsize=(16, 15), 
                         sharey="row", sharex="col")

for r, measure_r in enumerate(temp.columns[1:]):
    for c, measure_c in enumerate(temp.columns[:-1]):
        ax = axes[r,c]
        if r < c:
            ax.set_axis_off()
        else:        
            sns.regplot(
                y=temp[measure_r],
                x=temp[measure_c],
                ax=ax,
                scatter_kws=dict(alpha=0.5),
                color="tab:red" if measure_r.split("_")[0]==measure_c.split("_")[0] 
                    else "tab:green" if measure_r.split("_")[1]==measure_c.split("_")[1] 
                    else "tab:blue"
            )
            rho = temp_res.loc[measure_r, (measure_c, "rho")]
            p = (temp_res.loc[measure_r, (measure_c, "p")] + temp_res.loc[measure_c, (measure_r, "p")]) / 2
            ax.annotate(
                f"$Z(rho) = {rho:.02f}$"
                f"\n$p(exact) = {p:.04f}$",
                xy=(0.05,0.95),
                xycoords="axes fraction",
                ha="left", va="top",
                size=12
            )
            if ax.get_subplotspec().is_last_row():
                ax.set_xlabel(
                    measure_c \
                        .replace("falff", "fALFF").replace("gcor", "GCOR").replace("lcor", "LCOR") \
                        .replace("_long", " longitudinal\n$[Z(beta)]$").replace("_cross", " cross-sectional\n$[Z(beta)]$"),
                weight="semibold", 
                size=12
            )
            else:
                ax.set_xlabel("")
            if ax.get_subplotspec().is_first_col():
                ax.set_ylabel(
                    measure_r \
                        .replace("falff", "fALFF").replace("gcor", "GCOR").replace("lcor", "LCOR") \
                        .replace("_long", " longitudinal\n$[Z(beta)]$").replace("_cross", " cross-sectional\n$[Z(beta)]$"), 
                    weight="semibold", 
                    size=12
                )
            else:
                ax.set_ylabel("")

fig.tight_layout()
fig.savefig(join(wd, "plots", "figS4_brain_spc.pdf"), bbox_inches="tight")
fig.savefig(join(wd, "plots", "figS4_brain_spc.png"), bbox_inches="tight", dpi=300)

# %% [markdown]
### MRI group comparisons & development

# %% [code]
# maximum effect size
es_lim = np.nanmax(np.abs(np.r_[
    mri_bl_confounds[("ANCOVA: PP vs. NP: Age", "g")].values,
    mri_long_unpaired[(f"ANCOVA: PP-{tp} vs. NP: Age", "g")].values,
    mri_long_paired.loc[:, (na(), "g")].values.flatten()
]))

fig, axes = plt.subplots(3,3, figsize=(17,18), sharex=False, constrained_layout=True, gridspec_kw=dict(hspace=0.1))
axes = axes.ravel()

for (lab, measure, contrast, name), i in zip(cl_iter, [0,1,2, 3,4, 6,7]):
    
    temp_mri = pd.concat(
        [data.loc[(subs_NP, ), :].melt(["ID", "group_str", "weeks_postpartal"], lab), # all NP
         data.loc[(subs_PP, "T0", ), :].melt(["ID", "group_str", "weeks_postpartal"], lab).replace({"PP": "PP-T0"})] + \
        [data.loc[(subs_PP_long, tp), :].melt(["ID", "group_str", "weeks_postpartal"], lab).replace({"PP": f"PP-{tp}"}) for tp in ["T1", "T2", "T3", "T4", "T5"]]
    )
    temp_mri.loc[temp_mri.group_str=="NP", "weeks_postpartal"] = -3.5

    sns.scatterplot(
        temp_mri,
        x="weeks_postpartal",
        hue="ID",
        y="value",
        ax=axes[i],
        legend=None,
        alpha=0.4,
        zorder=3,
        palette=colors
    )
    sns.lineplot(
        temp_mri,
        x="weeks_postpartal",
        y="value",
        hue="ID",
        ax=axes[i],
        legend=None,
        alpha=0.4,
        zorder=2,
        palette=colors
    )
    
    for group in temp_mri.group_str.unique():   

        axes[i].boxplot(
            x=temp_mri.loc[temp_mri.group_str==group, "value"],
            positions=[temp_mri.loc[temp_mri.group_str==group, "weeks_postpartal"].mean()],
            widths=2,
            zorder=10,
            showfliers=False,
            medianprops=dict(c="k")       
        )
        axes[i].scatter(
            y=[temp_mri.loc[temp_mri.group_str==group, "value"].mean()],
            x=[temp_mri.loc[temp_mri.group_str==group, "weeks_postpartal"].mean()],
            zorder=11,
            c="k",
            s=30
        )
    
    # LMM
    for var in ["Weeks PP", "Weeks PP^2"]:
        p = mri_long_lmm.loc[(measure, contrast, name), (var, "-lg(p)")]
        q =  mri_long_lmm.loc[(measure, contrast, name), (var, "-lg(q)")]
        if q > -np.log10(0.05): s = "★"
        elif p > -np.log10(0.05): s = "☆"
        else: s = "ns"
        axes[i].annotate(
            f"{'Linear' if var=='Weeks PP' else 'Quadratic'} effect: {s}",
            xy=(0.6, 0.05 + (0.07 if var=='Weeks PP' else 0)),
            xycoords="axes fraction",
            size=11
        )
    
    # Labels
    axes[i].set_title(f"{measure} - {contrast}: {name} cluster",  weight="semibold", size=13, pad=10)
    axes[i].set_ylabel(f"Average {measure} $[Z]$", weight="semibold")
    axes[i].set_xticks([-3.5] + list(range(0,28,3)))
    axes[i].set_xticklabels(["NP", "0\nT0", "3\nT1", "6\nT2", "9\nT3", "12\nT4", "15", "18", "21", "24\nT5", "27"])
    axes[i].set_xlabel("Group | Weeks postpartum | Time point", weight="semibold", labelpad=7)
    
    
    # significance
    comp_p = pd.DataFrame(
        np.zeros((7,7)),
        columns=["NP"]+[f"PP-{tp}" for tp in tps],
        index=["NP"]+[f"PP-{tp}" for tp in tps]
    )
    comp_q, comp_es = comp_p.copy(), comp_p.copy()

    # NP vs PP T0
    comp_p.loc["NP", "PP-T0"] = mri_bl_confounds.loc[(measure, contrast, name), ("ANCOVA: PP vs. NP: Age", "-lg(p)")]
    comp_q.loc["NP", "PP-T0"] = mri_bl_confounds.loc[(measure, contrast, name), ("ANCOVA: PP vs. NP: Age", "-lg(p)")]
    comp_es.loc["NP", "PP-T0"] = mri_bl_confounds.loc[(measure, contrast, name), ("ANCOVA: PP vs. NP: Age", "g")]

    # NP vs PP each timepoint
    for tp in tps:
        comp_p.loc["NP", f"PP-{tp}"] = mri_long_unpaired.loc[(measure, contrast, name), (f"ANCOVA: PP-{tp} vs. NP: Age", "-lg(p)")]
        comp_q.loc["NP", f"PP-{tp}"] = mri_long_unpaired.loc[(measure, contrast, name), (f"ANCOVA: PP-{tp} vs. NP: Age", "-lg(q)")]
        comp_es.loc["NP", f"PP-{tp}"] = mri_long_unpaired.loc[(measure, contrast, name), (f"ANCOVA: PP-{tp} vs. NP: Age", "g")]
        
    # PP vs PP each combi
    for k in [c for c in mri_long_paired.columns if ("T-test" in c[0]) & (c[1]=="-lg(p)")]:
        comp_p.loc["PP-"+k[0].split(" ")[1], "PP-"+k[0].split(" ")[3]] = mri_long_paired.loc[(measure, contrast, name), k]
        comp_es.loc["PP-"+k[0].split(" ")[1], "PP-"+k[0].split(" ")[3]] = mri_long_paired.loc[(measure, contrast, name), (k[0], "g")]
        
    for k in [c for c in mri_long_paired.columns if ("T-test" in c[0]) & (c[1]=="-lg(q)")]:
        comp_q.loc["PP-"+k[0].split(" ")[1], "PP-"+k[0].split(" ")[3]] = mri_long_paired.loc[(measure, contrast, name), k]


    comp_es = comp_es + comp_es.T
    comp_es = comp_es.replace({np.nan:0})
    comp_p = comp_p + comp_p.T
    comp_q = comp_q + comp_q.T
    
    comp_es = comp_es.drop("NP", axis="columns").drop("PP-T5", axis="index")
    comp_p = comp_p.drop("NP", axis="columns").drop("PP-T5", axis="index")
    comp_q = comp_q.drop("NP", axis="columns").drop("PP-T5", axis="index")
    
    comp_sig = lgp_to_star(comp_p, comp_q, "")
    mask = np.zeros(comp_es.shape)        
    mask[np.tril_indices_from(mask, k=-1)] = True
    
    # plot heatmap
    sig_ax = axes[i].inset_axes((0, -0.7, 1, 0.4))
    if i==0:
        cbar_ax = axes[5].inset_axes((0.1, 0.94, 0.8, 0.06)) 
        cbar_ax.set_title("Heatmaps: effect sizes $[g]$", weight="semibold", size=12, pad=10)
    ax = sns.heatmap(
        comp_es,
        vmin=-es_lim,
        vmax=es_lim,
        cmap="RdBu_r",
        cbar=True if i==0 else False,
        cbar_ax=cbar_ax,
        cbar_kws=dict(orientation="horizontal"),
        annot=comp_sig,
        linewidths=1,
        linecolor="w",
        fmt="s",
        mask=mask,
        ax=sig_ax
    )
    ax.xaxis.tick_top()
    ax.yaxis.tick_right()
    ax.set_yticks(np.arange(0.5, 5.6))
    ax.set_yticklabels(["NP"]+[f"PP-{tp}" for tp in tps[:5]], rotation=0)

    if i in [0,3,6]:
        axes[i].annotate(
            measure, 
            xy=(-0.27,0.5), 
            xycoords="axes fraction", 
            rotation=90,
            size=18, weight="semibold", ha="center", va="center"
        )

[ax.axis("off") for ax in axes if ax.axis() == (0.0, 1.0, 0.0, 1.0)]

# save
fig.savefig(join(wd, "plots", "fig2_boxes.pdf"), bbox_inches="tight")
fig.savefig(join(wd, "plots", "fig2_boxes.png"), bbox_inches="tight", dpi=300)

# %% [markdown]
### Spatial associations

# %% [code]
fig = plt.figure(figsize=(22,18))
gs = GridSpec(1, 2, figure=fig, width_ratios=(0.5,1), wspace=0.15)
measures = ["fALFF", "LCOR", "GCOR"]
receptors = set(spc_cross[spc_cross.loc[:, ("T0", "q(norm)")] < 0.05].index.get_level_values("Receptor"))
receptors = [r for r in pred_aba.columns if r in receptors] + [r for r in pred_pet.columns if r in receptors]

## BRAINS
gs_brains = GridSpecFromSubplotSpec(1 + 3 + 1 + len(receptors), 1, gs[:,0], wspace=0.2, hspace=0.2, 
                                    height_ratios=[0.3] + [1]*3 + [0.6] + [1]*len(receptors))
kwargs = dict(
    display_mode="lzry",
    annotate=False,
    cbar_tick_format="%.1f",
    colorbar=True, 
    plot_abs=False,
)
# MRI
ax = fig.add_subplot(gs_brains[0])
ax.set_axis_off()
ax.annotate("Baseline resting-state alteration maps \n$[average\ Z(PP\ vs.\ NP]$",
            xy=(0.5,0.5), ha="center", va="center", size=13, weight="semibold")
for r, m in enumerate(measures, start=1):
    temp_z = pd.read_csv(join(wd, "results", f"spc_pp_data_{m}.csv"), index_col=[0,1,2])
    temp_z = (temp_z.loc[(subs_PP, na(), "PP"), :] - temp_z.loc[(subs_NP, na(), "NP"), :].mean()) / temp_z.loc[(subs_NP, na(), "NP"), :].std()
    temp_z = temp_z.groupby("idx_tp").mean().T
    ax = fig.add_subplot(gs_brains[r])
    plot_glass_brain(
        parcel_data_to_volume(temp_z["T0"].values, parc)[0], 
        title=m, 
        axes=ax,
        **kwargs,
        cmap="icefire"
    )
# ABA/PET
ax = fig.add_subplot(gs_brains[4])
ax.set_axis_off()
ax.annotate("Receptor maps (baseline colocalization q < .05)\n$[normalized\ density]$",
            xy=(0.5,0.3), ha="center", va="center", size=13, weight="semibold")
for r, atlas in enumerate(receptors, start=5):
    ax = fig.add_subplot(gs_brains[r])
    plot_glass_brain(
        parcel_data_to_volume(minmax_scale(pred_aba_pet[atlas]), parc)[0], 
        title=atlas, 
        axes=ax,
        **kwargs,
        cmap="viridis" if atlas in pred_aba.columns else "magma"
    )

## SCATTER/BOXPLOTS
temp = spc_cross[spc_cross.loc[:, ("T0", "q(norm)")] < 0.05]
gs_boxes = GridSpecFromSubplotSpec(1 + 1 + 1 + int(np.ceil(len(temp) / 3)), 3, gs[:,1], wspace=0.2, hspace=0.3, 
                                   height_ratios=[0.075, 1, 0.25] + [1]*int(np.ceil(len(temp) / 3)))

# BASELINE
ax = fig.add_subplot(gs_boxes[0,:])
ax.set_axis_off()
ax.annotate("Colocalization between receptor maps and baseline resting-state alterations",
            xy=(0.5,0.5), ha="center", va="center", size=13, weight="semibold")
temp = spc_zr.loc[(na(), "T0", "PP")].reset_index() \
    .melt(id_vars="idx_metric", value_vars=spc_zr.columns, var_name="receptor", value_name="Z")
axes = [fig.add_subplot(gs_boxes[1, c]) for c in range(len(measures))]
for c, (ax, m) in enumerate(zip(axes, measures)):
    ax.sharey(axes[0])
    kwargs = dict(
        data=temp[temp.idx_metric==m],
        x="receptor",
        y="Z",
        dodge=False,
        ax=ax
    )
    sns.stripplot(
        **kwargs,
        hue="Z",
        hue_norm=(-temp["Z"].abs().max(), temp["Z"].abs().max()),
        palette="icefire",
        jitter=0.3,
        legend=None,
        alpha=0.3,
        edgecolor="k",
        linewidth=0.5,
        zorder=1
    )
    sns.boxplot(
        **kwargs,
        fliersize=0,
        width=0.5,
        boxprops=dict(facecolor="none"),
        zorder=2,
        linewidth=1,
    )
    receptors = [l.get_text() for l in ax.get_xticklabels()]
    ax.scatter(
        x=receptors,
        y=temp[temp.idx_metric==m].groupby("receptor").mean().loc[receptors],
        zorder=20,
        c=["orange" if r in pred_pet.columns else "lightgreen" for r in receptors],
        s=30,
        ec="k"
    ) 
    for x, r in enumerate(spc_zr.columns):
        if spc_cross.loc[(m,r), ("T0", "q(norm)")] < 0.05: s = "★"
        elif spc_cross.loc[(m,r), ("T0", "p(norm)")] < 0.05: s = "☆"
        else: s = ""
        ax.text(
            s=s, 
            x=x, 
            y=spc_zr.loc[(na(), "T0", "PP", m), r].max() + 0.08, 
            ha="center", va="center", size=12
        )  
    ax.axhline(c="k", zorder=0, lw=1)
    ax.set_ylim(None, 0.85)
    ax.set_xlabel("Receptor maps", weight="semibold")
    ax.set_title(m, weight="semibold", size=12)
    ax.set_xticks(ax.get_xticks())
    ax.set_xticklabels(ax.get_xticklabels(), rotation=-30, rotation_mode="anchor", ha="left")
    if c==0:
        ax.set_ylabel("Spatial colocalization $[Z(rho)]$", weight="semibold")
    else:
        ax.set_ylabel("")
    
# LONGITUDINAL
ax = fig.add_subplot(gs_boxes[2,:])
ax.set_axis_off()
ax.annotate("Longitudinal development of significant baseline colocalizations",
            xy=(0.5,0.05), ha="center", va="center", size=13, weight="semibold")
temp = spc_cross[spc_cross.loc[:, ("T0", "q(norm)")] < 0.05]
axes = [fig.add_subplot(gs_boxes[3,0]), fig.add_subplot(gs_boxes[3,1]), fig.add_subplot(gs_boxes[3,2]),
        fig.add_subplot(gs_boxes[4,0]), fig.add_subplot(gs_boxes[4,1]), fig.add_subplot(gs_boxes[4,2]),
        fig.add_subplot(gs_boxes[5,0]), fig.add_subplot(gs_boxes[5,1])]
if len(axes) != len(temp):
    raise ValueError("Adjust longitudinal axes list!")

for i, (idx, ax) in enumerate(zip(temp.index, axes)):
    m, r = idx
    
    temp = data_spc.loc[(na(), na(), "PP"), [f"{m}_{r}", "weeks_postpartal", "ID"]]
    sns.scatterplot(
        temp,
        x="weeks_postpartal",
        hue="ID",
        y=f"{m}_{r}",
        ax=ax,
        legend=None,
        alpha=0.4,
        zorder=3,
        palette=colors
    )
    sns.lineplot(
        temp,
        x="weeks_postpartal",
        y=f"{m}_{r}",
        hue="ID",
        ax=ax,
        legend=None,
        alpha=0.4,
        zorder=2,
        palette=colors
    )
    temp_mean = temp.groupby("idx_tp").mean()
    temp_max = temp.groupby("idx_tp").max()
    for tp in tps:   
        ax.boxplot(
            x=temp.loc[(na(), tp, "PP"), f"{m}_{r}"],
            positions=[temp_mean.loc[tp, "weeks_postpartal"]],
            widths=2,
            zorder=10,
            showfliers=False,
            medianprops=dict(c="k")
        )
        ax.scatter(
            y=temp_mean.loc[tp, f"{m}_{r}"],
            x=temp_mean.loc[tp, "weeks_postpartal"],
            zorder=11,
            c="k",
            s=30
        )        
        if spc_cross.loc[(m,r), (tp, "q(norm)")] < 0.05: s = "★"
        elif spc_cross.loc[(m,r), (tp, "p(norm)")] < 0.05: s = "☆"
        else: s = ""
        ax.text(
            s=s, 
            x=temp_mean.loc[tp, "weeks_postpartal"], 
            y=temp_max.loc[tp, f"{m}_{r}"] + 0.05,
            ha="center", va="center", size=11
        )  
        # LMM
        for var in ["Weeks PP", "Weeks PP^2"]:
            if spc_long.loc[(m,r), (var, "q")] < 0.05: s = "★"
            elif spc_long.loc[(m,r), (var, "p")] < 0.05: s = "☆"
            else: s = "ns"
            ax.annotate(
                f"{'Linear' if var=='Weeks PP' else 'Quadratic'}: {s}",
                xy=(0.4 + (0.35 if var=='Weeks PP' else 0), 0.04),
                xycoords="axes fraction",
                ha="left",
                size=10
            )
    ax.set_ylim(None, temp_max[f"{m}_{r}"].max() + 0.1)
    ax.axhline(c="k", zorder=0, lw=1)
    ax.set_title(f"{m} - {r}", weight="semibold", size=12)
    ax.set_xticks(list(range(0,28,3)))
    if ax.get_subplotspec().is_last_row():
        ax.set_xticklabels(["0\nT0", "3\nT1", "6\nT2", "9\nT3", "12\nT4", "15", "18", "21", "24\nT5", "27"])
        ax.set_xlabel("Weeks postpartum | Time point", weight="semibold", labelpad=8)
    else:
        ax.set_xticklabels([])
        ax.set_xlabel("")
    if ax.get_subplotspec().is_first_col():
        ax.set_ylabel("Spatial colocalization $[Z(rho)]$", weight="semibold")
    else:
        ax.set_ylabel("")
        
fig.savefig(join(wd, "plots", "fig3_spatial_correlations.pdf"), bbox_inches="tight", dpi=300)
fig.savefig(join(wd, "plots", "fig3_spatial_correlations.png"), bbox_inches="tight", dpi=300)

# %% [markdown]
### Receptor cross correlation

# %% [code]
temp_res = pd.read_csv(join(wd, "results", "spc_receptors.csv"), index_col=0, header=[0,1])
fig, axes = plt.subplots(len(pred_aba_pet.columns)-1, len(pred_aba_pet.columns)-1, figsize=(19,18), 
                         sharey="row", sharex="col")

for r, receptor_r in enumerate(pred_aba_pet.columns[1:]):
    for c, receptor_c in enumerate(pred_aba_pet.columns[:-1]):
        ax = axes[r,c]
        if r < c:
            ax.set_axis_off()
        else:        
            rho = temp_res.loc[receptor_r, (receptor_c, "rho")]
            p = (temp_res.loc[receptor_r, (receptor_c, "p")] + temp_res.loc[receptor_c, (receptor_r, "p")]) / 2
            sns.regplot(
                y=minmax_scale(pred_aba_pet[receptor_r]),
                x=minmax_scale(pred_aba_pet[receptor_c]),
                ax=ax,
                scatter_kws=dict(alpha=0.5),
                color="tab:red" if p < .05 else "tab:blue"
            )
            ax.annotate(
                f"$Z(rho) = {rho:.02f}$"
                f"\n$p(exact) = {p:.04f}$",
                xy=(0.05,0.05),
                xycoords="axes fraction",
                ha="left", va="bottom",
                size=12
            )
            if ax.get_subplotspec().is_last_row():
                ax.set_xlabel(f"{receptor_c}\n$[Normalized\ density]$", weight="semibold", size=12)
            else:
                ax.set_xlabel("")
            if ax.get_subplotspec().is_first_col():
                ax.set_ylabel(f"{receptor_r}\n$[Normalized\ density]$", weight="semibold", size=12)
            else:
                ax.set_ylabel("")

fig.tight_layout()
fig.savefig(join(wd, "plots", "figS6_receptor_spc.pdf"), bbox_inches="tight")
fig.savefig(join(wd, "plots", "figS6_receptor_spc.png"), bbox_inches="tight", dpi=300)

# %% [markdown]
### Spatial association scatters

# %% [code]
temp = spc_cross[spc_cross.loc[:, ("T0", "q(norm)")] < 0.05]
measures = temp.index.get_level_values("Measure").unique()

fig = plt.figure(figsize=(19,33))
gs = GridSpec(temp.shape[0] + len(measures) + len(measures)-1, len(tps), hspace=0.2, height_ratios=[1]*3 + [0.2, 1, 1, 0.2] + [1]*6)

## BRAINS
parcellater = NiftiLabelsMasker(parc, parc_labels).fit()
temp_mri = pd.DataFrame(
    {m: parcellater.transform(sec_level_res[m.lower()]["con1"]["map_par_effectsize"]).squeeze() for m in measures},
    index=parc_labels
)
for r, m in zip([0, 4, 7], measures):
    ax = fig.add_subplot(gs[r,:])
    plot_stat_map(
        parcel_data_to_volume(temp_mri[m], parc)[0],
        cut_coords=np.arange(-30,71,10),
        display_mode="z",
        cmap="icefire",
        vmax=temp_mri[m].abs().max(),
        symmetric_cbar=True,
        axes=ax,
        title=m+" (baseline comparison: beta)",
        annotate=False,
    )
    
## DISTANCE PLACEHOLDER
for r in [3,6]:
    ax = fig.add_subplot(gs[r,:])
    ax.set_axis_off()

## SCATTERS
for r, (m, receptor) in zip([1,2, 5, 8,9,10,11,12], temp.index):
    
    temp_z = pd.read_csv(join(wd, "results", f"spc_pp_data_{m}.csv"), index_col=[0,1,2])
    temp_z = (temp_z.loc[(subs_PP, na(), "PP"), :] - temp_z.loc[(subs_NP, na(), "NP"), :].mean()) / temp_z.loc[(subs_NP, na(), "NP"), :].std()
    temp_z = temp_z.groupby("idx_tp").mean().T
    
    for c, tp in enumerate(tps):
        ax = fig.add_subplot(gs[r,c])
        
        sns.regplot(
            x=temp_z[tp],
            y=pred_aba_pet[receptor],
            scatter=False,
            ax=ax
        ) 
        sns.scatterplot(
            x=temp_z[tp],
            y=pred_aba_pet[receptor],
            hue=temp_mri[m],
            legend=False,
            palette="icefire",
            hue_norm=(-temp_mri[m].abs().max(), temp_mri[m].abs().max()),
            edgecolor="k",
            alpha=0.6,
            ax=ax
        )
        ax.annotate(
            f"$rho = {corr(pred_aba_pet[receptor], temp_z[tp], rank=True):.02f}$",
            xy=(0.05,0.95), 
            xycoords="axes fraction", 
            ha="left", va="top",
            size=12,
            bbox=dict(boxstyle="round", fc="w", alpha=0.7)
        )
        
        if r in [2, 5, 12]:
            ax.set_xlabel(f"Mean {m} $[Z\ vs.\ NP]$")
        else:
            ax.set_xlabel("")
        
        if c==0:
            ax.set_ylabel(f"{receptor} $[normalized\ density]$")
            ax.annotate(
                f"{m}-{receptor}\ncolocalization", 
                xy=(-0.4,0.5), 
                xycoords="axes fraction", 
                rotation=90,
                size=15, weight="semibold", ha="center", va="center"
            )
        else:
            ax.set_ylabel("")
            
        if r in [1, 5, 8]:
            ax.set_title(tp, weight="semibold", size=14, pad=9)
        else:
            ax.set_title("")

fig.savefig(join(wd, "plots", "figS5_spatial_correlation_scatters.pdf"), bbox_inches="tight", dpi=300)
fig.savefig(join(wd, "plots", "figS5_spatial_correlation_scatters.png"), bbox_inches="tight", dpi=300)

# %% [markdown]
### Longitudinal associations

# %% [code]
fig = plt.figure(figsize=(18, 16.5))
gs = GridSpec(4, 4, figure=fig,
              width_ratios=(1.2,1,1,1),
              hspace=1.5, wspace=0.05)


## HEATMAPS
y_labs = assoc_behav[("Interaction", "coef")].index.get_level_values("X").unique()
y_labs = [c for c in y_labs if "fALFF" in c] + [c for c in y_labs if "LCOR" in c] + [c for c in y_labs if "GCOR" in c]
lim = np.max(np.abs(assoc_behav[("Interaction", "coef")].to_list() + assoc_horm[("Interaction", "coef")].to_list()))

for i, (title, ax, x_labs, coef, p, q) in enumerate(zip(
    ["Hormones", "Behavior"],
    [fig.add_subplot(gs[:2,0]), fig.add_subplot(gs[2:4,0])],
    [['log(Estradiol)', 'log(Progesterone)', 'log(Estradiol/Progesterone)'],
     ['EPDS', 'MPAS - total', 'MPAS - attachment', 'MPAS - hostility', 'MPAS - interaction']],
    [assoc_horm[("Interaction", "coef")].reset_index().pivot(columns="X", index="Y"),
     assoc_behav[("Interaction", "coef")].reset_index().pivot(columns="Y", index="X")],
    [assoc_horm[("Interaction", "p")].reset_index().pivot(columns="X", index="Y"),
     assoc_behav[("Interaction", "p")].reset_index().pivot(columns="Y", index="X")],
    [assoc_horm[("Interaction", "q")].reset_index().pivot(columns="X", index="Y"),
     assoc_behav[("Interaction", "q")].reset_index().pivot(columns="Y", index="X")]
)):

    annot = p.copy().astype(str)
    annot[p <= 0.05] = "☆"
    annot[p > 0.05] = ""
    annot[q <= 0.05] = "★"
    
    ax_cbar = ax.inset_axes((0,1.1,1,0.03))
    sns.heatmap(
        data=coef.loc[y_labs, (na(), na(), x_labs)],
        cmap="RdBu_r",
        vmin=-lim,
        vmax=lim,
        center=0,
        ax=ax,
        cbar=True, 
        cbar_ax=ax_cbar,
        cbar_kws=dict(label=f"Weeks PP × {'MRI' if title=='Hormones' else title} interaction effect", 
                      orientation="horizontal"),
        square=True,
        linewidths=1,
        linecolor="k",
        annot=annot.loc[y_labs, (na(), na(), x_labs)],
        fmt="s",
        annot_kws=dict(size=20, ha="center", va="center", fontweight="bold")
    )
    ax_cbar.set_xticks([-0.03, 0, 0.03])
    
    ax_cbar.spines["outline"].set(visible=True, lw=.8, edgecolor="black")
    sns.despine(top=False, right=False, bottom=False, left=False)
    ax.set_title(title, weight="semibold", size=14, pad=10)
    ax.set_ylabel("")
    ax.set_xlabel("")
    ax.set_xticklabels(x_labs, rotation=-35, ha="left", rotation_mode="anchor")

    ax_lab = ax.inset_axes((-2.2 if title=="Hormones" else -1.35, 0,0.1,1))
    ax_lab.set_ylim(ax.get_ylim())
    ax_lab.set_axis_off()
    n_falff = len([c for c in y_labs if "fALFF" in c])
    n_lcor = len([c for c in y_labs if "LCOR" in c])
    n_gcor = len([c for c in y_labs if "GCOR" in c])
    for r, m in zip([(0.5, n_falff-0.5), 
                    (n_falff+0.5, n_falff+n_lcor-0.5), 
                    (n_falff+n_lcor+0.5, n_falff+n_lcor+n_gcor+-0.5)], 
                    metrics):
        ax_lab.plot([0,0], [r[0]-0.2, r[1]+0.2], c="k")
        ax_lab.text(s=m, x=-0.1, y=np.mean(r), rotation=90, va="center", ha="center", size=12, weight="semibold")
        
        
## DEVELOPMENT
gs_horm = GridSpecFromSubplotSpec(2, 3, subplot_spec=gs[:2,1:], wspace=0.25)
gs_behav = GridSpecFromSubplotSpec(2, 3, subplot_spec=gs[2:4,1:], wspace=0.25)

for r, (y_labs, y_vars, axes, sig_locs) in enumerate(zip(
    [['Estradiol', 'Progesterone', 'Estradiol/Progesterone'],
     ['EPDS', 'MPAS - total', 'MPAS - attachment', 'MPAS - hostility', 'MPAS - interaction']],
    [["estradiol_log", "progesterone_log", "ep_ratio_log"], 
     ["epds",  "mpas", "mpas_attachment", "mpas_hostility", "mpas_interaction"]],
    [[fig.add_subplot(loc) for loc in [gs_horm[0,0], gs_horm[0,1], gs_horm[1,0]]],
     [fig.add_subplot(loc) for loc in [gs_behav[0,0], gs_behav[0,1], gs_behav[1,0], gs_behav[1,1], gs_behav[1,2]]]],
    [["top", "top", "bottom"],
     ["top", "bottom", "bottom", "bottom", "bottom"]]
)):
    
    for c, (y, y_lab, ax, sig_loc) in enumerate(zip(y_vars, y_labs, axes, sig_locs)):
        
        temp = data_spc.loc[(subs_PP_long, na(), "PP"), [y, "weeks_postpartal", "ID"]].dropna()

        sns.scatterplot(
            temp,
            x="weeks_postpartal",
            hue="ID",
            y=y,
            ax=ax,
            legend=None,
            alpha=0.4,
            zorder=3,
            palette=colors
        )
        sns.lineplot(
            temp,
            x="weeks_postpartal",
            y=y,
            hue="ID",
            ax=ax,
            legend=None,
            alpha=0.4,
            zorder=2,
            palette=colors
        )
        temp_mean = temp.groupby("idx_tp").mean()
        temp_max = temp.groupby("idx_tp").max()
        for tp in temp.index.get_level_values("idx_tp").unique():   
            ax.boxplot(
                x=temp.loc[(subs_PP_long, tp, "PP"), y],
                positions=[temp_mean.loc[tp, "weeks_postpartal"]],
                widths=2,
                zorder=10,
                showfliers=False,
                medianprops=dict(c="k")
            )
            ax.scatter(
                y=temp_mean.loc[tp, y],
                x=temp_mean.loc[tp, "weeks_postpartal"],
                zorder=11,
                c="k",
                s=30
            )     
        
        # LMM
        for var in ["Weeks PP", "Weeks PP^2"]:
            p = dem_data_long.loc[y_lab if r==1 else f"log({y_lab})", (var, "p")]
            if p < 0.05: s = "☆"
            else: s = "ns"
            ax.annotate(
                f"{'Linear' if var=='Weeks PP' else 'Quadratic'} effect: {s}",
                xy=(0.45, (0.05 if sig_loc == "bottom" else 0.83) + (0.08 if var=='Weeks PP' else 0)),
                xycoords="axes fraction",
                size=11
            )
                
        ax.set_title(y_lab, size=12, fontweight="semibold")
        ax.set_xticks(list(range(0,28,3)))
        if any(s in y_lab.lower() for s in ["mpas", "epds"]):
            ax.set_ylabel("(Sub)scale score")
        else:
            ax.set_ylabel("Hormone level $[log]$")

        if ax.get_subplotspec().is_last_row():
            ax.set_xticklabels(["0\nT0", "3\nT1", "6\nT2", "9\nT3", "12\nT4", "15", "18", "21", "24\nT5", "27"])
            ax.set_xlabel("Weeks postpartum | Time point", weight="semibold", labelpad=8)
        else:
            ax.set_xticklabels([])
            ax.set_xlabel("")


fig.savefig(join(wd, "plots", "fig4_hormone_behavior.pdf"), bbox_inches="tight")
fig.savefig(join(wd, "plots", "fig4_hormone_behavior.png"), bbox_inches="tight", dpi=300)

# %% [markdown]
### RTM sensitivity

# %% [code]
fig = plt.figure(figsize=(18,8))
gs = GridSpec(2, 3, figure=fig, 
              width_ratios=(1,1,1), height_ratios=(0.8,1),
              hspace=0.3, wspace=0.2)

# GLM brains
ax_brain = fig.add_subplot(gs[0,:])
ax_brain.set_axis_off()
ax_brain_con1 = ax_brain.inset_axes((0,0,0.48,1))
ax_brain_con2 = ax_brain.inset_axes((0.52,0,0.48,1))

vmin = 0
vmax = 6

for con, ax in zip(["con1", "con2"], [ax_brain_con1, ax_brain_con2]):
    
    map_unthresh = sec_level_res_rtm["gcor"][con]["map_par_logp"]
    map_thresh = sec_level_res_rtm["gcor"][con]["map_nonpar_logp"]
        
    # plot main glass brain
    gb = plot_glass_brain(
        map_unthresh, 
        display_mode="lzry",
        alpha=0.4,
        colorbar=False, 
        cmap="cold_white_hot" if con=="con1" else "cold_white_hot_r",
        figure=fig,
        axes=ax,
        vmin=vmin,
        vmax=vmax
    )
        
    # significant cluster
    if (map_thresh.get_fdata() > cl_thresh).any():
        
        # plot overlays
        gb.add_contours(
            img=threshold_img(map_thresh, cl_thresh), 
            levels=[1], 
            colors="k", 
            linewidth=5,
            filled=True,
            alpha=0.5
        )
    
    # titles
    ax.set_title(f"T0 (longitudinal excluded): PP {'>' if con=='con1' else '<'} NP $(n = {data.loc[(subs_rtm,'T0','PP'),:].shape[0]}\ |\ "
                    f"{data.loc[(subs_rtm,'T0','NP'),:].shape[0]})$", 
                    size=16, weight="semibold", pad=10)
    if con=="con1": 
        ax.annotate(
            "GCOR", 
            xy=(-0.04,0.5), 
            xycoords="axes fraction", 
            rotation=90,
            size=16, weight="semibold", ha="center", va="center"
        )
    
    # colorbar
    norm = mpl.colors.Normalize(vmin=vmin, vmax=vmax)
    sm = plt.cm.ScalarMappable(
        cmap="black_red_r" if con=="con1" else "black_blue_r", 
        norm=norm)
    sm.set_array([])
    plt.colorbar(
        sm,
        label=r"$-lg(p)$",
        orientation="horizontal",
        cax=ax.inset_axes((0,0,1,0.07)),
    )
    

# Cluster development

ax_clust1 = fig.add_subplot(gs[1,0])
ax_clust2 = fig.add_subplot(gs[1,1])

for (lab, measure, contrast, name), ax in zip(cl_iter_rtm, [ax_clust1, ax_clust2]):
    
    temp_mri = pd.concat(
        [data.loc[(subs_NP, ), :].melt(["ID", "group_str", "weeks_postpartal"], lab)] + \
        [data.loc[(subs_PP_long, tp), :].melt(["ID", "group_str", "weeks_postpartal"], lab).replace({"PP": f"PP-{tp}"}) for tp in ["T1", "T2", "T3", "T4", "T5"]]
    )
    temp_mri.loc[temp_mri.group_str=="NP", "weeks_postpartal"] = -3.5

    sns.scatterplot(
        temp_mri,
        x="weeks_postpartal",
        hue="ID",
        y="value",
        ax=ax,
        legend=None,
        alpha=0.4,
        zorder=3,
        palette=colors
    )
    sns.lineplot(
        temp_mri,
        x="weeks_postpartal",
        y="value",
        hue="ID",
        ax=ax,
        legend=None,
        alpha=0.4,
        zorder=2,
        palette=colors
    )
    
    for group in temp_mri.group_str.unique():   

        ax.boxplot(
            x=temp_mri.loc[temp_mri.group_str==group, "value"],
            positions=[temp_mri.loc[temp_mri.group_str==group, "weeks_postpartal"].mean()],
            widths=2,
            zorder=10,
            showfliers=False,
            medianprops=dict(c="k")       
        )
        ax.scatter(
            y=[temp_mri.loc[temp_mri.group_str==group, "value"].mean()],
            x=[temp_mri.loc[temp_mri.group_str==group, "weeks_postpartal"].mean()],
            zorder=11,
            c="k",
            s=30
        )
    
    # LMM
    for var in ["Weeks PP", "Weeks PP^2"]:
        p = mri_long_lmm_rtm.loc[(measure, contrast, name), (var, "-lg(p)")]
        if p > -np.log10(0.05): s = "☆"
        else: s = "ns"
        ax.annotate(
            f"{'Linear' if var=='Weeks PP' else 'Quadratic'} effect: {s}",
            xy=(0.6, 0.05 + (0.07 if var=='Weeks PP' else 0)),
            xycoords="axes fraction",
            size=11
        )
    
    # Labels
    ax.set_title(f"{measure} - {contrast}: {name} {'cluster' if measure!='NBS' else 'average connectivity'}", weight="semibold")
    ax.set_ylabel(f"Average {measure if measure!='NBS' else 'network connectivity'} $[Z]$", weight="semibold")
    ax.set_xticks([-3.5] + list(range(0,28,3)))
    ax.set_xticklabels(["NP", "0\nT0", "3\nT1", "6\nT2", "9\nT3", "12\nT4", "15", "18", "21", "24\nT5", "27"])
    ax.set_xlabel("Group | Weeks postpartal | Timepoint", weight="semibold", labelpad=8)
    

# SPC at baseline

ax_spc = fig.add_subplot(gs[1,2])
temp = spc_zr_rtm.loc[(na(), "T0", "PP"),:].reset_index().melt(id_vars="idx_metric", value_vars=spc_zr_rtm.columns)

sns.stripplot(
    temp,
    x="variable",
    hue="value",
    y="value",
    dodge=False,
    ax=ax_spc,
    palette="icefire",
    edgecolor="k",
    linewidth=0.5,
    jitter=0.3,
    legend=None,
    alpha=0.4,
    zorder=1
)
sns.boxplot(
    temp,
    x="variable",
    #hue="variable",
    y="value",
    ax=ax_spc,
    fliersize=0,
    width=0.5,
    boxprops=dict(facecolor="none"),
    zorder=2,
    linewidth=1,
)
receptors = [l.get_text() for l in ax_spc.get_xticklabels()]
ax_spc.scatter(
    x=receptors,
    y=temp[temp.idx_metric=="gcor"].groupby("variable").mean().loc[receptors],
    zorder=20,
    c=["orange" if r in pred_pet.columns else "lightgreen" for r in receptors],
    s=30,
    ec="k"
) 

for x, r in enumerate(spc_zr_rtm.columns):
    
    if spc_sig_rtm[r].values[0] < 0.05: s = "☆"
    else: s = ""
    ax_spc.text(
        s=s, 
        x=x, 
        y=spc_zr_rtm.loc[(na(), "T0", "PP", "gcor"), r].max() + 0.08, 
        ha="center", va="center", size=12
    )  
        
ax_spc.axhline(c="k", zorder=0, lw=1)
ax_spc.set_xticks(ax_spc.get_xticks())
ax_spc.set_xticklabels(ax_spc.get_xticklabels(), rotation=-30, rotation_mode="anchor", ha="left")
ax_spc.set_ylim(-0.85,0.85)
ax_spc.set_xlabel("Receptor maps", weight="semibold", size=11)
ax_spc.set_ylabel("Spatial colocalization $[Z(rho)]$", weight="semibold")
ax_spc.set_title("Baseline colocalization with GCOR", weight="semibold")

fig.savefig(join(wd, "plots", "figS2_rtm_sensitivity.pdf"), bbox_inches="tight")
fig.savefig(join(wd, "plots", "figS2_rtm_sensitivity.pdf"), bbox_inches="tight", dpi=400)

# %% [markdown]
## Save notebook to HTML

# %% [code]
!!jupyter nbconvert 3_results_analyses_plots.ipynb --to html

# %% [code]
```
